# From flexible to anticipatory processing: alpha and beta oscillatory signatures of feedback-guided strategy adaptation and memory updating

**DOI:** 10.64898/2026.05.10.724182

**Authors:** Maya Al Safadi, Alex Chatburn, Zachariah Cross, Shane Dawson, Ina Bornkessel-Schlesewsky

## Abstract

When humans learn under conditions of uncertainty, they dynamically adjust how they prepare for and respond to feedback. In navigating uncertain environments, the brain minimizes error by continuously refining internal models via memory updating (MU). Feedback is critical for MU, and anticipatory neural mechanisms shape how feedback is processed, likely reflecting learned environmental certainty. However, the literature has largely focused on post-feedback activity, leaving pre-feedback certainty-related mechanisms less understood. The present study aims to address this gap by examining how certainty modulates anticipatory states, preceding feedback and subsequent MU. We examined oscillatory activity prior to performance feedback in a reanalysis of EEG data previously published by Hassall and colleagues (2023). Twenty-one participants (16 female, *M*_age_ = 25.81 years) predicted the strength of cartoon characters with varying predictability levels which were learned through exposure. Feedback on prediction accuracy was presented via an animated rising bar. Results revealed that theta power is modulated by accumulative feedback. Linear mixed-effects models revealed an interaction between predictability-related certainty and learning stage: in late learning, higher performance was associated with increased pre-feedback alpha and beta power for low-certainty trials, whereas in early learning, higher performance was associated with decreased beta power. These learning-related modulations in alpha and beta power suggest that initial learning is marked by adaptable exploratory processing. Subsequent learning exhibited increased alpha-mediated inhibition and beta-related anticipatory activity for lower certainty trials, indicative of dynamic strategy refinement and selective engagement of task-relevant information. These results demonstrate that certainty shapes preparatory oscillatory activity associated with MU.

## 1. Introduction

The adaptation of predictive models via external feedback is crucial for learning and memory (Zheng et al., 2022). A predictive model comprises our expectations, based on prior knowledge and experiences stored in memory (Hoppe et al., 2022), which aid adaptation to the environment (Hoppe et al., 2022). Previous literature suggests that error feedback supports long-term memory formation through the processes of MU (Zheng et al., 2022). However, our understanding of the mechanisms involved in memory updating and the optimal conditions for memory updating remains incomplete. Additionally, previous research investigating the effect of error feedback on memory performance has typically focused on examining error-related neural activity post error feedback, with no consideration of neural activity preceding error feedback (Cavanagh et al., 2012). This is despite evidence that anticipatory (pre-feedback) neural activity can vary with learned environmental predictability, and/or inferred certainty, and that such anticipatory states can influence memory performance (Scholz et al., 2017; Ostrowski & Rose, 2024; Weismüller et al., 2019; Salari & Rose, 2016; Toosi et al., 2017; Baslar & Stampfer, 1985; Van der Borght et al., 2016; Butterfield & Metcalfe, 2006). The remainder of the introduction is structured as follows: First we begin by outlining Error-Driven Learning (EDL) theories and the previously established link between EDL and theta power. We then discuss the neural correlates often linked with memory performance and outline the need for investigating pre-feedback neural activity.

Finally, we discuss the link between memory certainty and the process of memory updating before we outline the aims and hypothesis of this current study. In the present study, “predictability refers to the probabilistic structure of the task, whereas “certainty” reflects the learner’s inferred belief strength arising from this structure.

### 1.1 Error feedback and memory updating

While there is a vast literature on how error feedback impacts memory performance and learning, there is limited knowledge on the underlying mechanisms supporting this interaction (Cavanagh et al., 2009; 2012; Hoppe et al., 2022; Zheng et al., 2022; Wahlheim et al., 2022). Prior studies demonstrate that error feedback significantly enhances memory performance in comparison to no feedback, and this has been attributed to the additional opportunity to encode the correct answer and thereby correct originally incorrect retrievals (Pfabigan et al., 2011; Carneiro et al., 2021; Hoffrage et al., 2000; Kornell et al., 2009; Butler et al., 2008; Finn & Metcalfe, 2010; Guran et al., 2020; Hebsher et al., 2019; Cavanagh et al., 2009; 2012). These findings support EDL theories, which suggest that performance feedback precipitates memory updating by weakening memory co-activations that result in errors and strengthening ones that result in correct outcomes (Hoppe et al., 2022; Kopp & Wolff, 2000). This leads to the modification and further stabilization of new correct memories (Hoppe et al., 2022; Zheng et al., 2022; Wahlheim et al., 2022). Theta power (∼3-8Hz) is commonly investigated in the context of error feedback, and it has been suggested that an increase in frontal and midfrontal theta power post error feedback indexes memory updating and results from EDL mechanisms such as the strengthening and weakening of co-activations following an error (Bush et al., 2002; Cavanagh et al., 2009, 2012; Luu et al., 2003; van de Vijver et al., 2011; Wang et al., 2017). This is supported by various studies that reported increased theta power for items following feedback-based error correction compared with items for which errors remained uncorrected (Cavanagh et al., 2009, 2012; van de Vijver et al., 2011) and by studies demonstrating that theta power increases following error feedback and decreases following corrective feedback (Bush et al., 2002; Luu et al., 2003; Wang et al., 2017). Prior studies suggest that theta power is modulated by errors and posit that the increase in theta power following error feedback reflects surprise and the need to adapt behavior and update information (Cavanagh et al., 2009, 2012; van de Vijver et al., 2011). It has been previously postulated that theta power reflects communication between the medial prefrontal cortex (mPFC) and the lateral prefrontal cortex (lPFC) as a result of an error, signaling the need to adjust behavior to optimize performance (Cavanagh et al., 2009).

### 1.2 The neural correlates of error feedback

When attempting to investigate the neural correlates of error feedback, studies often focus on observing electrophysiological events post error feedback (Bush et al., 2002; Cavanagh et al., 2009, 2012; Luu et al., 2003; van de Vijver et al., 2011; Wang et al., 2017). This limits our understanding of how the brain may anticipate and prepare for encoding errors and how this might impact the memory updating process. Studies demonstrate that *pre-stimulus* neural activity is linked to subsequent memory outcomes (Scholz et al., 2017; Ostrowski & Rose, 2024; Weismüller et al., 2019; Salari & Rose, 2016; Toosi et al., 2017; Baslar & Stampfer, 1985). For example, increases in pre-stimulus beta (∼13–30 Hz) power have been shown to predict better subsequent memory performance and are thought to reflect a preparatory, top-down mechanism which maintains a neural state that supports stimuli encoding, which involving attentional allocation and inhibitory control processes. (Scholz et al., 2017; Ostrowski & Rose, 2024; Salari & Rose, 2016). Similarly, an increase in pre-stimulus alpha (∼8-12Hz) power was also shown to be linked to better memory performance and was suggested to reflect an inhibition of already encoded stimuli, in favor of new information (Ostrowski & Rose, 2024; Toosi et al., 2017; Baslar & Stampfer, 1985). This alpha-mediated inhibition is proposed to create rhythmic windows of reduced neuronal excitability that gate information processing in irrelevant brain regions, thereby suppressing task-irrelevant input (Jensen & Mazaheri, 2010). Collectively, these mechanisms are thought to aid in stabilizing newly encoded information in long-term memory (Scholz et al., 2017). However, pre-stimulus correlates such as beta and alpha power have primarily been investigated in the context of memory encoding paradigms and have not been investigated in the context of error feedback. If pre-stimulus activity influences memory performance and is thought to reflect preparatory and inhibitory processes that influence memory, it is plausible to suggest that pre-feedback neural activity may influence the process of memory updating in a similar manner via error feedback and in turn predict memory performance.

### 1.3 Certainty and memory updating

Memory certainty reflects the graded subjective belief regarding the probability of various outcomes when making decisions (Kiani et al., 2014). Error feedback has been found to impact memory correction based on certainty levels (Butterfield & Metcalfe, 2001; 2006; Butterfield & Mangels, 2003; Huelser & Metcalfe, 2012; Kang et al., 2011; Sitzman, Rhodes, & Tauber, 2014; Metcalfe & Eich, 2019). For instance, feedback to errors committed with high certainty improved memory more substantially in comparison to errors committed with low certainty (Butterfield & Metcalfe, 2001; 2006; Butterfield & Mangels, 2003; Huelser & Metcalfe, 2012; Kang et al., 2011; Sitzman, Rhodes, & Tauber, 2014; Metcalfe & Eich, 2019). The enhancing effect of high certainty errors is attributed to these errors being highly surprising and thus capturing more attention than low confidence errors (Butterfield & Metcalfe, 2006), thereby leading to a better encoding of the correct answer (Van der Borght et al., 2016).

Several neural correlates of certainty in memory updating have been identified, such as the N100 and the P200 event-related potentials (ERP; Polezzi et al., 2008; Schuermann et al., 2012; Flor et al., 2002; Pizzagalli et al., 2003; Wills et al., 2007). The N100 and P200 were used to investigate certainty in terms of prediction errors and how the brain responds to error feedback, mostly from a classical conditioning and operant learning perspective (Polezzi et al., 2008; Schuermann et al., 2012; Flor et al., 2002; Pizzagalli et al., 2003; Wills et al., 2007). Results from these studies illustrate that a more pronounced N100 and P200 amplitude are associated with higher prediction errors and higher uncertainty (Polezzi et al., 2008; Schuermann et al., 2012; Flor et al., 2002; Pizzagalli et al., 2003; Wills et al., 2007). Previous research has also linked certainty to pre-stimulus alpha power, observing an increase in alpha power in certain experimental blocks compared to uncertain ones (Toosi et al., 2017). Pre-stimulus alpha power may index attentional inhibition driven by memory uncertainty prior to feedback, influencing subsequent feedback processing and memory performance. Overall, certainty may shape how error feedback is processed by modulating attentional, inhibitory, and preparatory mechanisms such as alpha and beta power, which in turn influence memory updating.

### 1.4 The present study

The current study aimed to further investigate the underlying mechanisms of memory updating. Prior literature has typically examined brain activity such as theta power following feedback without considering neural processes that occur prior to feedback and how these might affect the memory updating process (Cavanagh et al., 2009, 2012; van de Vijver et al., 2011; Luu et al., 2003; Wang et al., 2017). The current study investigated time-frequency neural activity (alpha and beta power) occurring prior to feedback, as pre-stimulus modulations of these neural correlates have been previously linked to memory outcomes (Ostrowski & Rose, 2024; Scholz et al., 2017; Salari & Rose 2016) in addition to feedback-related theta power. Moreover, this study aimed to explore the potential role of distinct predictability levels in the process of memory updating and the possible link between predictability (inferred certainty) levels and pre-feedback time frequency activity. To this end, we analyzed oscillatory neural activity occurring at two different time points, namely pre- and during-error feedback in data previously published by Hassall and colleagues (2023). In the original study, participants guessed the “strength” of six cartoon gnomes (i.e., cues) on a vertical bar, each associated with a different predictability level. The original study’s analysis investigated the neural correlates of continuous feedback via ERPs such as stimulus-preceding negativity (SPN). Our reanalysis examines oscillatory correlates of feedback processing and anticipatory states. In the current reanalysis, we hypothesized that feedback-related theta power would be higher for error feedback (i.e., lower performance points) in comparison to correct feedback (i.e., higher performance points) and that pre-feedback alpha and beta power would be modulated by predictability levels.

## 2. Methods

### 2.1 Original study

The present study was a reanalysis of Hassall and colleagues (2023), data set involving 21 participants (16 female, 2 left-handed, mean age = 25.81). All participants had normal or corrected-to-normal vision. Hassall et al (2023) aimed to investigate if continuous feedback processing can be accounted for by the discrepancy between expected and actual reward received (Colombo, 2014). This was investigated through associating different cues with different reward outcome probabilities (see Figure 1 for details on the original study’s probability distribution). In this study, feedback was presented as an animated bar which increased until it reached the correct outcome, to track how the brain processed continuous rather than discrete feedback. Participants were asked to guess the strength of cartoon gnome characters (i.e., cue) through selecting a location of a vertical bar outline: the higher the selection, the stronger the gnome. Subsequently, participants received feedback on their performance via the animated rising bar. There were six gnomes in total which participants encountered 25 times each for a total of 150 trials (see Figure 2 for the original study’s paradigm). To measure behavioral performance, points in each trial were calculated as the difference between participants guess and the actual outcome (in pixels) which was then converted to a value from 1-100 (higher points reflected closer guess to the actual outcome). Hassall et al (2023) recorded participants’ EEG using a 32-channel EEG system (EASYCAP GmbH) with a standard 10-20 montage. The original study’s EEG recording was sampled at 1000Hz and amplified (actiCHamp, Plus, Brain Products GmbH; Hassall et al., 2023).

**Figure 1.**
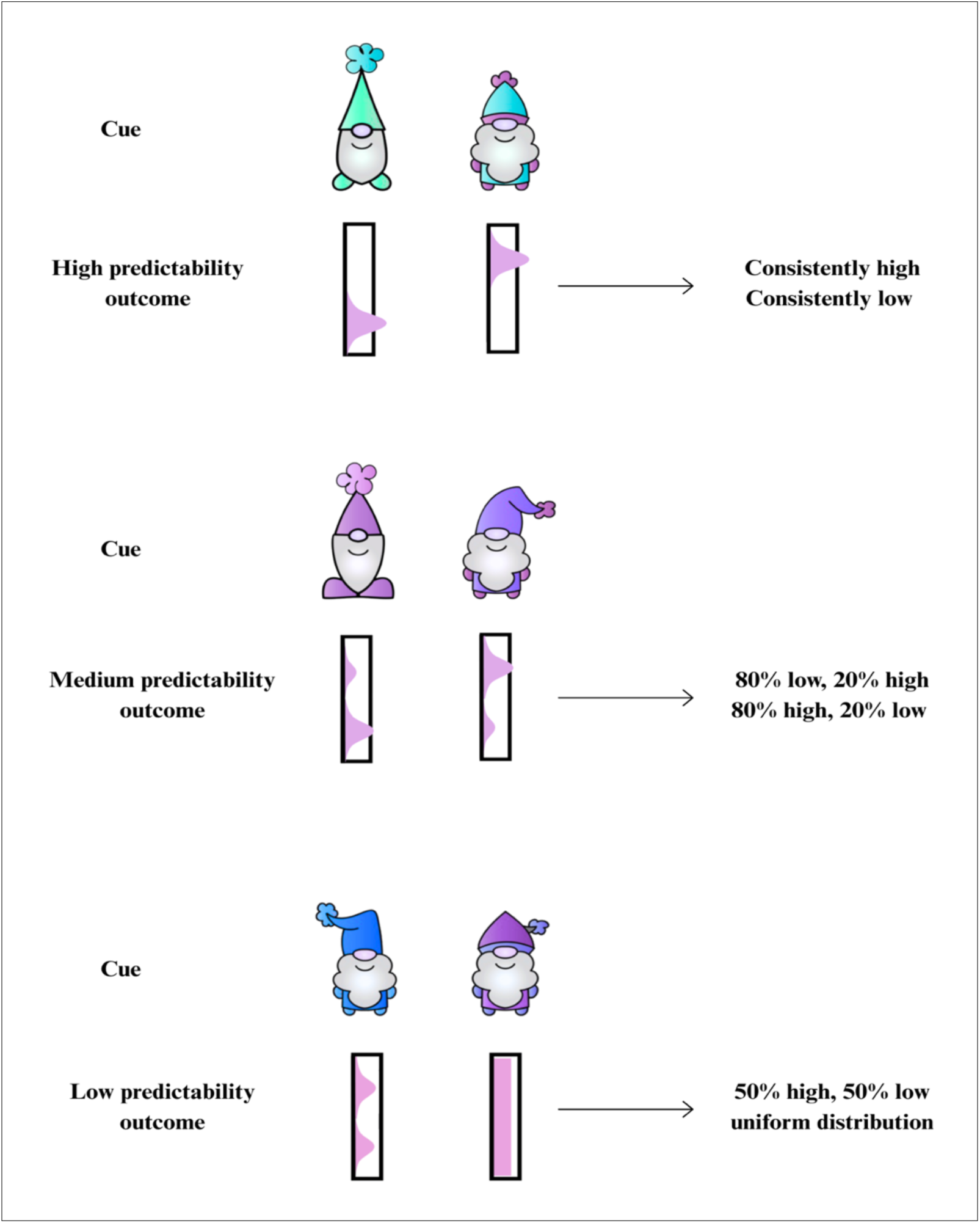
Reward outcome probability distributions associated with different cues in the original study. Each predictability condition incorporates two reward prediction likelihoods from the original study. Figure adapted from Hassall et al (2023).

**Figure 2.**
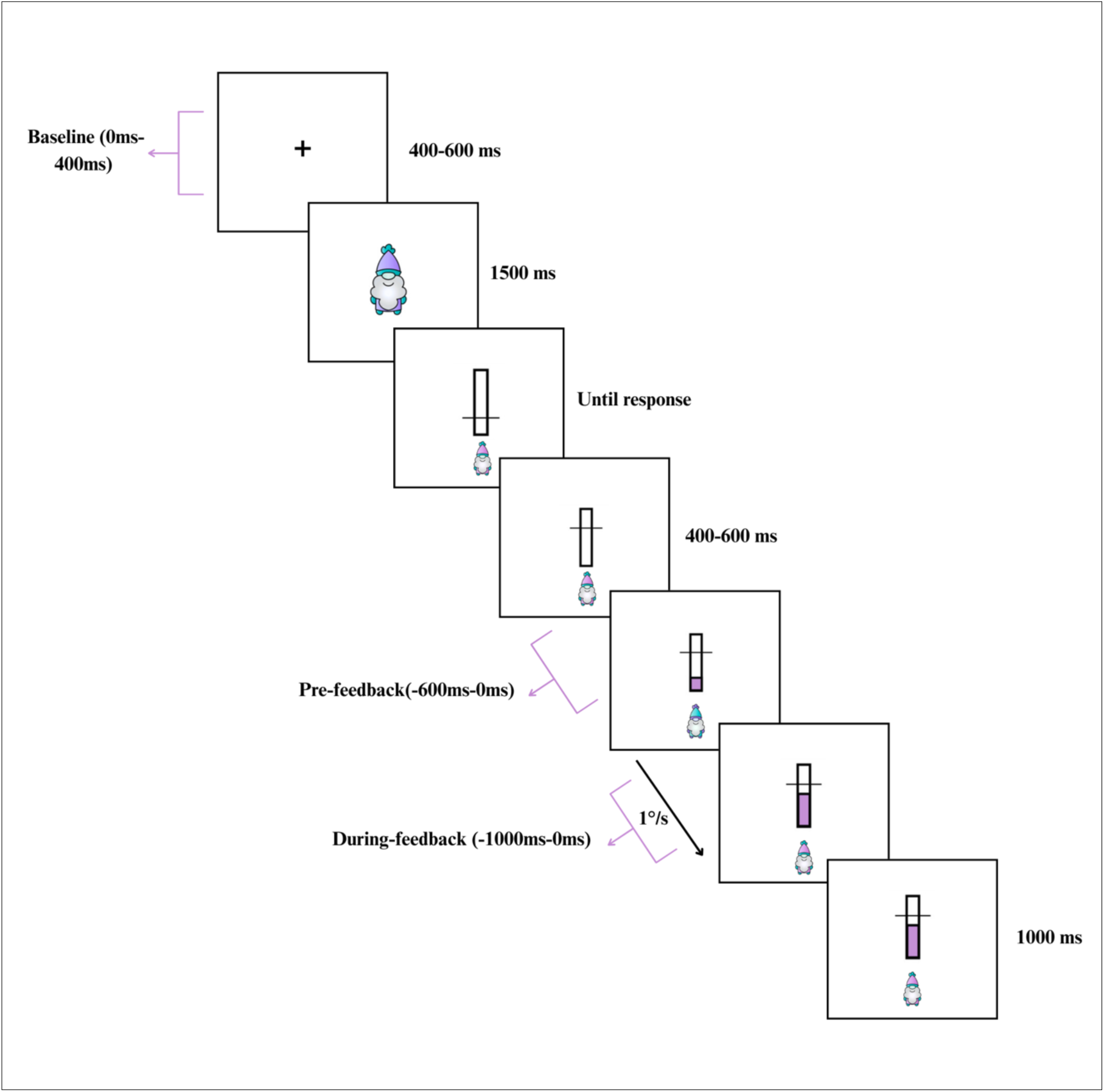
Hassall and colleagues (2023) original study paradigm. The purple-colored animated bar rose 1 degree per second until it reaches the outcome. The outcome subsequently remained on the screen for 1000 msec. The purple arrows represent epoched activity used for analysis. Figure adapted from Hassall et al (2023).

### 2.2 EEG pre-processing

EEG data were pre-processed using the MNE toolbox in Python version 3.10.17 (Gramfort et al., 2013). Data were re-referenced to TP9 and TP10. Non-overlapping, 3-second epochs were first generated across the continuous raw EEG data for each participant. Rejection thresholds were then computed from the epochs using *Autoreject* (Jas, Engemann, Raimondo, Bekhti, & Gramfort, 2016) package in MNE-Python, for inclusion in a subsequent ICA analysis for the detection and removal of artefacts. The ICA was subsequently fit to a 1Hz high pass and a 100Hz low pass filter. Independent components (ICs) were classified using the *ICLabel* (Pion-Tonachini et al., 2019) and they were categorized where components corresponding to eye movement, muscle movement and channel noise were removed from the raw EEG data. Components classified as “brain” or “other” (i.e., residual artifacts, mixed noise or unclear signal) were then filtered using a bandpass filter ranging from 0.1 to 30 Hz to eliminate low frequency drifts and high frequency noise. Raw data were epoched around events of interest as follows: from -1000ms to 1000ms around the pre-trial fixation cross to calculate baseline EEG activity, -1000ms to 1000ms relative to the beginning of the bar rising to capture pre-feedback brain activity and -1500ms and 500ms relative to the completion of the bar rising to capture feedback-related brain activity. These epochs allowed for the extraction of time frequency data and padding to avoid edge artifacts.

Feedback duration across all trials was calculated and all trials that included a bar rising period of less than 1 second were dropped from further analysis. This allowed for capturing multiple cycles of low frequency oscillations, resulting in better frequency resolution (Tallon-Bbaudry et al., 1999) around the events of interest. In total 48 trials were dropped. Subsequently, the *Autorejec*t function (Jas et al., 2016) was applied to remove or repair remaining bad epochs, and which were equal to or longer than 1 second across all participants. To examine broad topographic differences in the alpha, beta and theta frequency bands. EEG channels were assigned to 5 electrode clusters representing left anterior (i.e., F7, F3, FC1, FC5), right anterior (i.e., F8, F4, FC2, FC6), left posterior (i.e., P7, P3, CP5, CP1), right posterior (i.e., P8, P4, CP2, CP6) and midline regions (i.e., Pz, Oz, Cz, CPz, FCz). Mean power values were calculated as the average of activity at all channels within each pre-defined region of interest (ROI).

### 2.3 Time frequency analysis

Morlet wavelet analyses (Morlet et al., 1982) were implemented on artefact-free epochs using the *compute_tfr* function in MNE-python to estimate spectral activity in the alpha, beta and theta range. Frequency band limits were computed using the golden-mean approach (i.e., g), as described by Klimesch (2012) using a fixed peak alpha frequency (PAF) of 10 Hz. Frequency bands were unable to be adjusted based on participants’ individual alpha frequency (IAF) as there was no resting-state recording. The limits of each frequency band were defined by the algorithm is as follows; theta as 4.0-6.2Hz, alpha as 8.1-12.4Hz and beta as 16.2-24.7Hz.

Frequency bands were sampled linearly including 5 frequency bins per band. Oscillatory activity was obtained from smaller time windows within the longer epochs to avoid edge artefacts. Alpha and beta power were extracted from windows ranging from -600ms to 0ms relative to the beginning of the rising bar to capture time frequency brain activity pre-feedback (i.e., prior to participants learning the outcome of the trial). Theta power was extracted from windows ranging from -1000ms to 0ms relative to the end of the rising bar to capture feedback-related time frequency brain activity. Alpha, beta and theta power from 0ms to 400ms relative to the onset of the pre-trial fixation cross was also extracted across all trials to be included in subsequent statistical models as a covariate to control for baseline activity (Alday, 2019) and account for difference in frequency band power which occurred prior to events of interest.

### 2.4 Statistical analysis

Linear mixed-effects models (LMMs) were computed to examine the effects of interest while accounting for random effects using the *lme4* package (Bates et al., 2015; Bowsher, 2025) in R Version 4.4.3. The package *ggeffects* was used to compute adjusted predictions and marginal effects of LMMs outputs (Lüdecke, 2025). Data were visualized using the *ggplot2* function from the *tidyverse* collection of packages (Wickham et al., 2019). The following statistical models’ random effects structure was selected based on parsimonious selection procedure to avoid overparametrized models (Bates et al., 2018). The dependent variable in model 1 and 2 was performance points and fixed effects were time-frequency activity (i.e., pre-feedback alpha and pre-feedback beta), predictability level and topographical ROI. The dependent variable in model 3 was feedback-related theta power, and fixed effects were performance points, predictability conditions and topographical ROI. Covariates for model 1, 2 and 3 included a baseline measure of power by frequency band to account for differences in power prior to the occurrence of events of interest, and trial to account for changes in time-frequency activity across learning. To account for variability across cues (i.e., gnomes) and subjects, the random effects structure included by-subject random intercepts and slopes for condition, in addition to by-gnome random intercepts and slopes for pre-feedback alpha and beta. This complex random effect’s structure allows for variations in baseline performance across the different conditions, in addition to gnome-specific variability in the effects of alpha power (model 2), beta power (model 1) and performance points (model 3; Meteyard & Davies, 2020). The structure of statistical models for primary analyses are presented in Table 1. *p* values were calculated using the *lmerTest* package, based on Satterthwaite’s degrees of freedom (Kuznetsova et al., 2017). All predictor values were scaled and centered, and the alpha level was set at 0.05. Computations of models were optimized through using the Bound Optimization by Quadratic Approximation (BOBYQA) optimizer to increase robustness and stability of parameter estimation (Powell, 2010). To contrast categorical variables in the LMMs, the three predictability conditions were sum-to-zero contrast coded, such that coefficients reflect comparisons to the grand mean for all predictability conditions (Schad et al., 2022).

**Table 1.** LMM structure for primary analysis.

| *Model* | *Outcome* | *Fixed effects* | *Covariates* | *Random Intercepts* | *Random slopes* |
| --- | --- | --- | --- | --- | --- |
| 1 | Performance points | **Pre-feedback**<br>alpha power<br>feedback (-0.6s<br>to 0s)<br>Condition (low,<br>medium, high)<br>Roi | Baseline alpha<br>(0s to 0.4s)<br>Trial | + Subject<br>+ gnome | By-subject per<br>condition<br>By-gnome<br>according to pre-feedback alpha<br>power |
| 2 | Performance points | **Pre-feedback**<br>beta power (-<br>0.6s to 0s)<br>Condition (low,<br>medium, high)<br>Roi | Baseline beta<br>(0s to 4s)<br>Trial | + Subject<br>+ gnome | By-subject per<br>condition<br>By-gnome<br>according to pre-feedback beta<br>power |
| 3 | **Feedback-related**<br>theta power<br>(-1s to 0s) | Performance<br>points<br>Condition (low,<br>medium, high)<br>Roi | Baseline theta<br>(-1s to 0s)<br>Trial | + Subject<br>+ gnome | By-subject per<br>condition<br>By-gnome<br>according to<br>performance<br>points |

## 3. Results

### 3.1 Behavioral results

Performance across conditions is visualized in Figure 3. Overall, participants performed above chance in the “gnomes” task with an average of 67.7 out of 100 performance points (standard deviation: 23.4, range: 11-100). These results illustrate substantial inter-individual variability in performance across conditions. See original study for more details on the behavioral results (Hassall et al., 2023).

**Figure 3.**
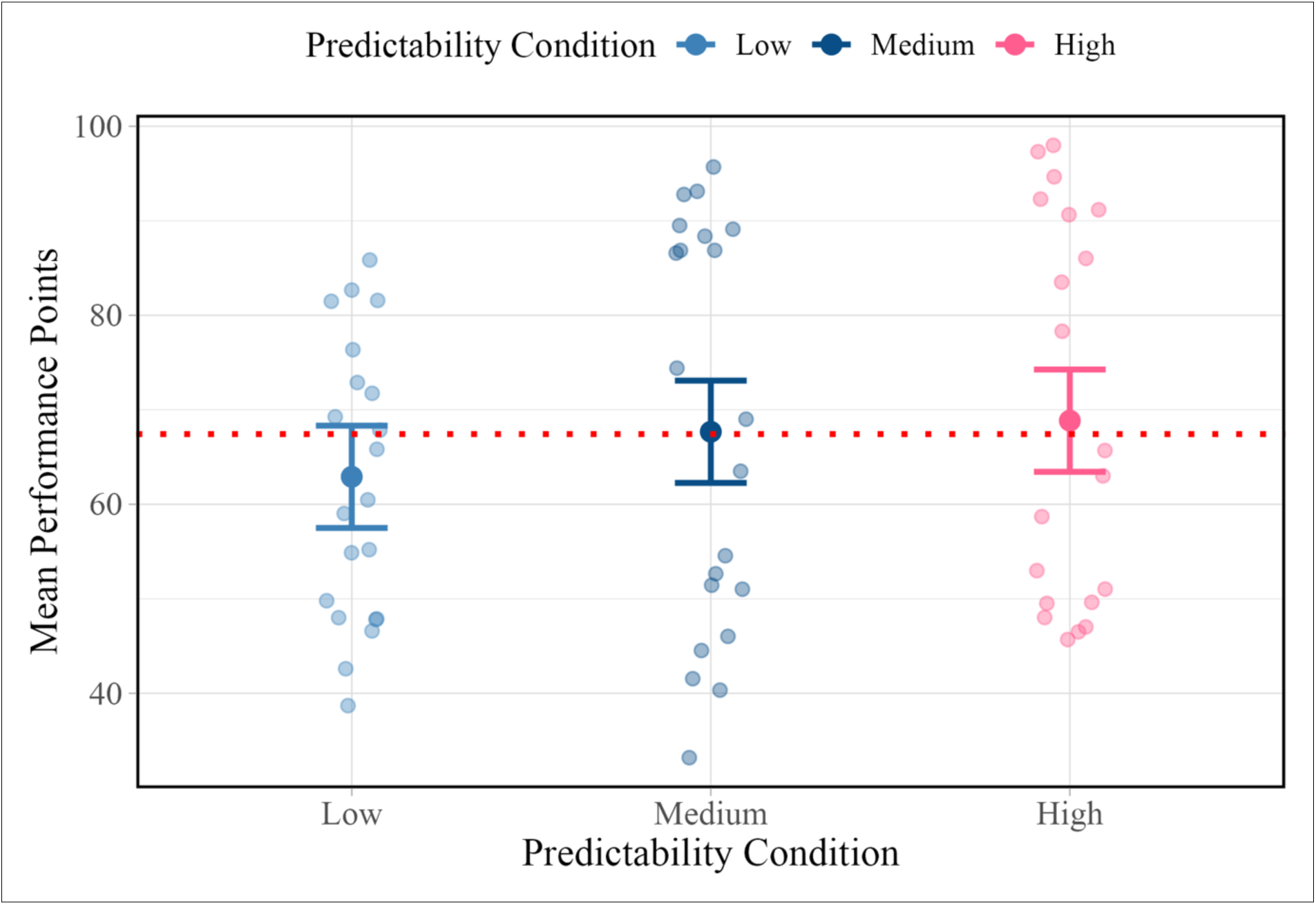
Average task performance in original study across the predictability conditions. Predictability level is present on the x-axis and mean performance on the y-axis with higher values reflecting better performance. The different colors represent the different predictability conditions, with light blue reflecting low predictability, dark blue reflecting medium predictability and pink reflecting high predictability.

### 3.2 Neurophysiological data

Topographical maps depicting raw pre-feedback alpha and beta activity across predictability conditions (low, medium, high) and learning stages (early, middle, late) are shown in Figure 4. Time-frequency representations of the pre-feedback epoch, showing power across a broad frequency range, are illustrated in Figure 5. Power spectral density (PSD) plots depicting pre-feedback (Appendix G) and feedback-related (Appendix H) activity across frequencies (1–30 Hz), separated by regions of interest and predictability conditions, the original spectra are shown alongside model fits, including the aperiodic component.

**Figure 4.**
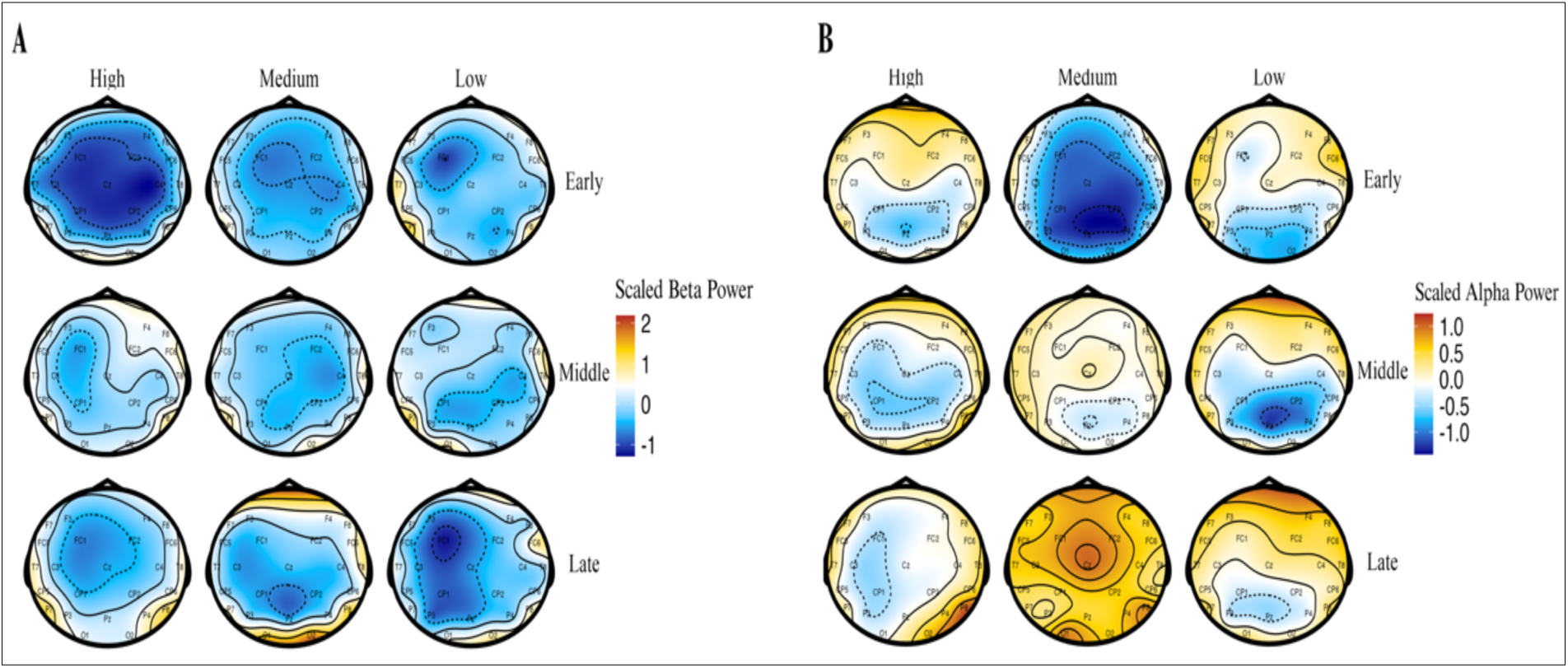
Topographical differences in pre-feedback alpha and beta between condition and learning stage. Topographical differences in pre-feedback beta power (A) and alpha power (B) faceted by different predictability conditions (i.e., high, medium and low) and learning stage (i.e., early, middle and late). Scaled power is presented on the z-axis, with warmer red shades reflecting higher power and cooler blue shades reflecting lower power.

**Figure 5.**
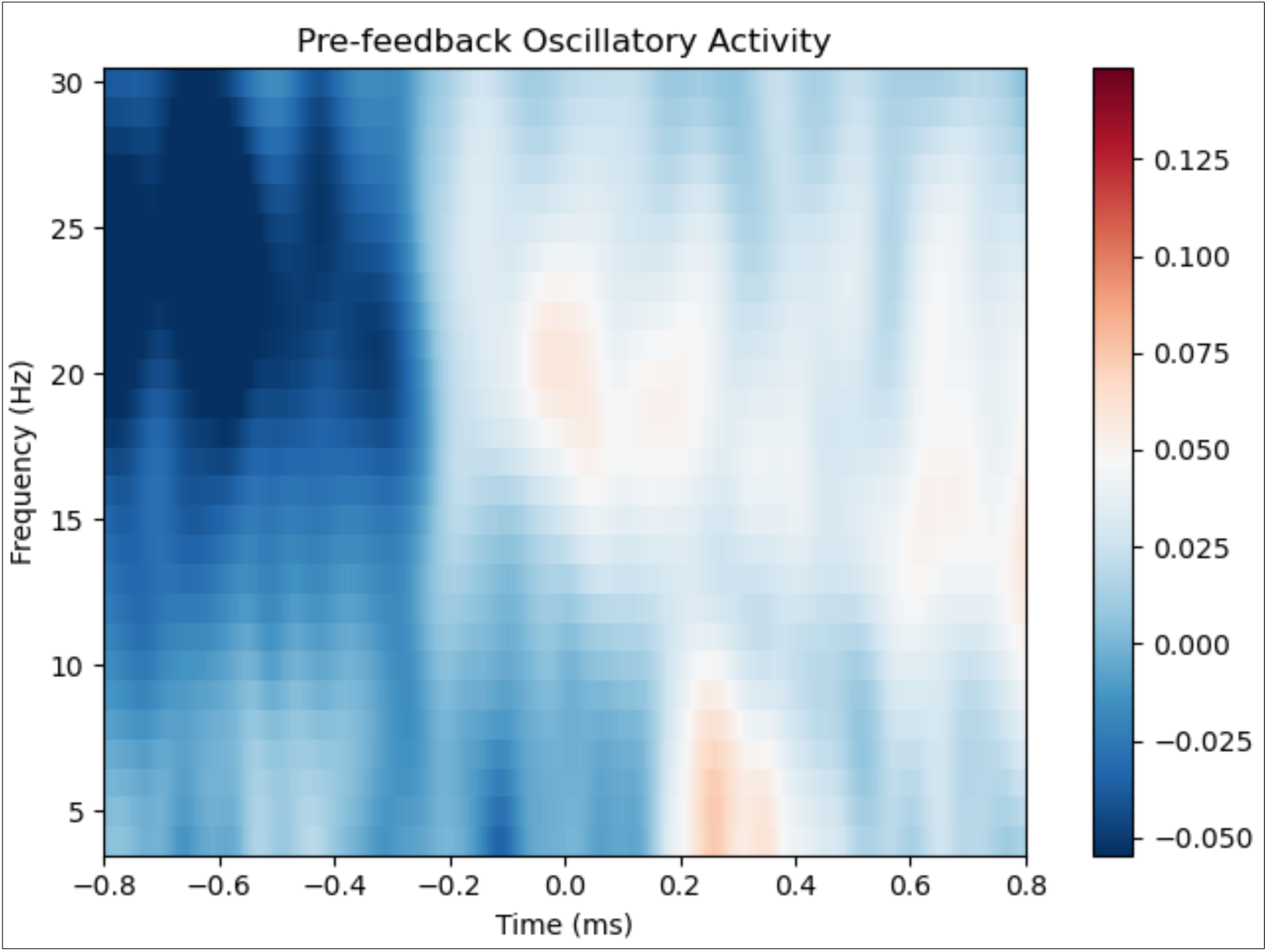
Broadband oscillatory differences across pre-feedback, feedback onset and post-feedback epoch time. Time in milliseconds in presented on the x-axis, with 0 reflecting beginning of feedback (i.e., animated rising bar). Frequency in presented on the y-axis ranging between 4-30Hz. Scaled power is presented on the z-axis, with warmer red shades reflecting higher power and cooler blue shades reflecting lower power.

### 3.3 Primary statistical models

#### 3.3.1 Pre-feedback alpha and beta power

It was hypothesized that pre-feedback alpha and beta power would be modulated by error feedback predictability (tested via model 1 and 2). Model 1 (pre-feedback alpha model) revealed significant interactions between pre-feedback alpha power and the high predictability condition (*p* = 0.001, β = -0.22, *SE* = 0.05, CI = 95% [-0.31 – -0.13]) and between pre-feedback alpha power and the low predictability condition (*p* = 0.014, β = 0.10, *SE* = 0.04, CI = 95% [0.02 – 0.18]), see Appendix A for the full model output. As is apparent from Figure 6, this reflects that in high predictability trials, higher pre-feedback alpha power was linked to lower performance while in low predictability trials, higher pre-feedback alpha power was associated with higher performance. Model 2 (pre-feedback beta model) revealed a significant interaction between pre-feedback beta power and low predictability items on performance points (*p* = 0.037, β = 0.06, *SE* = 0.03, CI = 95% [0.00 – 0.12]), with results reflecting that in low predictability trials, increased pre-feedback beta power was associated with higher performance points. See Appendix B for the full Model 2 output. This interaction is visualized in Figure 7. Additionally, a pre-feedback theta model was conducted for consistency. This model revealed no effect of interest to this paper’s research question and hypotheses (*p* = 0.052, β = 0.01, *SE* = 0.02, CI = 95% [0.03 – 0.05]), see Appendix F for full model output.

**Figure 6.**
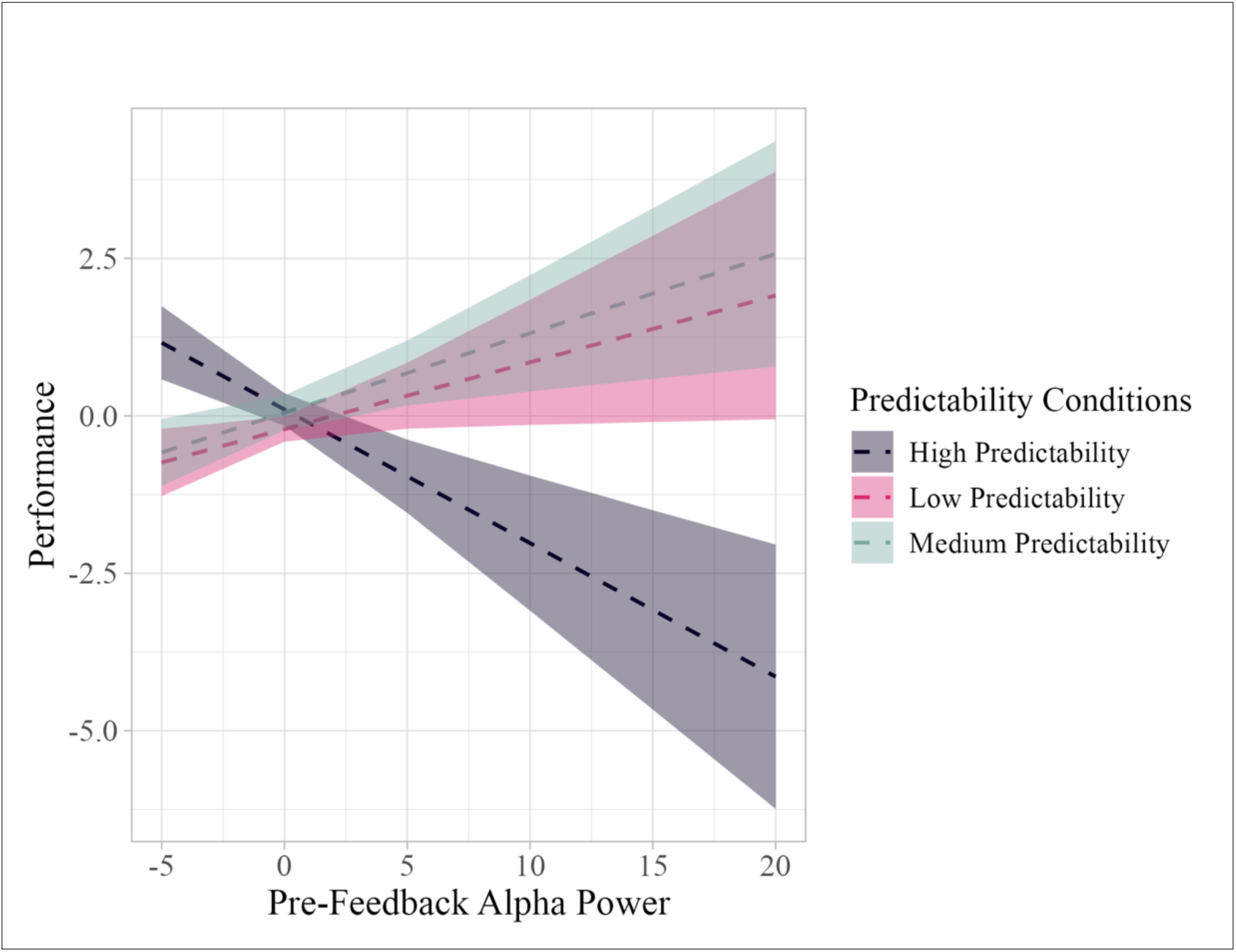
The interaction between pre-feedback alpha power and predictability levels. Pre-feedback alpha power is presented on the x-axis, with higher values reflecting higher alpha power. Scaled performance points are presented on the y-axis, with higher values reflecting better performance. The shaded regions reflect 83% confidence intervals. The different colors represent the different predictability conditions, with purple reflecting high predictability, pink reflecting low predictability and green reflecting medium predictability

**Figure 7.**
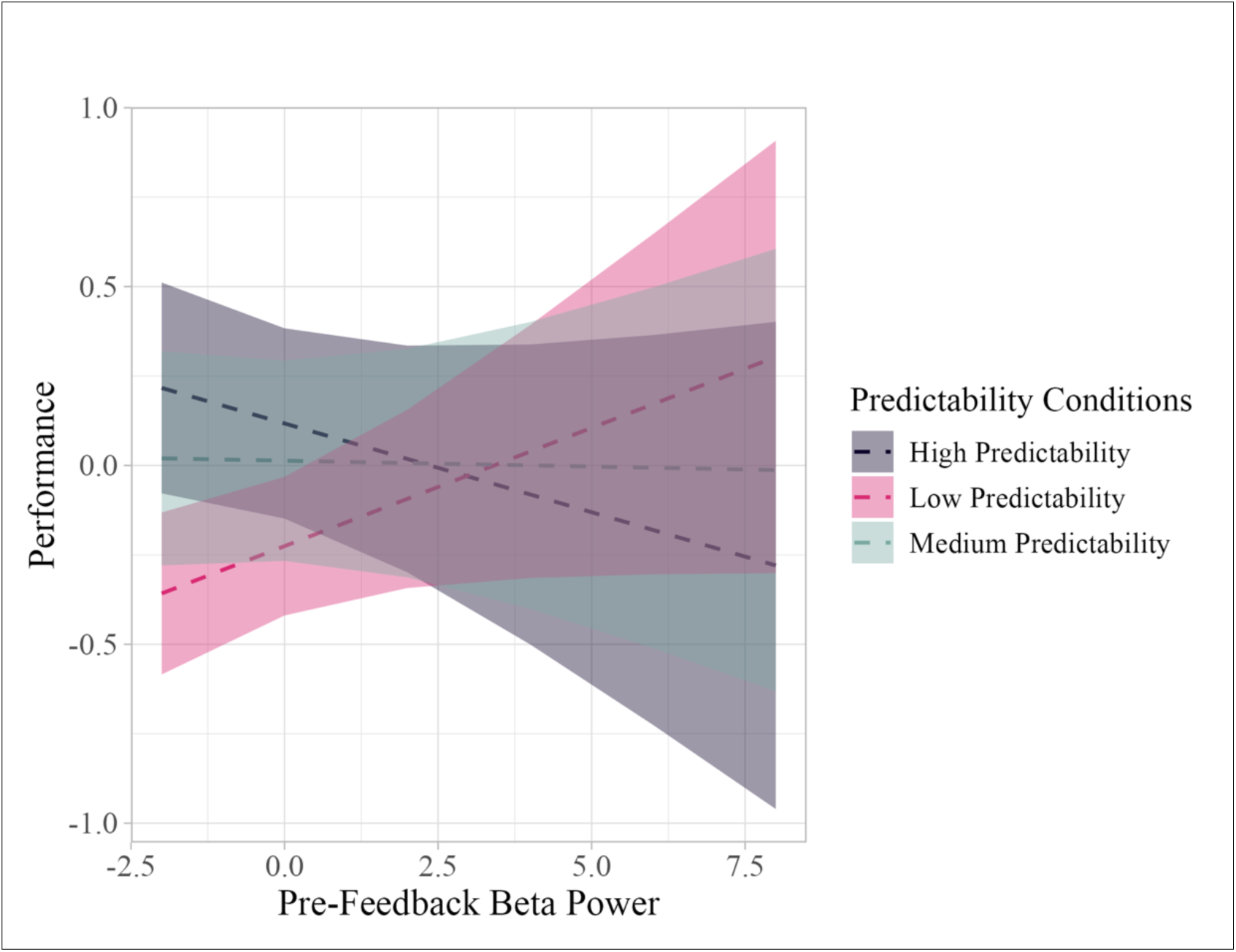
The interaction between pre-feedback beta power and predictability levels. Pre-feedback beta power is presented on the x-axis, with higher values reflecting higher alpha power. Scaled performance points are presented on the y-axis, with higher values reflecting better performance. The shaded regions reflect 83% confidence intervals. The different colors represent the different predictability conditions, with purple reflecting high predictability, pink reflecting low predictability and green reflecting medium predictability.

Model 2 (i.e., pre-feedback beta model) revealed a significant interaction between pre-feedback beta power and performance in the low and high predictability conditions; however, upon visual inspection of the plotted output of this model (see Figure 7), the plot illustrated overlapping confidence intervals, indicating that this relationship may not be statistically meaningful. Thus, a planned comparisons of high and low predictability regression slopes was conducted using the function *emtrends* from the *emmeans* (Lenth & Piaskowski, 2025) package. This estimated marginal means contrast, revealed a significant difference between the high and low predictability levels (Δ= −0.116, SE = 0.06, *t*(883) = −2.04, *p* = 0.042), reflecting that the high predictability condition was associated with lower performance in comparison to the low predictability condition.

#### 3.3.2 feedback-related theta power

It was hypothesized that feedback-related theta power would be higher for lower performance points (i.e., error feedback) in comparison to higher performance points (i.e., correct feedback; tested via Model 3). Model 3 revealed a significant interaction between performance and midline feedback-related theta power (*p* = 0.038, β = -0.06, *SE* = 0.03, CI = 95% [-0.11 – 0.00]), with results reflecting that lower performance was linked to higher feedback-related theta power, while higher performance was linked to lower feedback-related theta power. See Figure 8 for the illustration of Model 3 results and Appendix C for the full output of Model 3.

**Figure 8.**
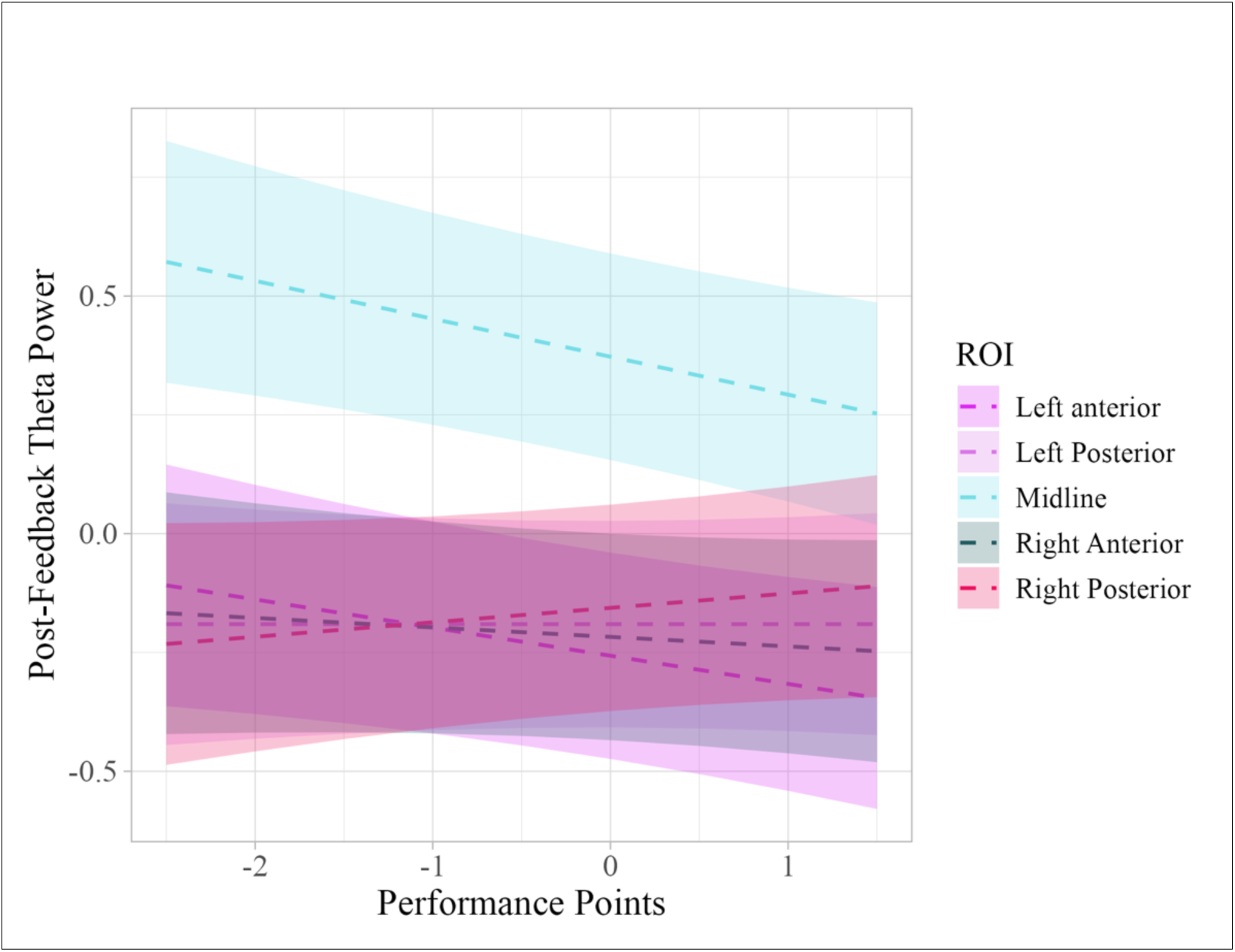
The effect of feedback on feedback-related theta power. Scaled performance points are presented on the x-axis, with higher values reflecting better performance. Feedback-related theta power is presented on the y-axis, with higher values reflecting higher theta power. The shaded regions reflect 83% confidence intervals. The different colors represent the different ROIs, with light and dark pink reflecting left anterior and posterior regions, blue reflecting midline region, green reflecting right anterior and red reflecting right posterior.

#### 3.3.3 Exploratory analysis - alpha and beta power across the course of learning

Plot and model outputs from the primary analysis of LMM 1 revealed an unexpected difference in pre-feedback alpha power across conditions, where there was a signification interaction between pre-feedback alpha power and high predictability stimuli, with results indicating that lower pre-feedback alpha power was associated with higher performance, while higher pre-feedback alpha power was associated with lower performance (see Figure 6). To further explore if participants adapted their learning over time to different predictability conditions, additional exploratory LMMs were conducted to analyze whether trial interacted with condition and pre-feedback alpha (i.e., Model 4) and beta power (i.e., Model 5) to predict performance.

Exploratory analyses involved 2 models identical in structure to those in the primary analysis, with trial allowed to interact with ROI, predictability condition and performance points rather than being treated as a covariate. See Figure 9 for the alpha and beta model visualization.

**Figure 9.**
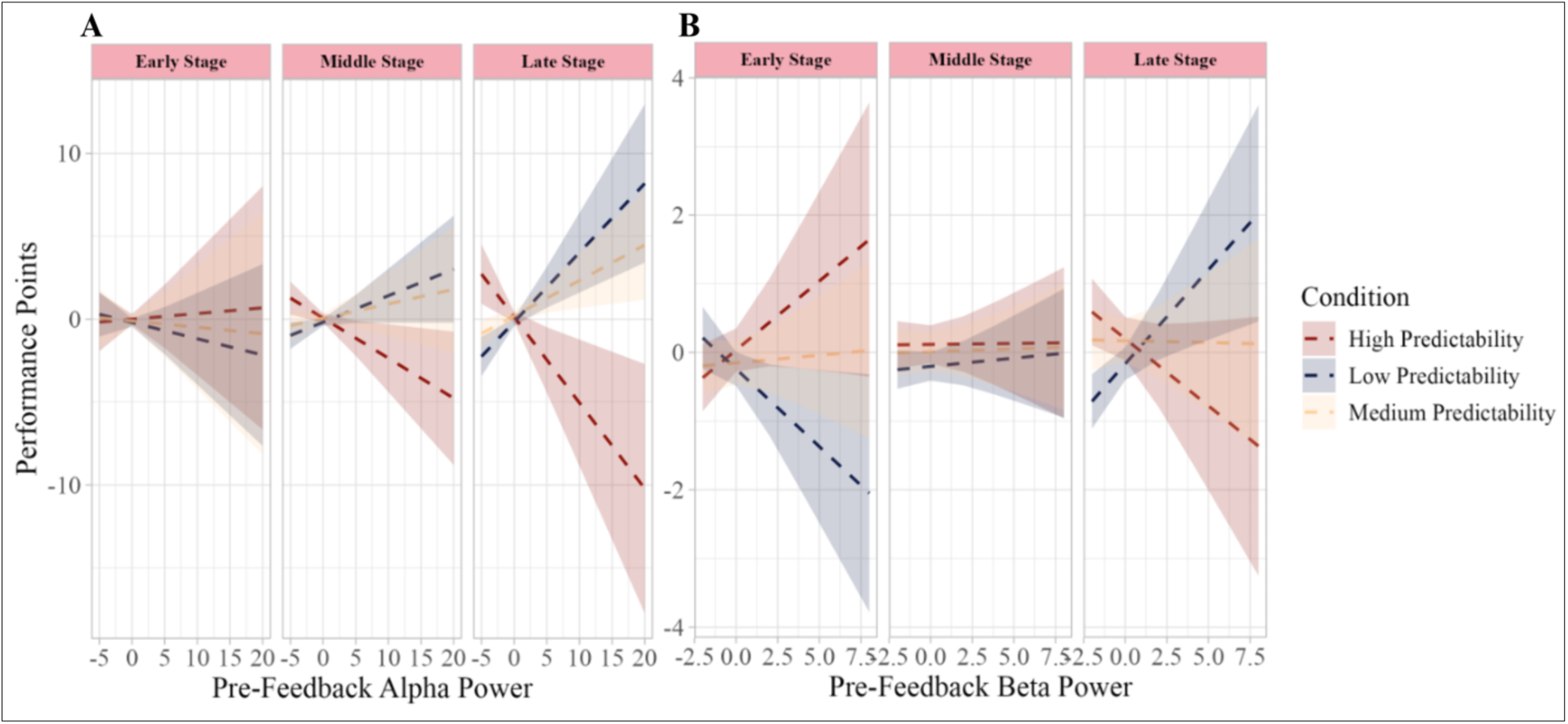
Interaction effect of pre-feedback alpha and beta, condition and trial on performance. Pre-feedback alpha power (A) and beta power (B) is presented on the x-axis, performance points presented on the y-axis. Facets are the different stages of learning across time (early, middle, and late). Shaded regions reflect confidence intervals of 83%.

### 3.4 Exploratory statistical models result

#### 3.4.1 Pre-feedback alpha and beta power

Exploratory models were conducted to further explore if trial (i.e., time-resolved learning across the experiment), condition, pre-feedback alpha (as tested via model 4) and beta (as tested via model 5) interacted to predict performance. See Appendix D for the full output of Model 4 and Appendix E for Model 5. There was a significant interaction between pre-feedback alpha power, high predictability condition and trial on performance (*p* = 0.001, β = -0.15, *SE* = 0.04, CI = 95% [-0.23 – -0.08]), and between pre-feedback alpha power, low predictability condition and trial on performance (*p* = 0.001, β = 0.11, *SE* = 0.03, CI = 95% [0.04 – 0.17]). As apparent from Figure 9A, this reflects that in the late learning stage, higher pre-feedback alpha was associated with higher performance when items were unpredictable, while higher alpha power was linked with lower performance when items were highly predictable. Similarly, model 5 also yielded a significant interaction between pre-feedback beta power, low predictability and trial on performance (*p* = 0.001, β = 0.09, *SE* = 0.02, CI = 95% [0.05 – 0.14]), and between pre-feedback beta power, high predictability and trial on performance (*p* = 0.003, β = -0.07, *SE* = 0.02, CI = 95% [-0.12 – -0.02]). As apparent from Figure 9B, this reflects that in the late stage of learning, higher pre-feedback beta was associated with higher performance when items were unpredictable, while higher beta power was linked with lower performance when items were highly predictable.

## 4. Discussion

This study aimed to examine pre- and feedback-related electrophysiological correlates of error processing and determine whether predictability (and inferred certainty) interacts with these correlates in relations to the memory updating process. To this end, we reanalyzed data previously published by Hassall and colleagues (2023), who investigated the neural correlates of continuous feedback using ERPs. The present findings revealed that feedback-related theta power was influenced by performance points, such that higher feedback-related theta power was linked to lower performance and vice versa. Additionally, results demonstrated a positive relationship between power and performance in the low predictability condition for both alpha and beta, but the inverse relationship for high predictability. We noted that the pre-feedback alpha pattern emerged over the course of learning, while the pre-feedback beta pattern was reversed. In the following, we will first discuss the pre-feedback alpha and beta patterns across the course of learning before moving onto discussing the feedback-related theta results. We will then discuss limitations and future directions before offering some conclusions.

### 4.1 Pre-feedback activity

It was hypothesized that pre-feedback alpha and beta power would be modulated by performance feedback and predictability levels. Our findings support this hypothesis, revealing a significant interaction between pre-feedback alpha and beta power, condition and trial on performance.

These findings are consistent with prior literature linking pre-stimulus neural activity to subsequent memory outcomes (Scholz et al., 2017; Ostrowski & Rose, 2024; Salari & Rose, 2016; Weismüller et al., 2019; Toosi et al., 2017; Baslar & Stampfer, 1985), as well as literature that demonstrates a relationship between memory certainty and pre-stimulus alpha and beta power (Toosi et al., 2017). Here, we reveal a link between high performance points and high pre-feedback alpha and beta power in the low predictability condition, suggesting that the inhibition-mediated increases via pre-feedback alpha power, may be more fruitful when items are more difficult to anticipate in comparison to when items are easily anticipated. Additionally, it is plausible to suggest that due to the unpredictable nature of these items, participants might have had to employ a preparatory strategy to further allocate attentional resources to the to-be-encoded items (Scholz et al., 2017; Ostrowski & Rose, 2024; Weismüller et al., 2019; Salari & Rose, 2016). The increased pre-feedback alpha and beta power linked with the low predictability condition in our study aligns with prior literature that demonstrated an increase in pre-stimulus beta power for uncertain trials in comparison to certain ones (Helvert et al., 2021; Palmer et al., 2019). However, does not align with prior studies that illustrate an increase in pre-stimulus alpha power for certain trials in comparison to uncertain trials. (Toosi et al., 2017). It is also possible that low predictability items are difficult to learn, and thus, require more allocated attentional resources and preparatory mechanisms, which are achieved via the inhibition-mediated increases via pre-feedback alpha and preparatory processes via beta power. Additionally, it is plausible that the higher performance linked with higher alpha and beta power in the low predictability condition could be due to the inhibition of irrelevant and already encoded stimuli in order to dedicate more selective attention to the yet-to-be-learnt stimuli (i.e., performance feedback following response) and due to participants employing further preparatory mechanisms to encode upcoming performance feedback, as they might not be certain of their response and they might be further prepared to encode feedback on their performance. Overall, low predictability items in this study may require further selective attention and preparatory mechanisms to be successfully encoded, which is reflected by the higher performance points.

At a descriptive level, the significant relationship between pre-feedback alpha and beta power, condition and trial on performance suggests that memory certainty plays a role in how the brain processes performance feedback, influencing the process of memory updating and in turn, performance. That the directionality of effects differed as a function of predictability may suggest that participants employed strategy adaptation based on their certainty of the outcome. This modulation, especially towards the end of the experimental session may also explain inconsistencies with prior findings that revealed higher alpha and beta power were linked with better memory performance (Ostrowski & Rose, 2024; Toosi et al., 2017; Baslar & Stampfer, 1985). An inhibition-based account may explain the lower performance that is associated with the higher pre-feedback alpha and beta power. In short, participants may have employed an inhibition strategy (reflected by the high alpha and beta power for high predictable items at the late learning stage) where already learned, highly predictable items are down-weighted to facilitate allocation of attention resources to upcoming information (Toosi et al., 2017). This interpretation aligns with what is already known in the literature (i.e., an increase in pre-stimulus alpha and beta power reflects inhibition of already learnt information in favor of learning new information; Ostrowski & Rose, 2024; Toosi et al., 2017; Baslar & Stampfer, 1985). However, the present findings suggest that such an inhibition strategy may be sub-optimal when items are highly predictable. When items are easily anticipated, further inhibition of predictable information may be unnecessary and as such stimuli provide limited new information and do not require further processing (Geng et al. & Behrmann, 2005). Elevated alpha and beta power for highly predictable items may reflect disengagement from feedback-related processing rather than adaptive preparatory control, consistent with the observed reduction in performance.

Predictive coding models offer a sound account of why predictability modulates anticipatory neural dynamics. These models propose that predictions about incoming sensory input are continuously generated and updated at each stage of visual processing, suggesting that highly predictable input, elicits reduced neural processing because it conveys little new information whether it is relevant or not, making predictable input less behaviorally relevant (Friston, 2010; Summerfield & Egner, 2009; Noonan et al., 2018; Rao & Ballard, 1999; Hutchinson & Turk-Browne, 2012; Stokes et al., 2012). From a predictive coding perspective, predictable input is inhibited due to it being treated like a distractor, as it is less behaviorally relevant and new.

Treating highly predictable items as distractors may explain the lower performance observed with higher alpha and beta power, as such items may be deemed irrelevant when no updating is required, consistent with increased alpha reflecting inhibitory attentional gating of predictable input (van Moorselaar & Slagter, 2020; Noonan et al., 2018), and increased beta reflecting the maintenance of stable predictions and the anticipation of expected input, resulting in a reduced need for updating (Arnal & Giraud, 2012). More specifically, the link between both high alpha and beta power and low performance points for highly predictable items in this study aligns with a prior study which demonstrated that employing an inhibition strategy for highly anticipatory outcomes is not effective, suggesting that the highly anticipatory items remove the need for further processing, which may result in worse memory performance (Noonan et al., 2018). This argument also aligns with prior studies which demonstrated increased pre-stimulus alpha and beta power for highly anticipatory events (Wang et al., 2019; Geng et al. & Behrmann, 2005; Wessel, 2018; Toosi et al., 2017).

### 4.2 Alpha and beta power: separate anticipatory mechanisms

At high level, pre-feedback alpha and beta power showed convergent patterns later in learning. This could in part, suggest there are overlapping roles in anticipatory regulation. When only considering the late learning stage alone, higher alpha and beta power were similarly associated with performance in a predictability dependent manner consistent with the view that both frequency bands can index anticipatory attention allocation or regulation prior to feedback are attention-allocating mechanisms, preparing the brain to selectively assign attention to upcoming stimuli (Ostrowski & Rose, 2024; Salari & Rose, 2016). However, examining learning-stage effects reveals important differences between alpha- and beta-related dynamics. In the exploratory alpha model (Model 4) pre-feedback alpha power showed little association with performance and predictability early in learning, suggesting that alpha-mediated inhibition strategies may not be strongly enacted when task structure is still being acquired. In short, participants at the beginning may be using a flexible, adaptable processing strategy where they encode all stimuli without the need to inhibit any information, as at the early learning stage all information is still new, and participants are learning the stimuli’s predictability patterns. This interpretation aligns with the notion that early learning benefits from minimal top-down constraint when all outcomes are uncertain.

In contrast, the beta model (i.e., Model 5) illustrates a clear link between pre-feedback beta power, performance and condition with a difference between low and high predictability conditions. This suggests that there are differences in the pre-feedback alpha and beta roles at the beginning of learning a new task, before task familiarity is established. However, the present data do not afford firm conclusions at this stage. A further difference we noted between pre-feedback alpha and beta is in the late learning stage. Although the alpha and beta models illustrate a similar pattern in the late learning stage, the medium predictability condition showed a slightly different pattern depending on the frequency band, where the beta model (i.e., Model 5) at this stage does not follow the same pattern as the low predictability condition, as is the case for the alpha model (i.e., Model 4). It is not clear why the medium predictability condition pattern differs based on the frequency band while the low and medium condition patterns are the same for both alpha and beta in late learning. However, one possible explanation is that, although both alpha and beta are implicated in attentional control, they subserve partially distinct mechanisms. Beta activity has been linked to the maintenance of the status quo and preparatory processes (Scholz et al., 2017; Ostrowski & Rose, 2024; Salari & Rose, 2016), whereas alpha is more closely associated with inhibitory gating in favor of selective processing (Jensen & Mazaheri, 2010). In a medium predictability context where outcomes are more uncertain, these mechanisms may be differently engaged, resulting in the observed divergence between alpha and beta.

### 4.3 feedback-related theta power

It was hypothesized that feedback-related theta power would be higher for error feedback in comparison to correct feedback. Our results support this hypothesis, as Model 3 revealed a significant effect where performance predicted feedback-related theta power. More specifically, we found that higher feedback-related theta power was linked to lower performance, whereas lower feedback-related theta was linked to better performance. This suggests that feedback modulates feedback-related theta power and, in turn, may have an influence on memory updating. These results replicate findings from prior studies that demonstrated a link between post-feedback theta power and error feedback, where theta power increased for error feedback in comparison to correct feedback, positing that the increase in theta power is indexing memory updating (Bush et al., 2002; Cavanagh et al., 2009, 2012; Luu et al., 2003; van de Vijver et al., 2011; Wang et al., 2017; van de Vijver et al., 2011). This present study extend this literature by showing that theta is similarly sensitive to both performance information under continues, accumulative feedback and discrete feedback.

While our pre-feedback alpha and beta findings are interpreted within a predictive coding framework, modulation of feedback-related theta power may reflect not only surprisal, but also behavioral adjustment in response to feedback. This idea aligns with prior studies suggesting that midline and midfrontal theta power reflects behavior adjustment (Cavanagh & Frank, 2014) and the degree of behavioral adaptation not just error magnitude or surprisal (van de Vijver et al., 2011). Therefore, the underlying neural mechanisms of discrete and continuous accumulative feedback may be similar, as both have been shown to evoke theta power. While this does not preclude more subtle mechanistic differences between these feedback types, it suggests broad commonalities in their neurophysiological effects, particularly in relation to the updating of memory representations.

### 4.4 Limitations and future directions

While the present study provides novel insights into the role of feedback and predictability levels in the process of memory updating and extends existing knowledge regarding the underlying mechanism of memory updating, methodological characteristics need to be considered when interpreting the current findings. For example, the reward predictability distribution linked with the medium predictability gnomes may not have been distinct enough from the reward predictability distribution linked with the high predictability gnomes. More specifically, medium predictability distribution was generated through combining the same two distributions of the high predictability gnomes with 80/20 weighting (Hassall et al., 2023; see Figure 1). As a result, this may have caused the medium-reward predictability cues to be too close to those of high-reward predictability cues, making it difficult to isolate the distinct influence of the high- and medium-predictability conditions on pre-feedback alpha and beta power. Future studies should attempt to replicate this design while ensuring that the distributions between predictability conditions do not overlap and are not too close. Doing so would allow clearer interpretations to be drawn. Additionally, aperiodic activity was not controlled for in the current analysis. This limits the specificity of our oscillatory interpretations, as EEG power reflects a combination of periodic (oscillatory) and aperiodic (1/f) components (Voytek et al., 2015; Donoghue et al., 2020). Consequently, observed changes in band-limited power may partly arise from shifts in the aperiodic background, rather than reflecting true oscillatory modulation (Voytek et al., 2015; Donoghue et al., 2020).

### 4.5 Conclusion

The present study sought to shed light on the underlying neural mechanisms of memory updating, emphasizing the role of pre-feedback oscillatory activity in conjunction with distinct predictability levels. Our findings provide evidence for the involvement of pre-feedback alpha and beta power in the memory updating process, with them illustrating a similar pattern at the end of learning but a different trajectory in the beginning of learning which might suggest that they are separable mechanisms, respectively. This study demonstrates the potentially different involvement of alpha and beta power across the course of learning, a novel finding that suggests a learning adaptation strategy reflected in these two neural correlates. Our findings illustrate that feedback-related theta power responds similarly to accumulative and discrete feedback, indicating potentially shared neural mechanisms. These findings advance our understanding of the neural mechanisms underlying memory, demonstrating that pre-feedback alpha and beta power linked to certainty, shape feedback processing and subsequent updating, and highlight feedback-related theta power as a shared neural correlate for both accumulative and discrete feedback.

## Appendix A: Model 1 output summary

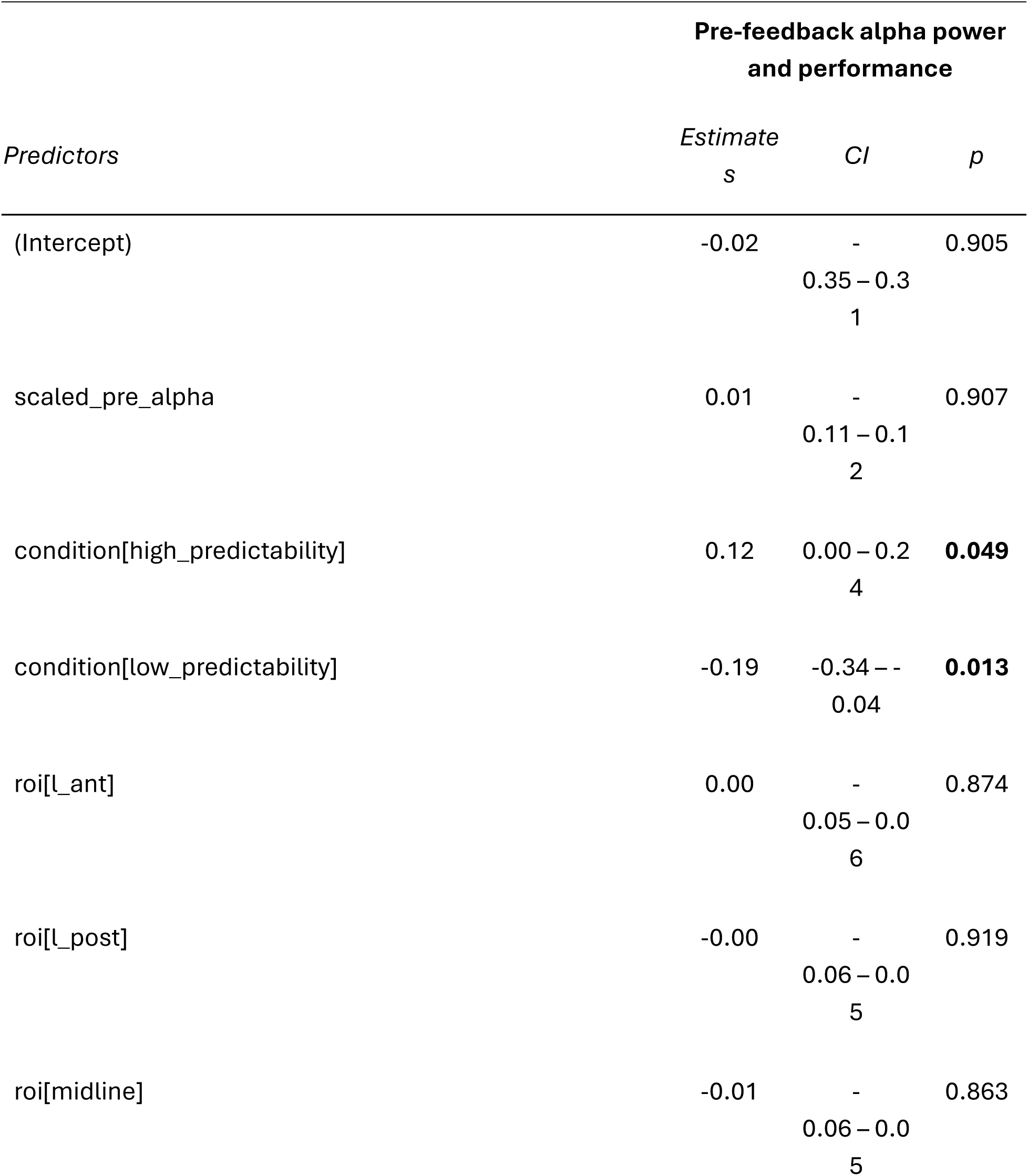
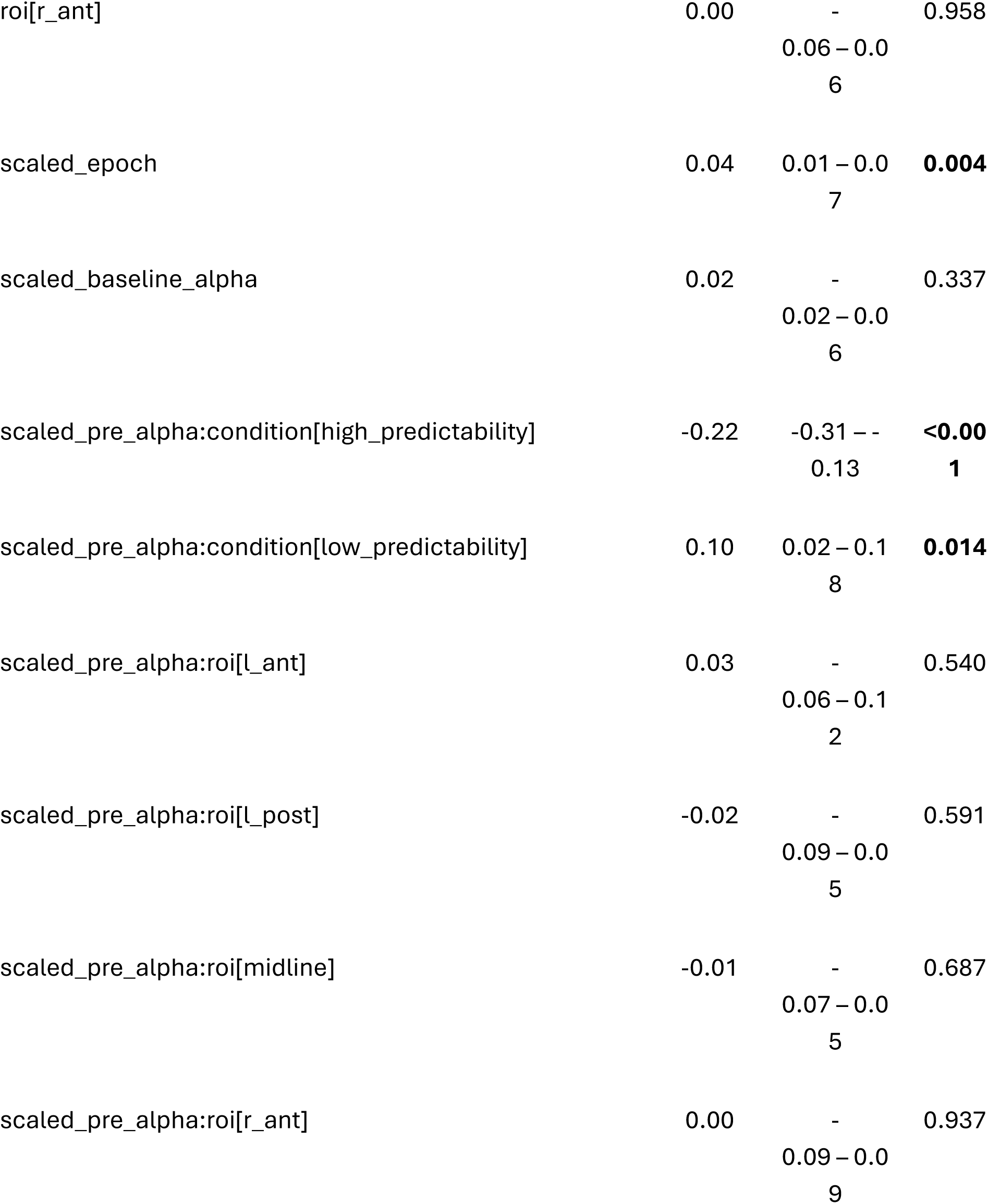
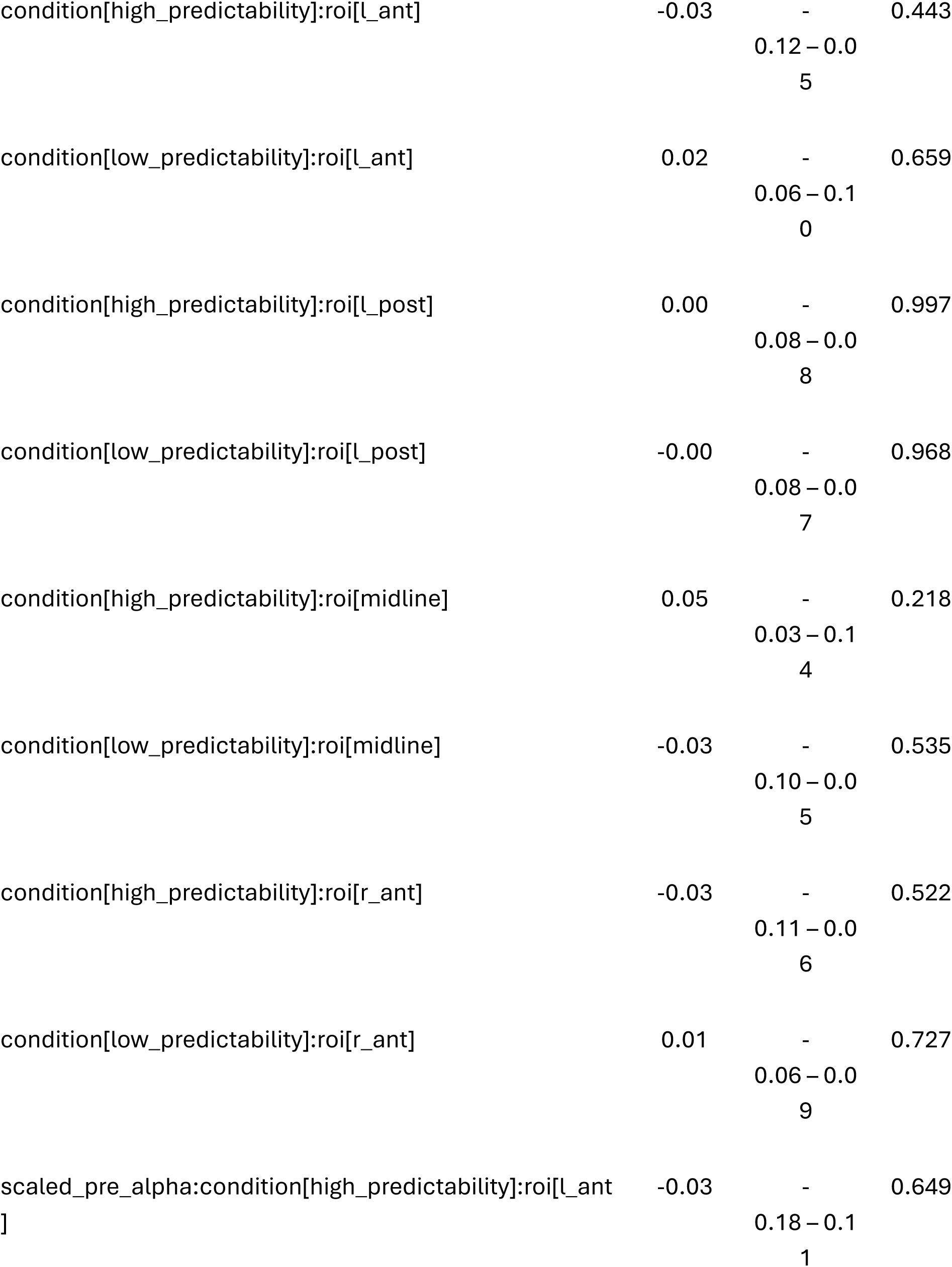
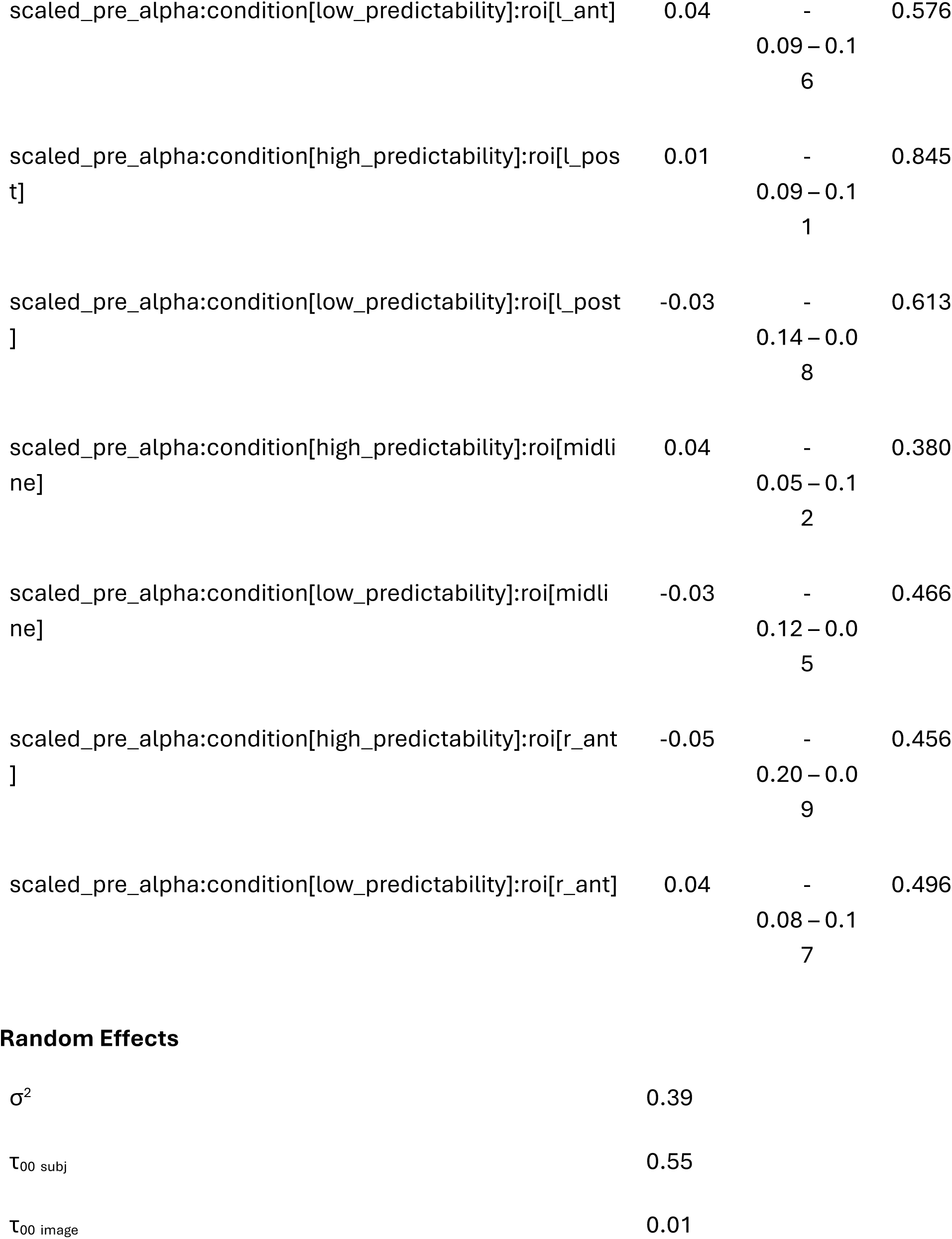
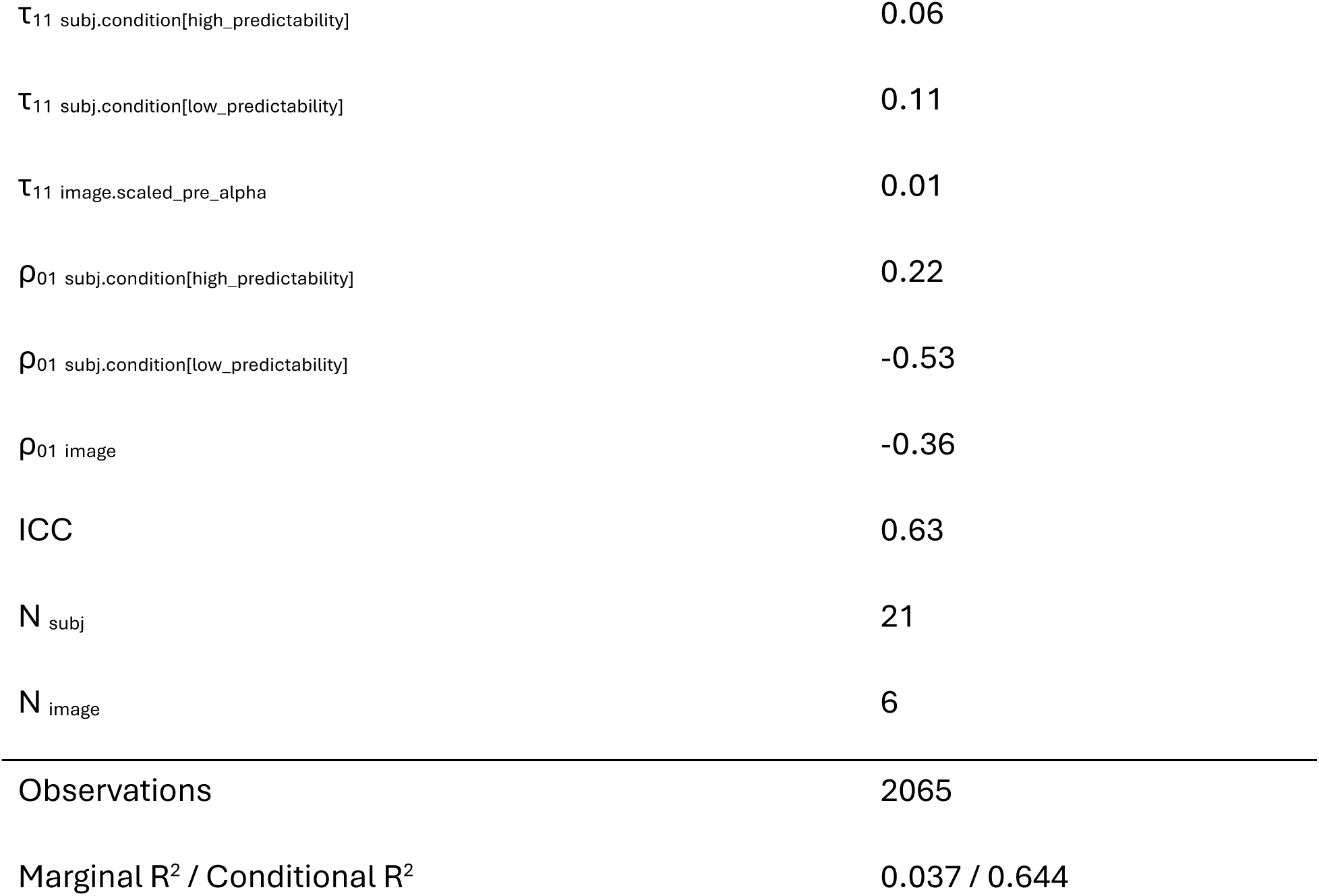

| *Predictors* | Pre-feedback alpha power and performance |  |  |
| --- | --- | --- | --- |
|  | *Estimate*<br>*s* | *CI* | *p* |
| (Intercept) | -0.02 | -<br>0.35 – 0.31 | 0.905 |
| scaled_pre_alpha | 0.01 | -<br>0.11 – 0.12 | 0.907 |
| condition[high_predictability] | 0.12 | 0.00 – 0.24 | **0.049** |
| condition[low_predictability] | -0.19 | -0.34 – -0.04 | **0.013** |
| roi[l_ant] | 0.00 | -<br>0.05 – 0.06 | 0.874 |
| roi[l_post] | -0.00 | -<br>0.06 – 0.05 | 0.919 |
| roi[midline] | -0.01 | -<br>0.06 – 0.05 | 0.863 |
| roi[r_ant] | 0.00 | -<br>0.06 – 0.0<br>6 | 0.958 |
| scaled_epoch | 0.04 | 0.01 – 0.0<br>7 | **0.004** |
| scaled_baseline_alpha | 0.02 | -<br>0.02 – 0.0<br>6 | 0.337 |
| scaled_pre_alpha:condition[high_predictability] | -0.22 | -0.31 – -<br>0.13 | **<0.00**<br>**1** |
| scaled_pre_alpha:condition[low_predictability] | 0.10 | 0.02 – 0.1<br>8 | **0.014** |
| scaled_pre_alpha:roi[l_ant] | 0.03 | -<br>0.06 – 0.1<br>2 | 0.540 |
| scaled_pre_alpha:roi[l_post] | -0.02 | -<br>0.09 – 0.0<br>5 | 0.591 |
| scaled_pre_alpha:roi[midline] | -0.01 | -<br>0.07 – 0.0<br>5 | 0.687 |
| scaled_pre_alpha:roi[r_ant] | 0.00 | -<br>0.09 – 0.0<br>9 | 0.937 |
| condition[high_predictability]:roi[l_ant] | -0.03 | -<br>0.12 – 0.0<br>5 | 0.443 |
| condition[low_predictability]:roi[l_ant] | 0.02 | -<br>0.06 – 0.1<br>0 | 0.659 |
| condition[high_predictability]:roi[l_post] | 0.00 | -<br>0.08 – 0.0<br>8 | 0.997 |
| condition[low_predictability]:roi[l_post] | -0.00 | -<br>0.08 – 0.0<br>7 | 0.968 |
| condition[high_predictability]:roi[midline] | 0.05 | -<br>0.03 – 0.1<br>4 | 0.218 |
| condition[low_predictability]:roi[midline] | -0.03 | -<br>0.10 – 0.0<br>5 | 0.535 |
| condition[high_predictability]:roi[r_ant] | -0.03 | -<br>0.11 – 0.0<br>6 | 0.522 |
| condition[low_predictability]:roi[r_ant] | 0.01 | -<br>0.06 – 0.0<br>9 | 0.727 |
| scaled_pre_alpha:condition[high_predictability]:roi[l_ant<br>] | -0.03 | -<br>0.18 – 0.1<br>1 | 0.649 |
| scaled_pre_alpha:condition[low_predictability]:roi[l_ant] | 0.04 | -<br>0.09 – 0.16 | 0.576 |
| scaled_pre_alpha:condition[high_predictability]:roi[l_post] | 0.01 | -<br>0.09 – 0.11 | 0.845 |
| scaled_pre_alpha:condition[low_predictability]:roi[l_post] | -0.03 | -<br>0.14 – 0.08 | 0.613 |
| scaled_pre_alpha:condition[high_predictability]:roi[midline] | 0.04 | -<br>0.05 – 0.12 | 0.380 |
| scaled_pre_alpha:condition[low_predictability]:roi[midline] | -0.03 | -<br>0.12 – 0.05 | 0.466 |
| scaled_pre_alpha:condition[high_predictability]:roi[r_ant] | -0.05 | -<br>0.20 – 0.09 | 0.456 |
| scaled_pre_alpha:condition[low_predictability]:roi[r_ant] | 0.04 | -<br>0.08 – 0.17 | 0.496 |

Random Effects
|  |  |
| --- | --- |
| $\sigma^2$ | 0.39 |
| $\tau_{00}$ subj | 0.55 |
| $\tau_{00}$ image | 0.01 |
| $\tau_{11}$ subj.condition[high_predictability] | 0.06 |
| $\tau_{11}$ subj.condition[low_predictability] | 0.11 |
| $\tau_{11}$ image.scaled_pre_alpha | 0.01 |
| $\rho_{01}$ subj.condition[high_predictability] | 0.22 |
| $\rho_{01}$ subj.condition[low_predictability] | -0.53 |
| $\rho_{01}$ image | -0.36 |
| ICC | 0.63 |
| $N_{\text{subj}}$ | 21 |
| $N_{\text{image}}$ | 6 |
| Observations | 2065 |
| Marginal $R^2$ / Conditional $R^2$ | 0.037 / 0.644 |

## Appendix B: Model 2 output summary

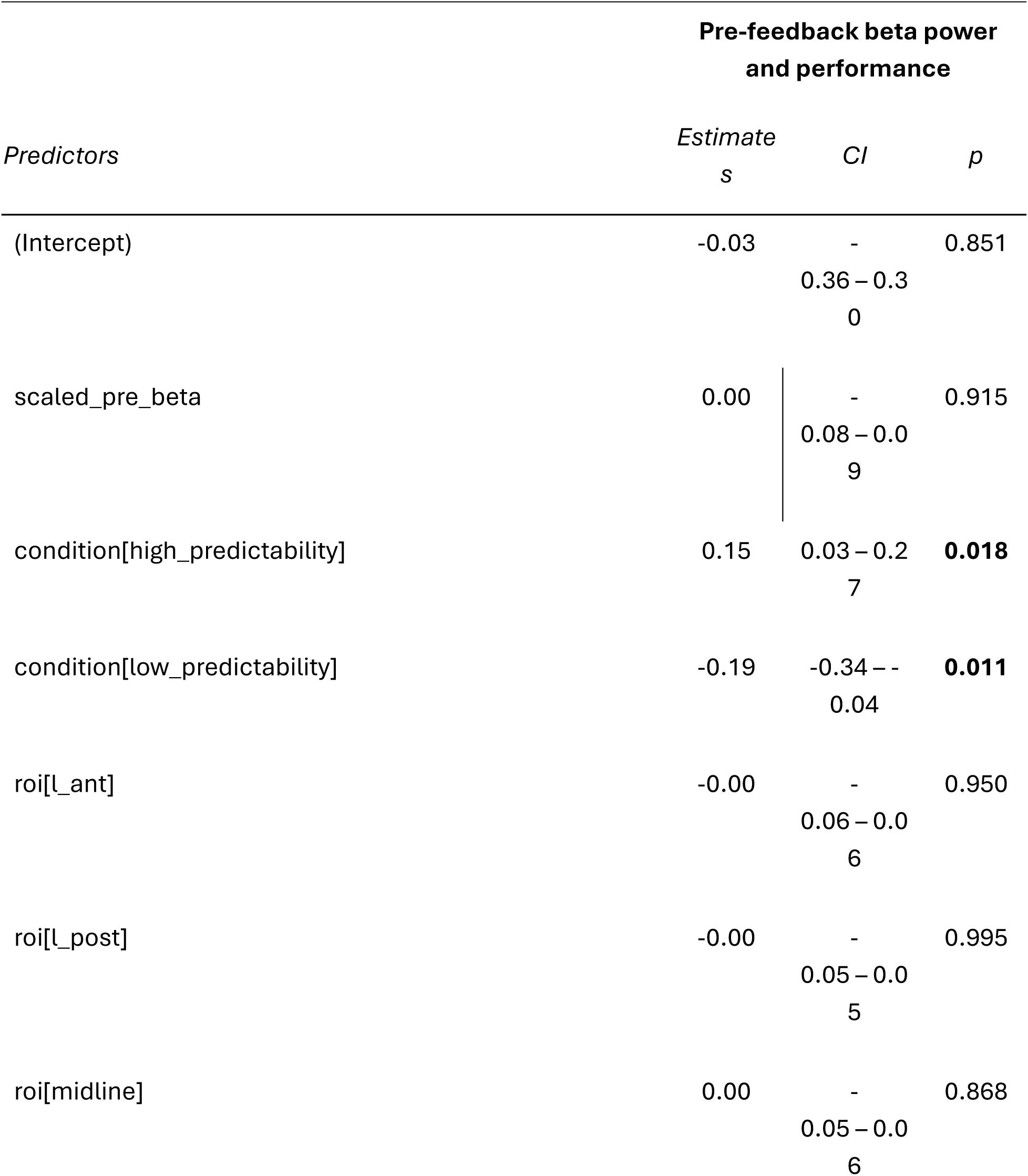
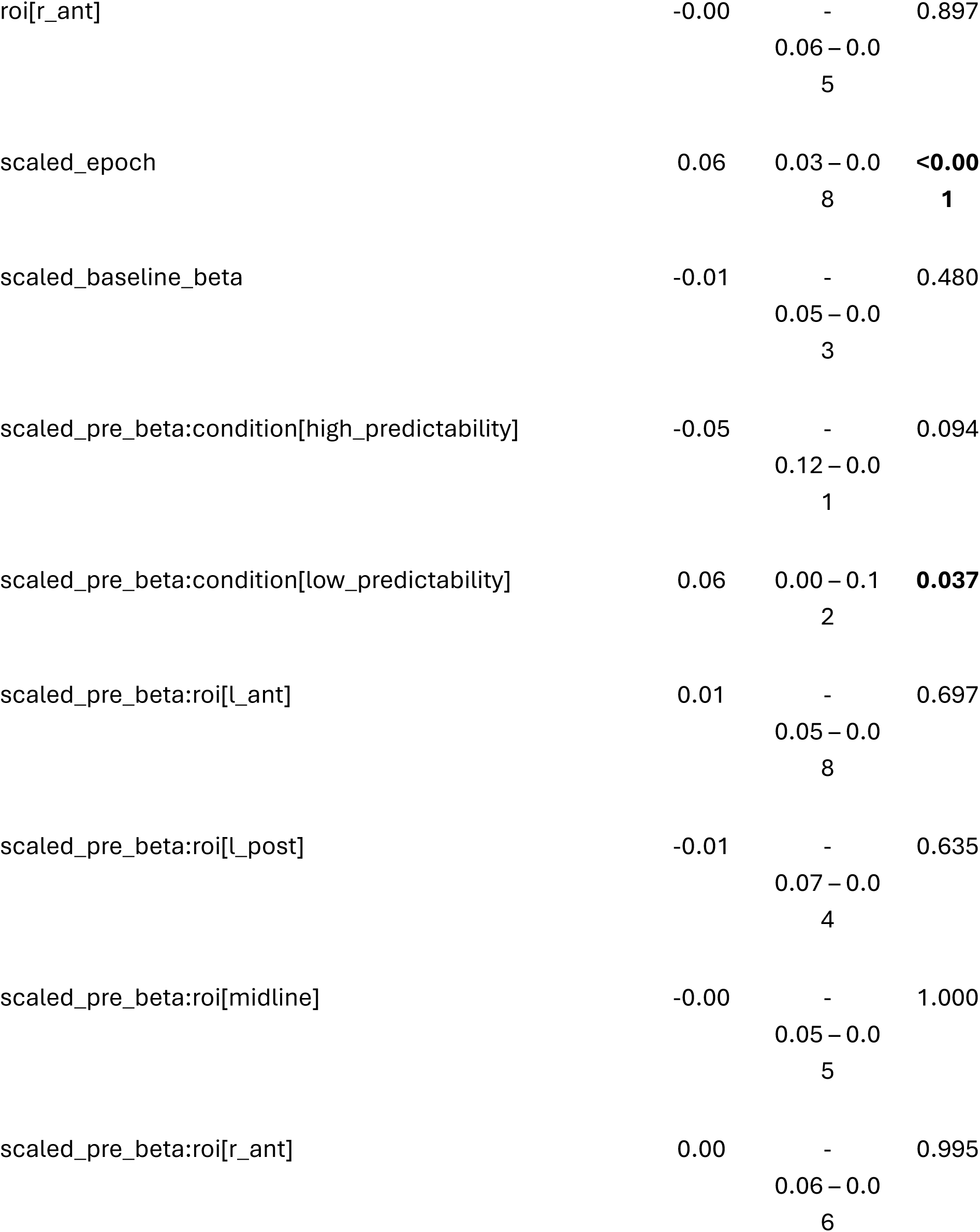
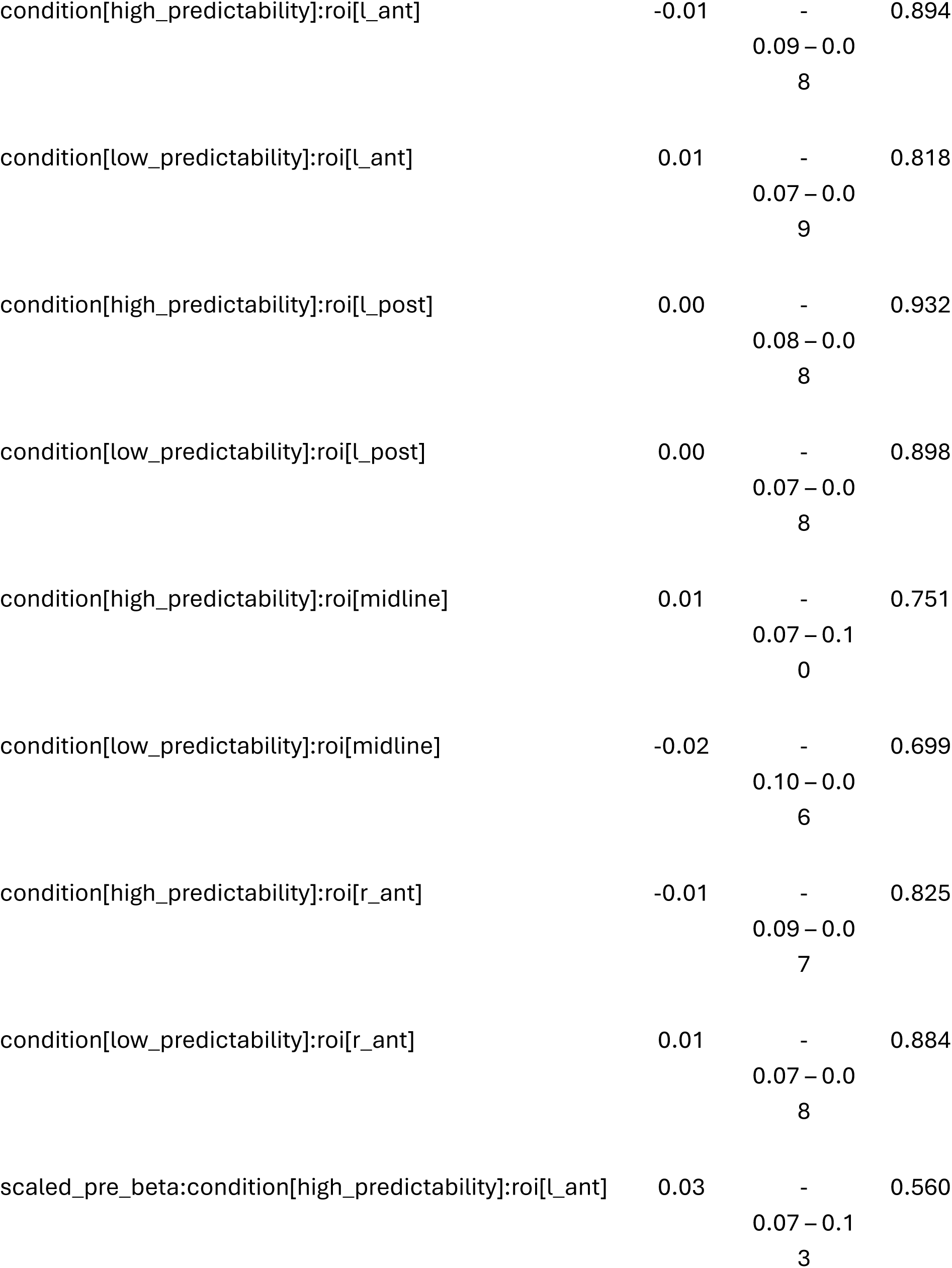
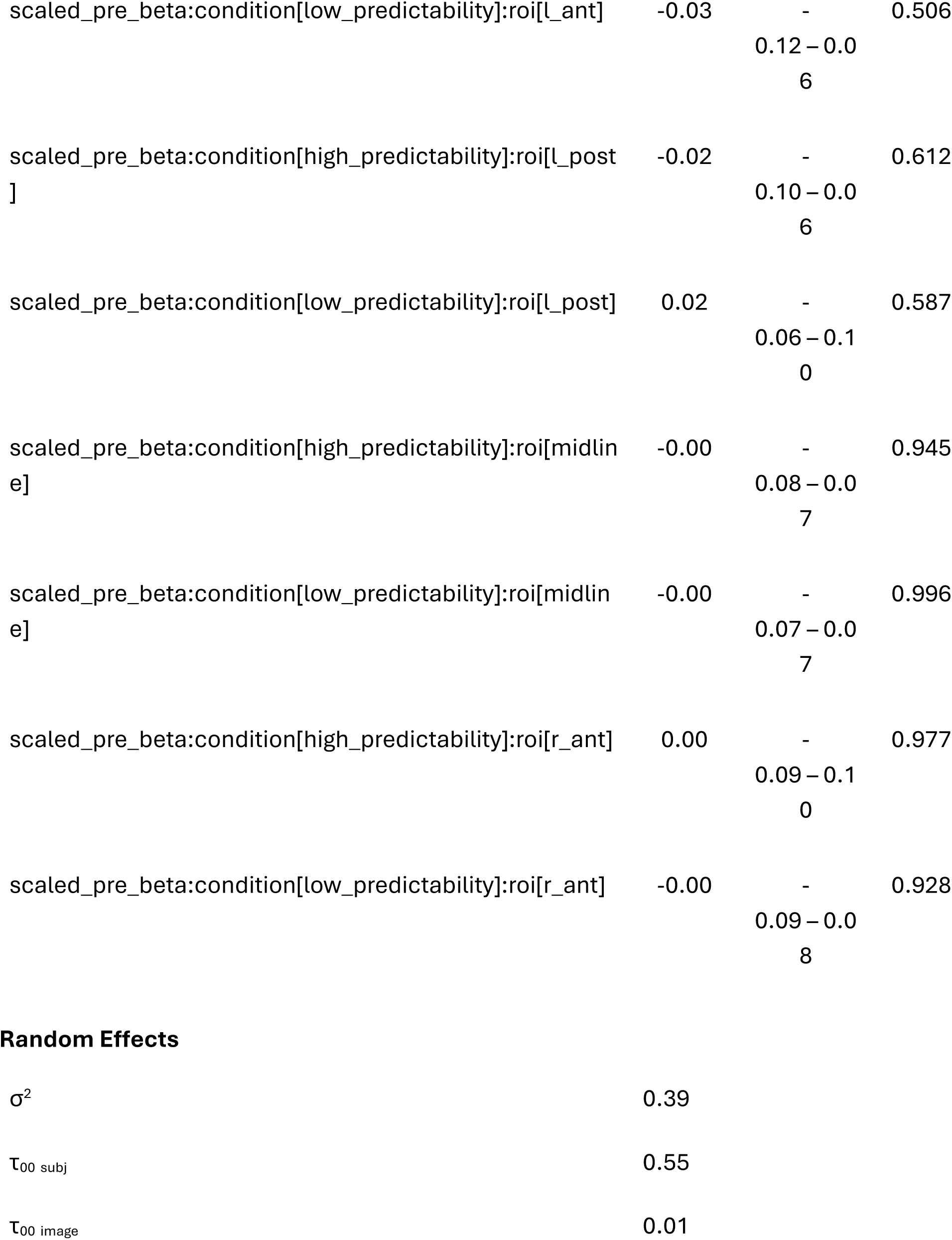
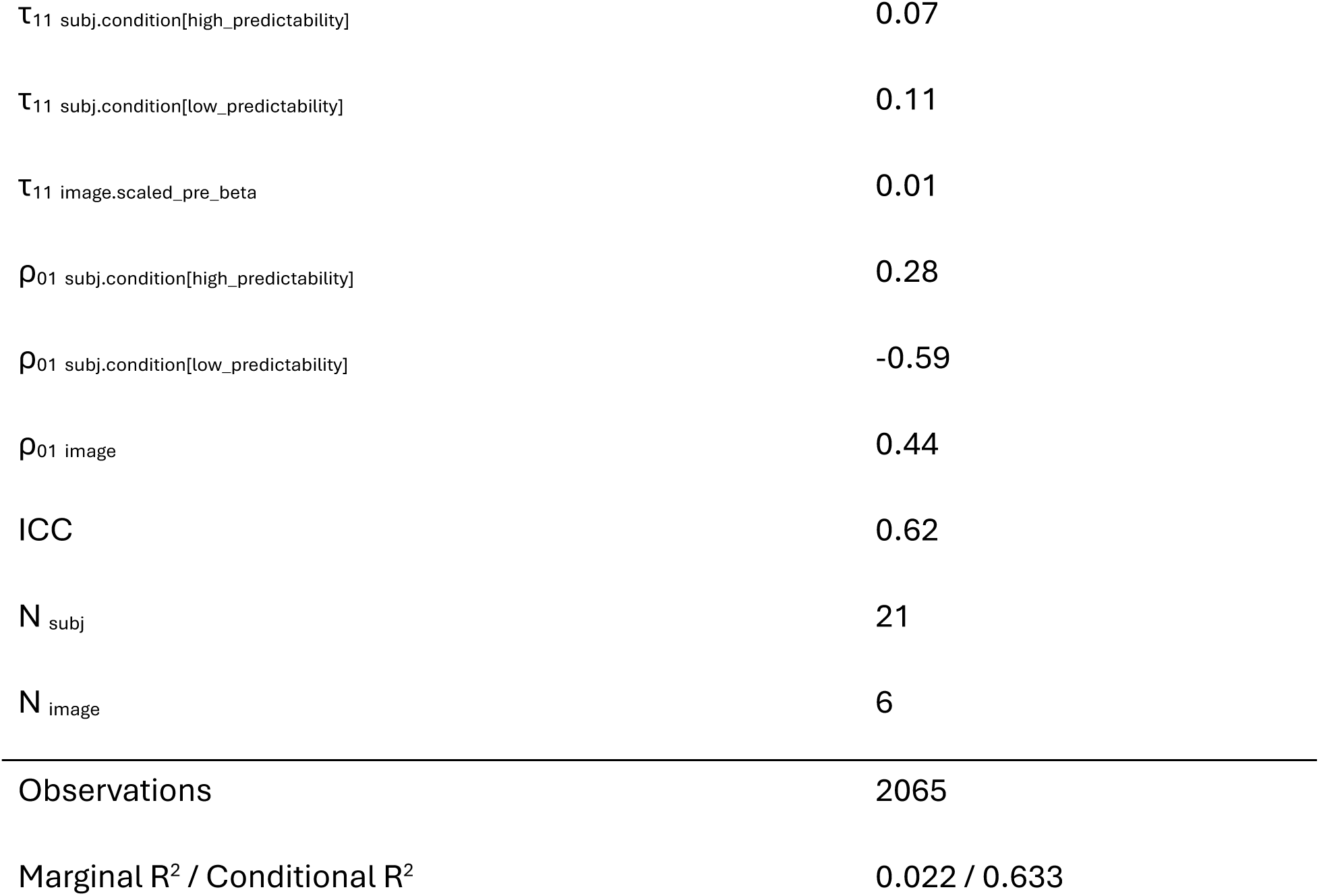

## Appendix C: Model 3 output summary

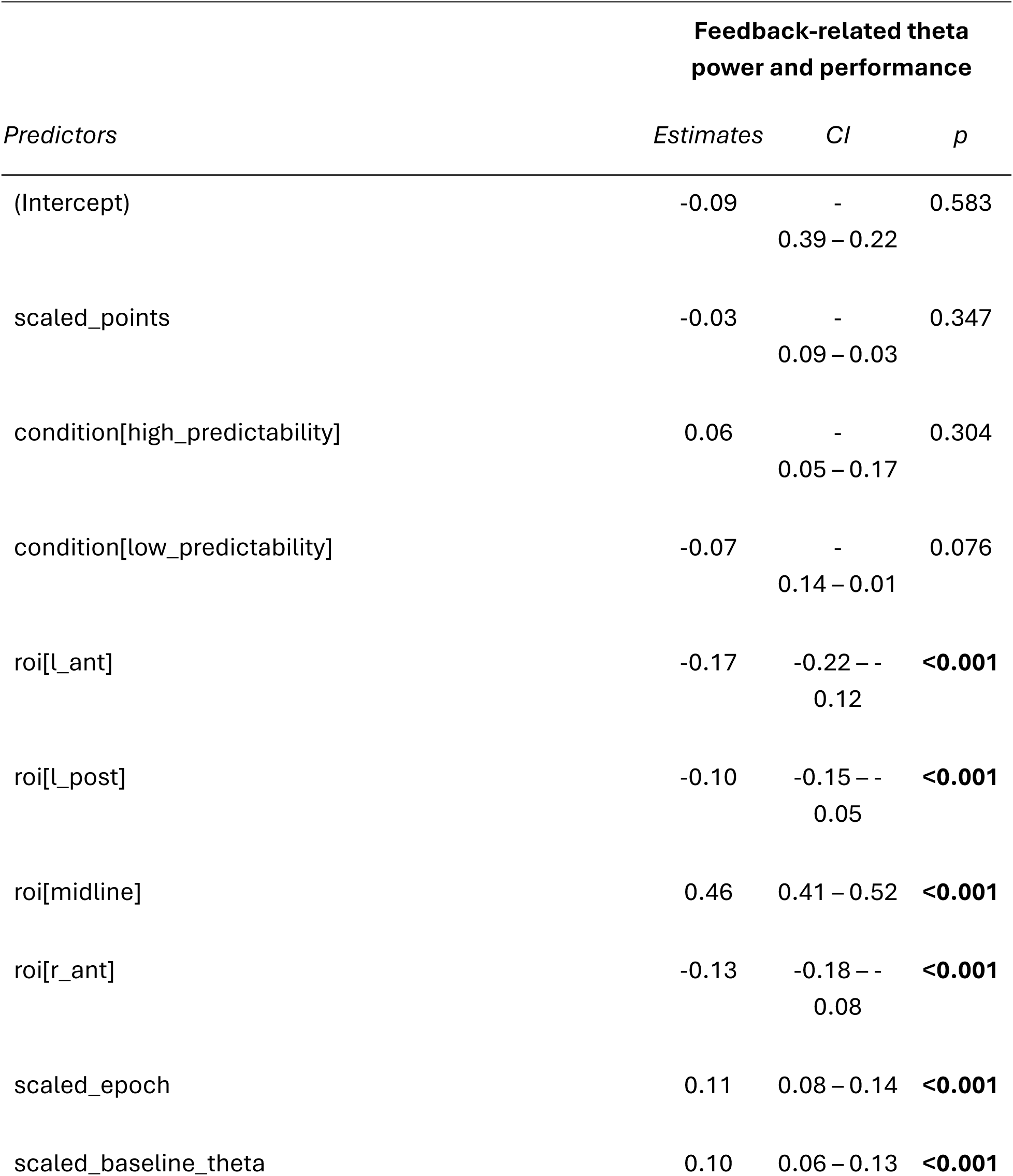
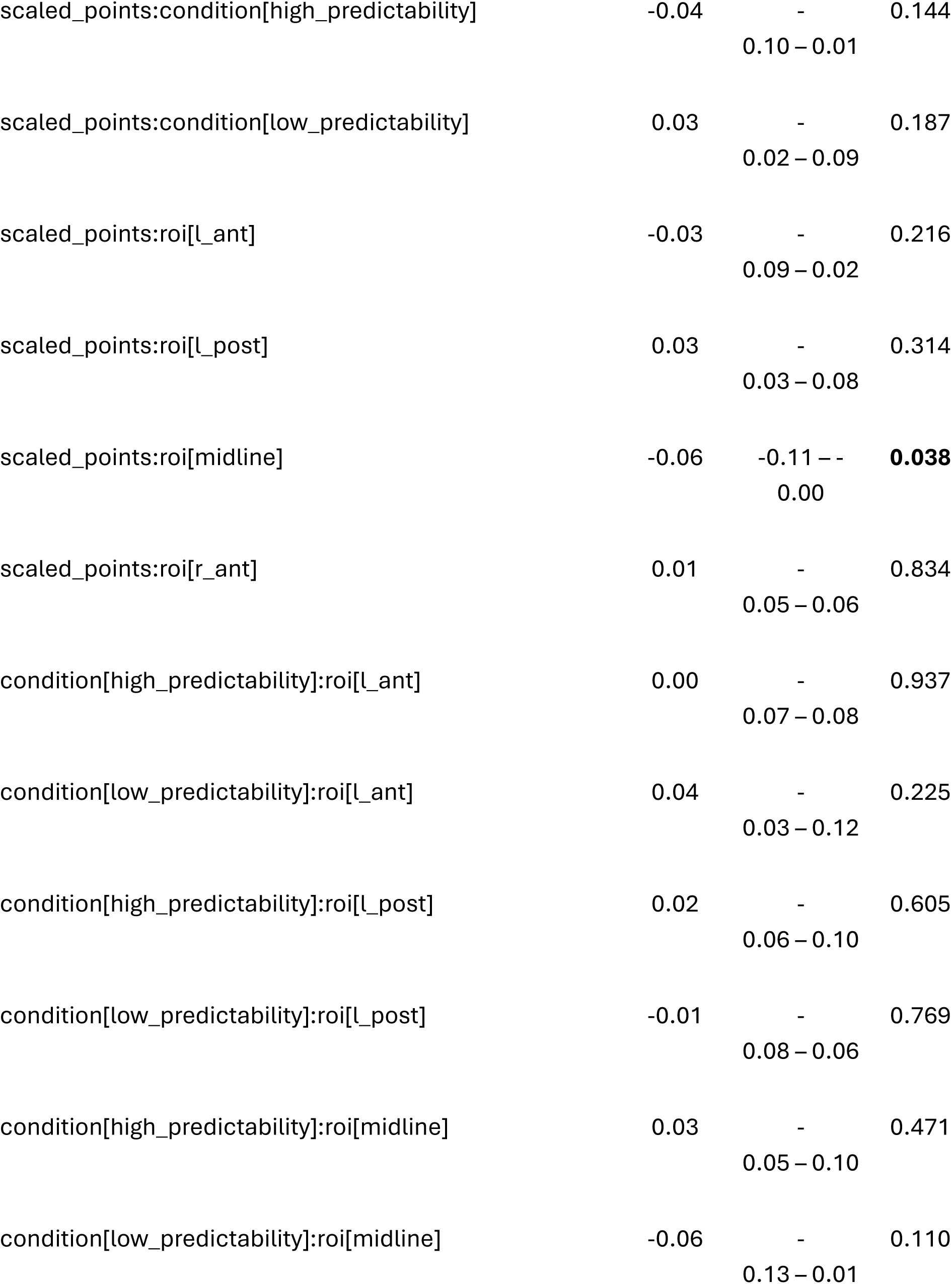
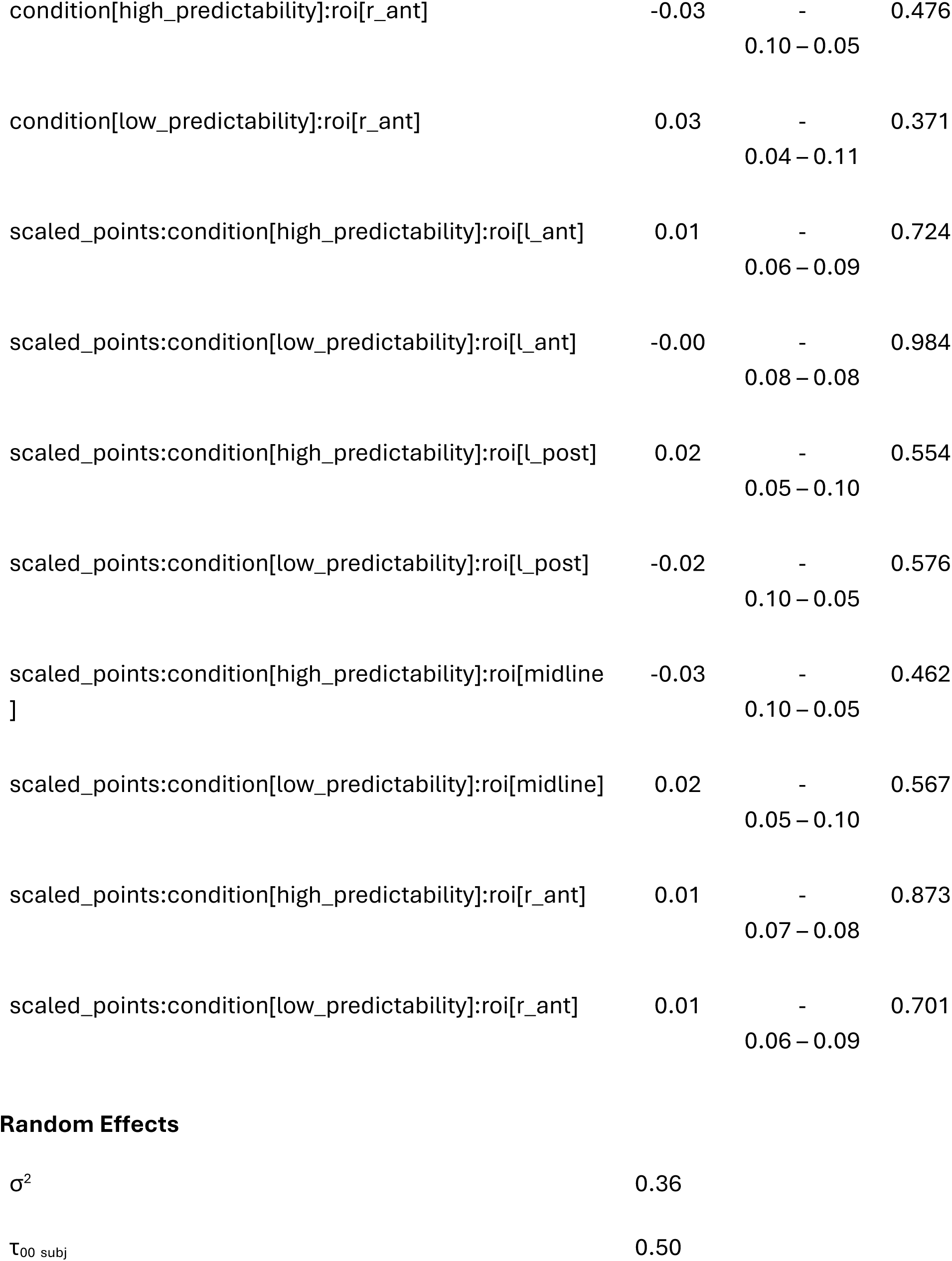
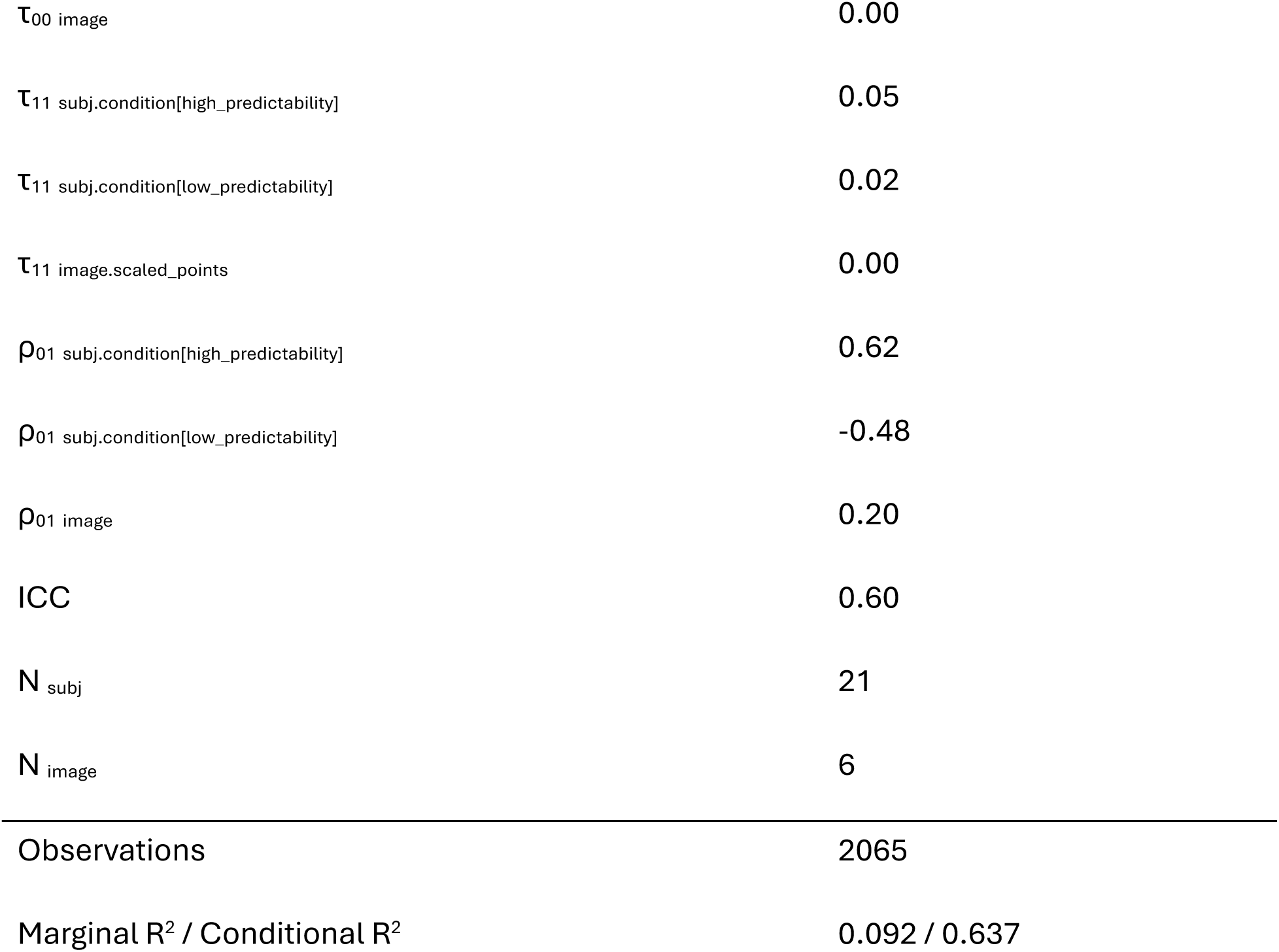

## Appendix D: Model 4 output summary

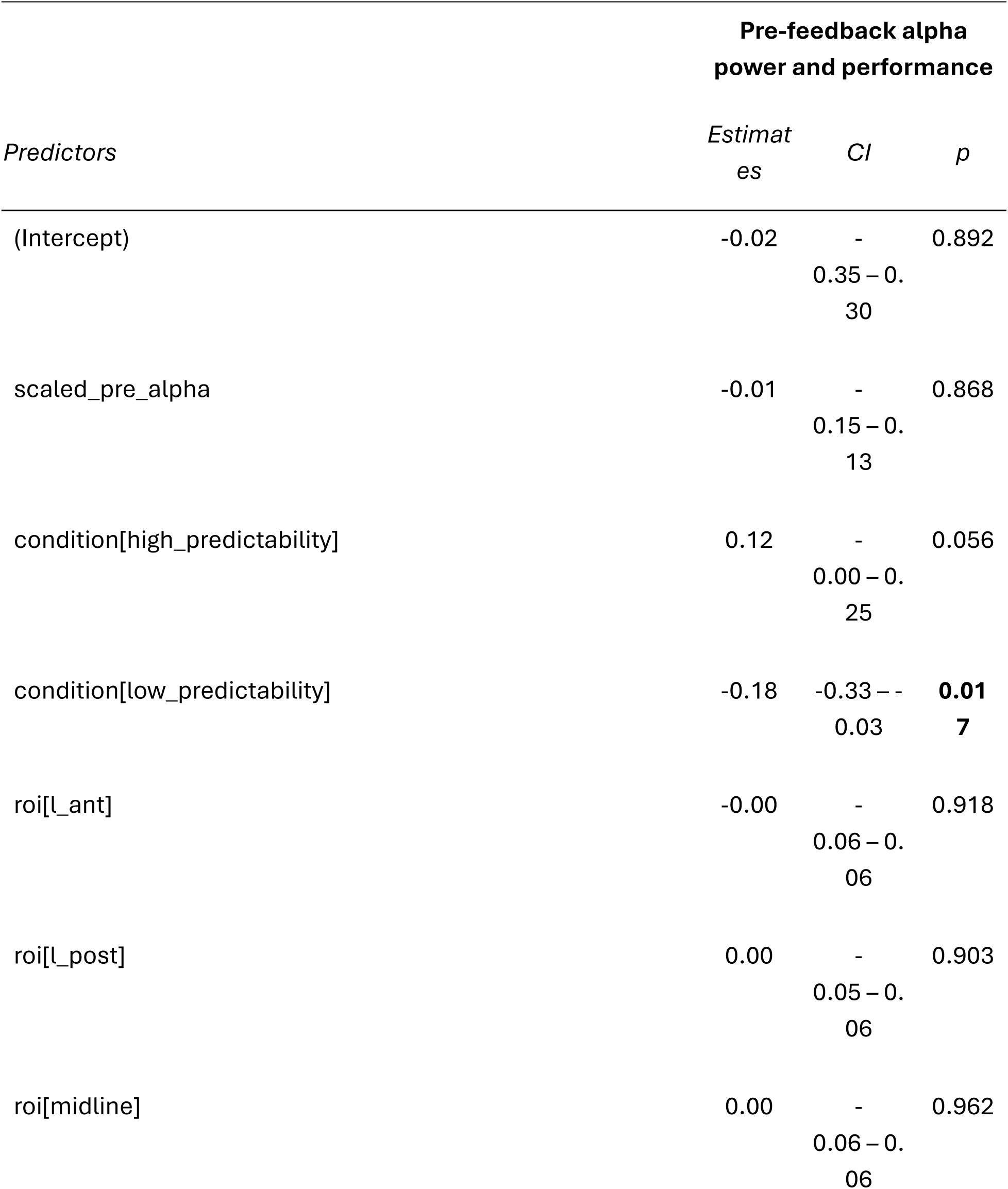
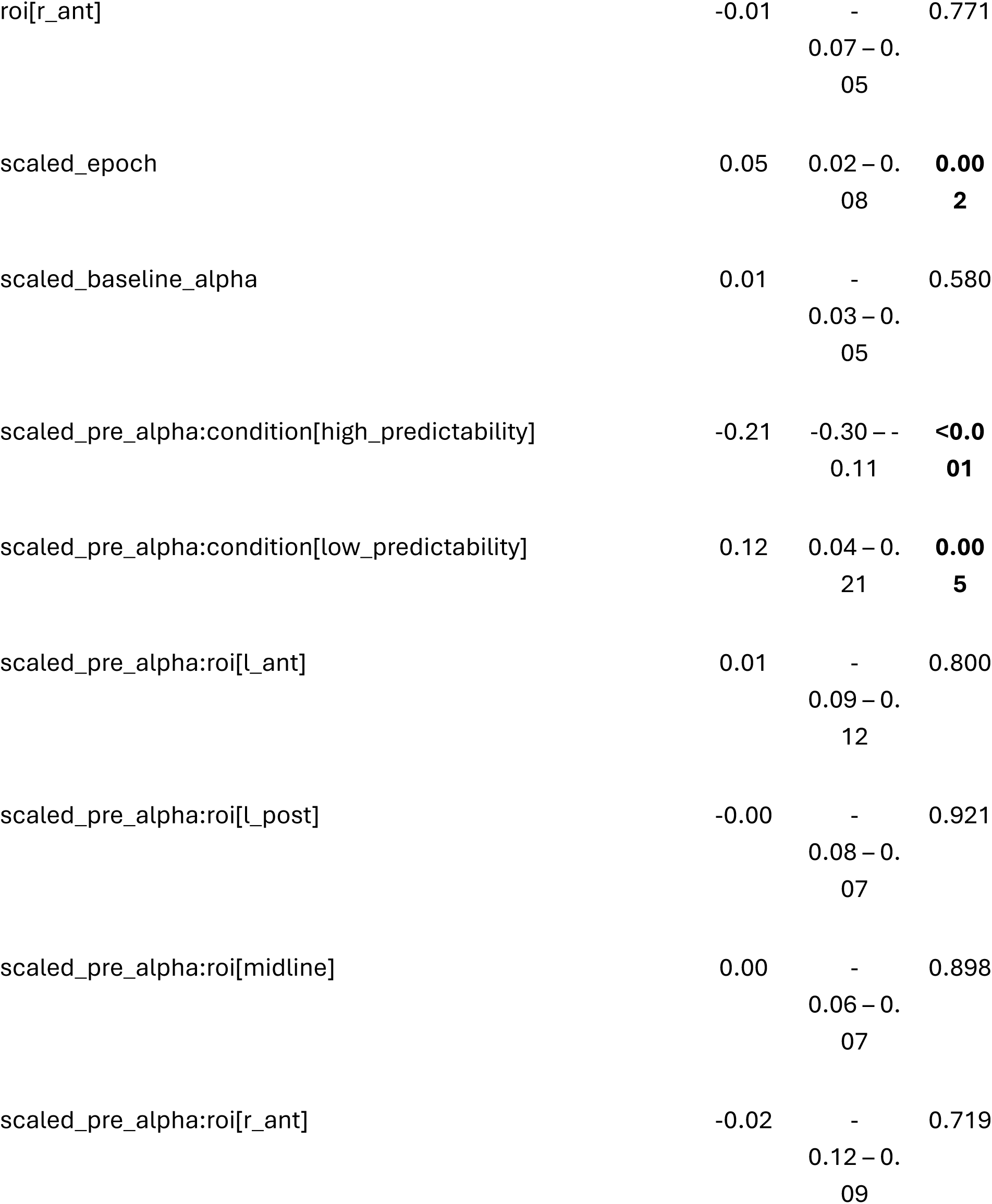
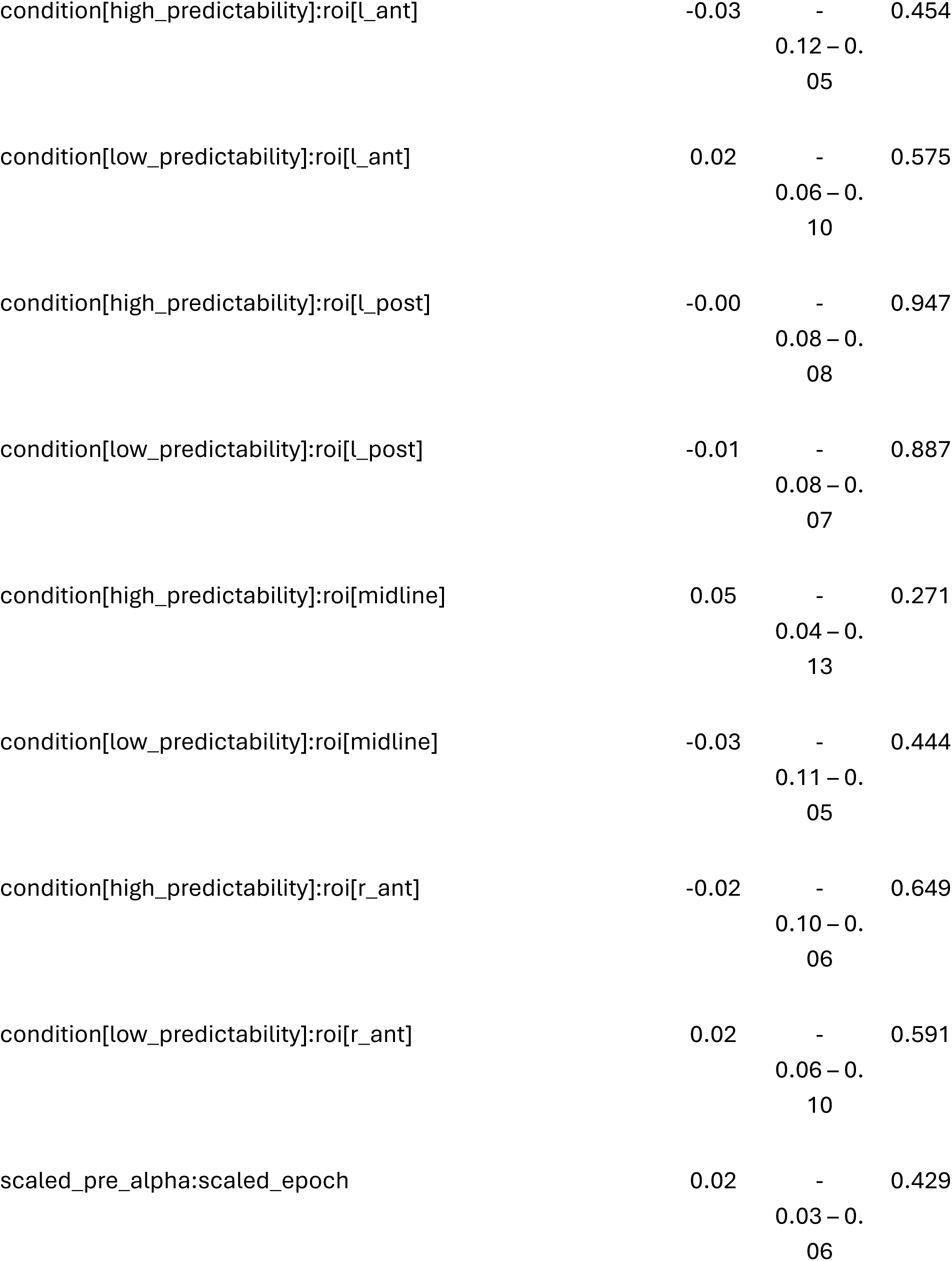
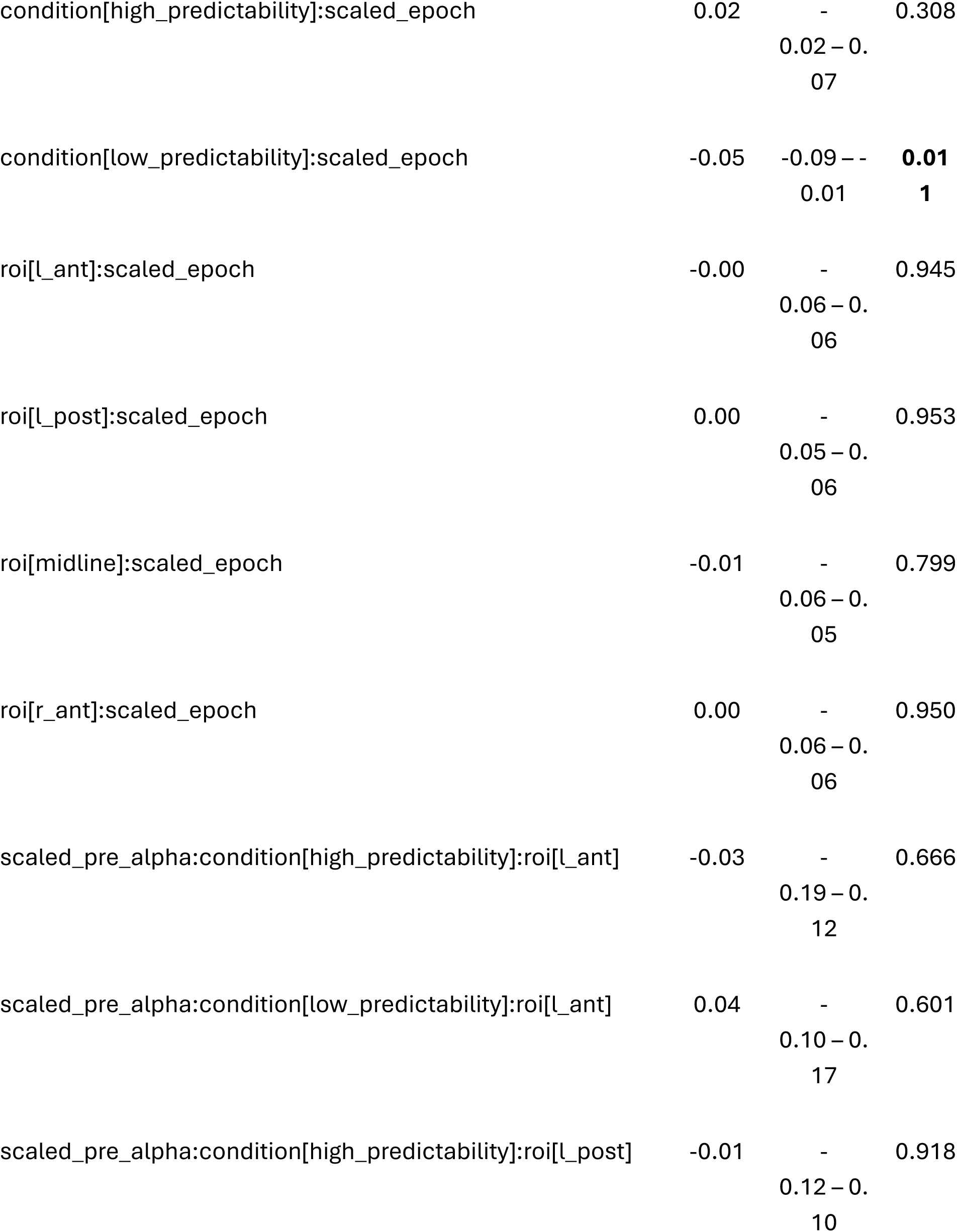
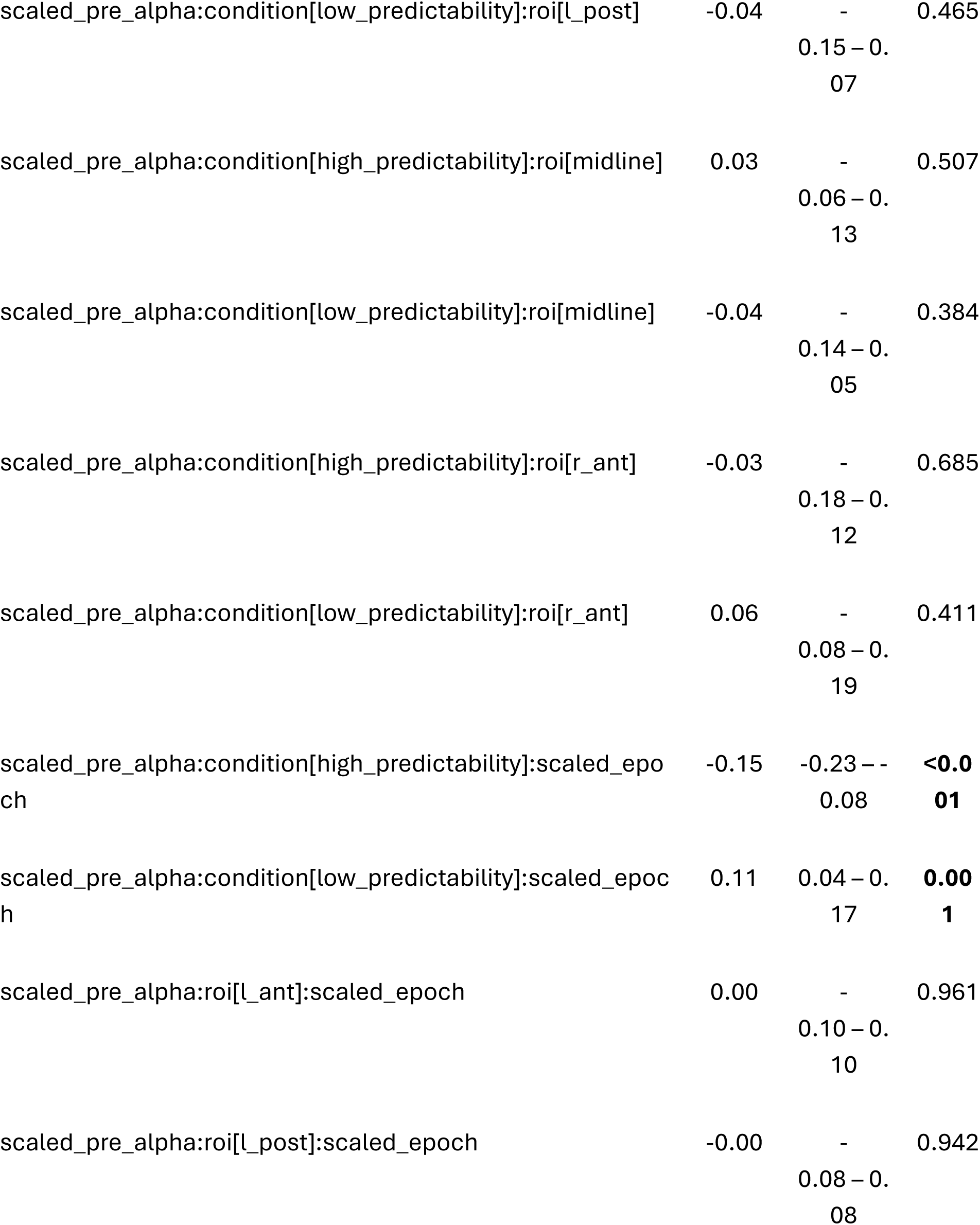
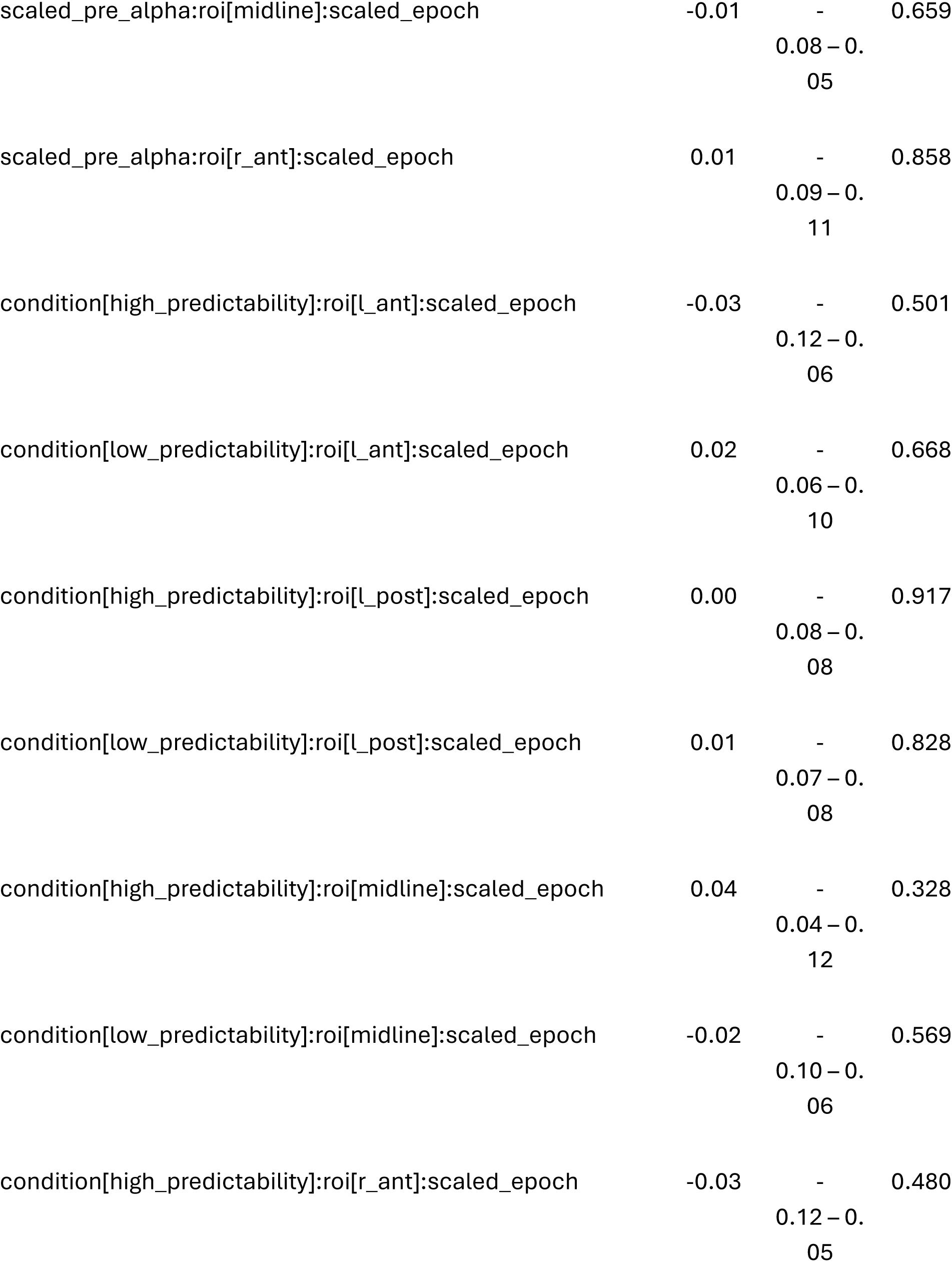
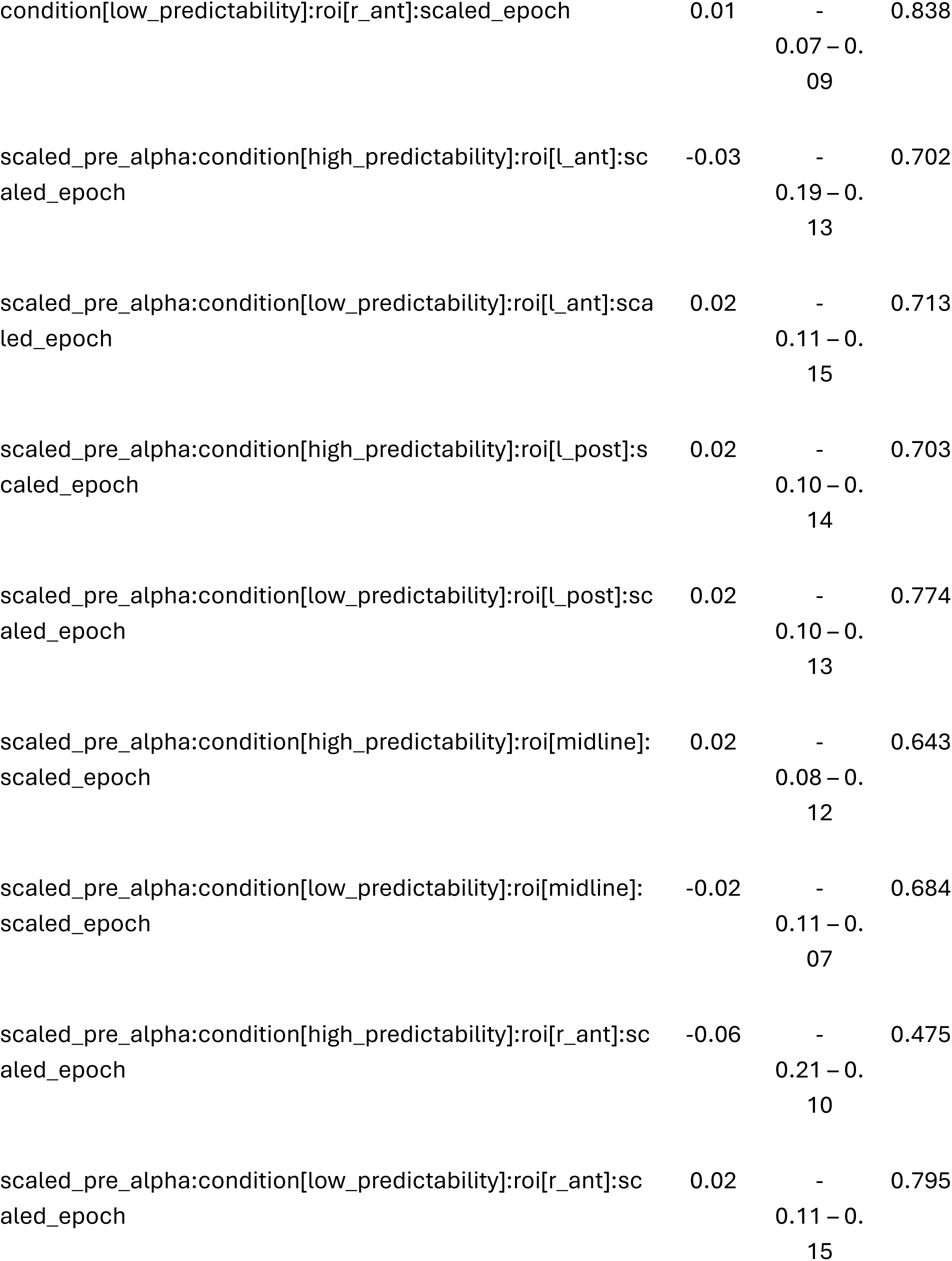
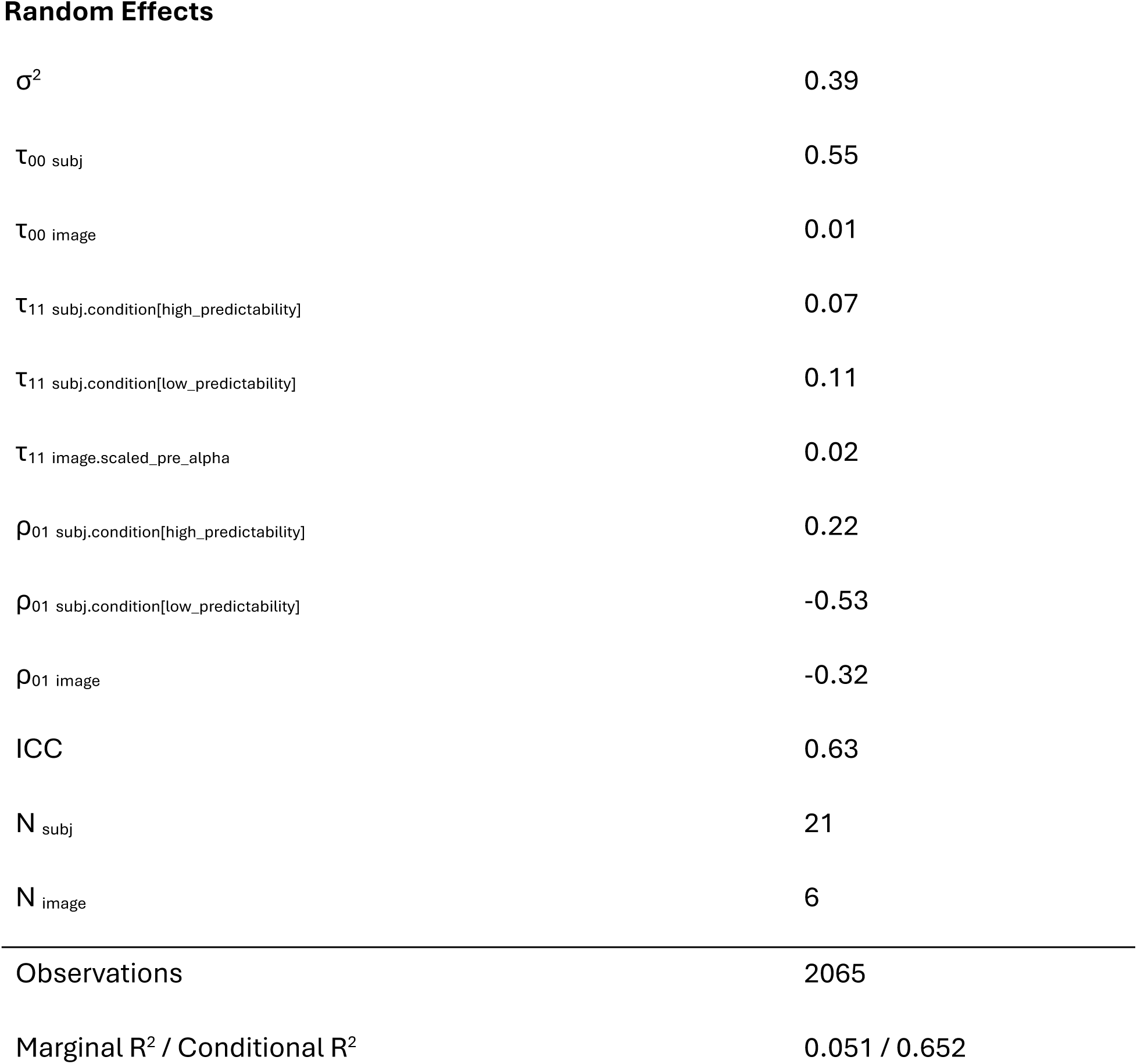

## Appendix E: Model 5 output summary

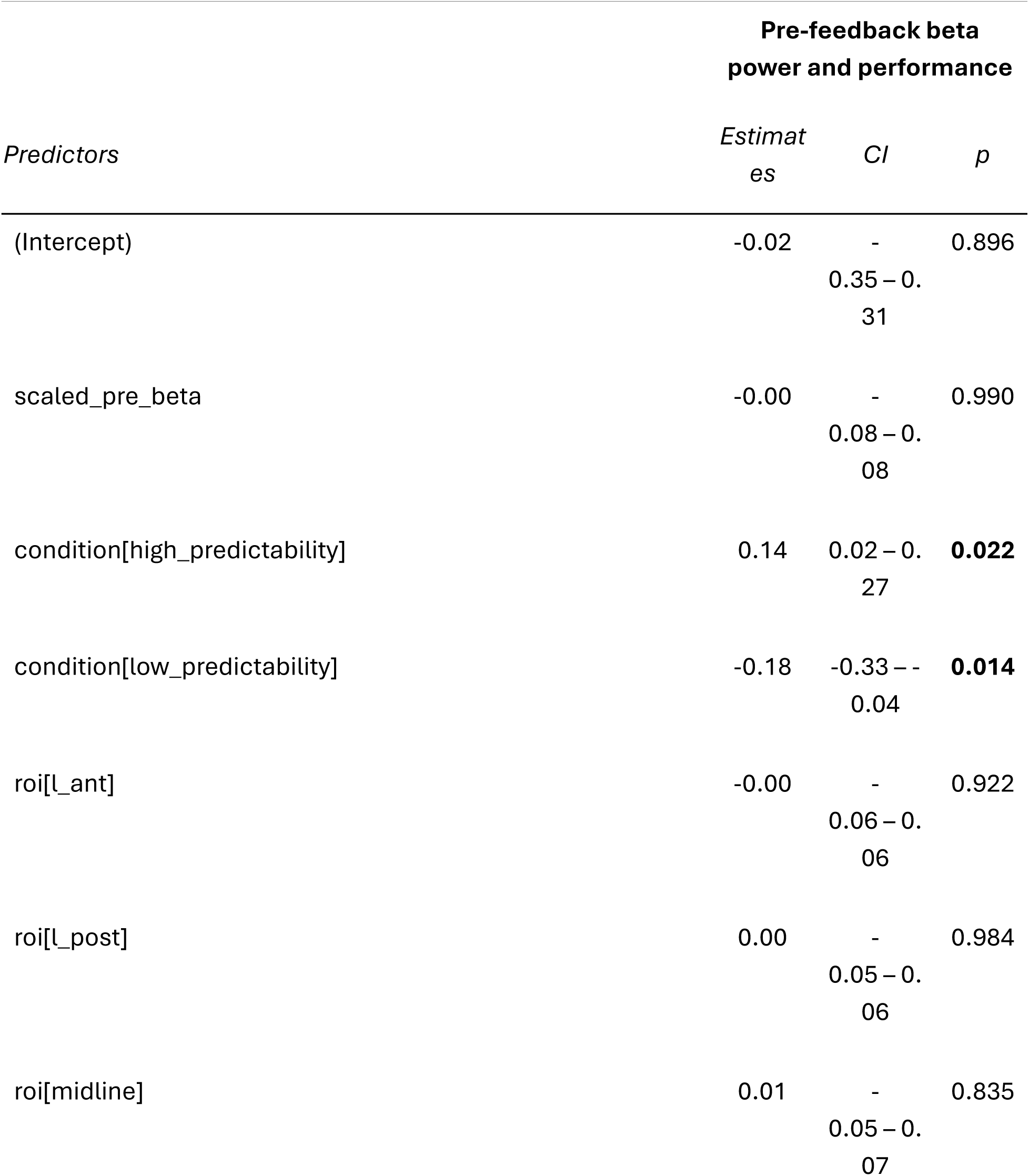
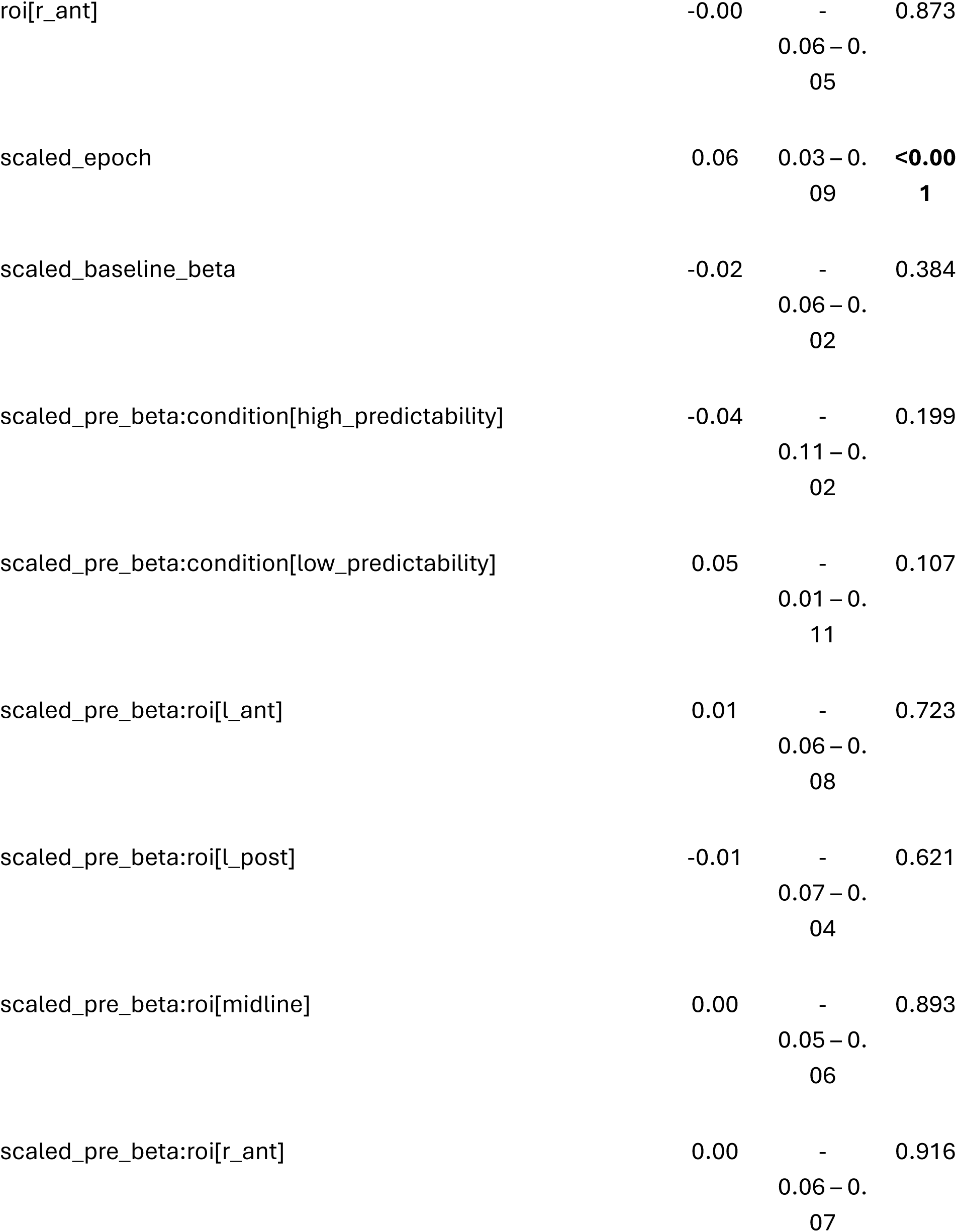
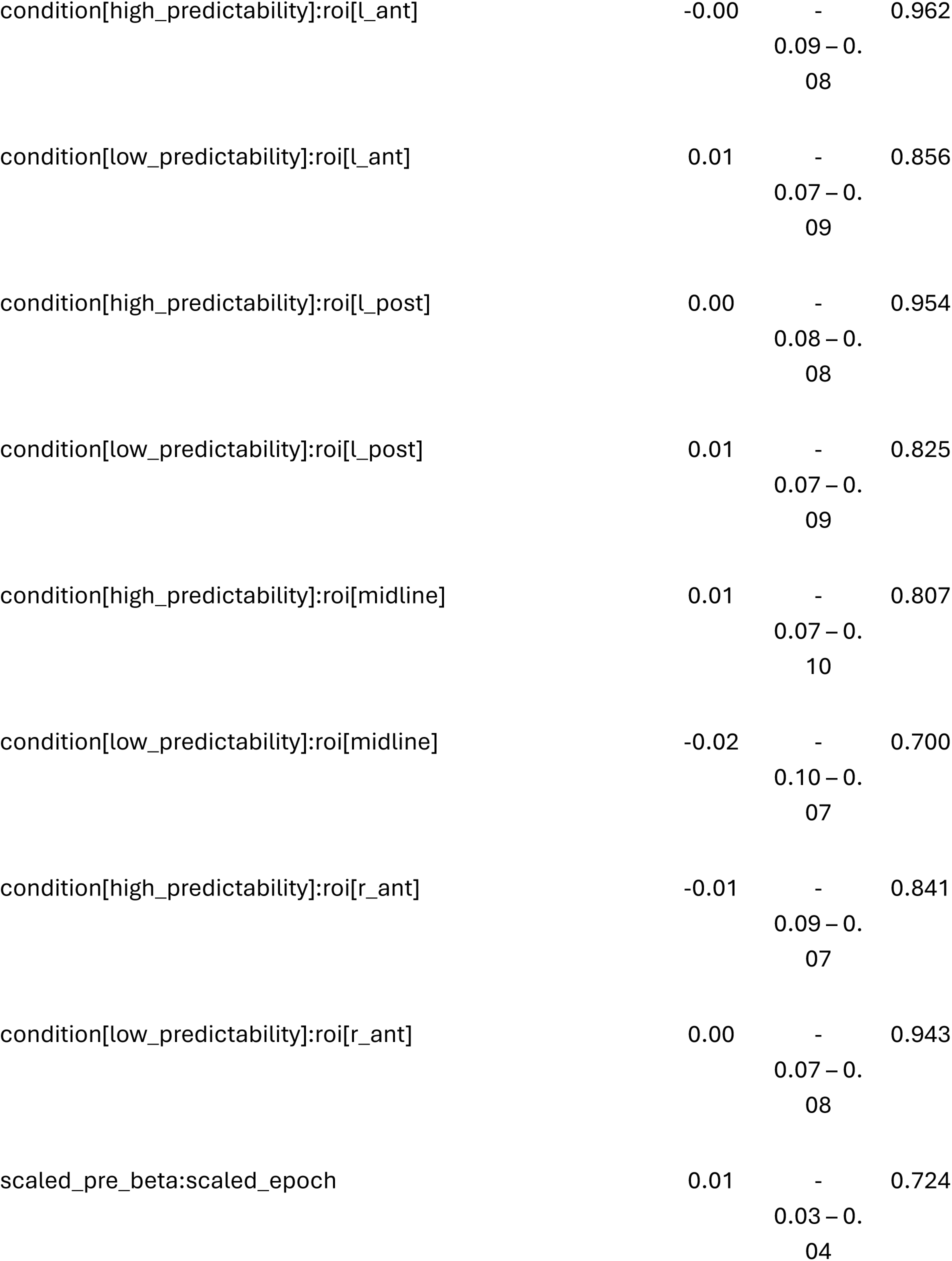
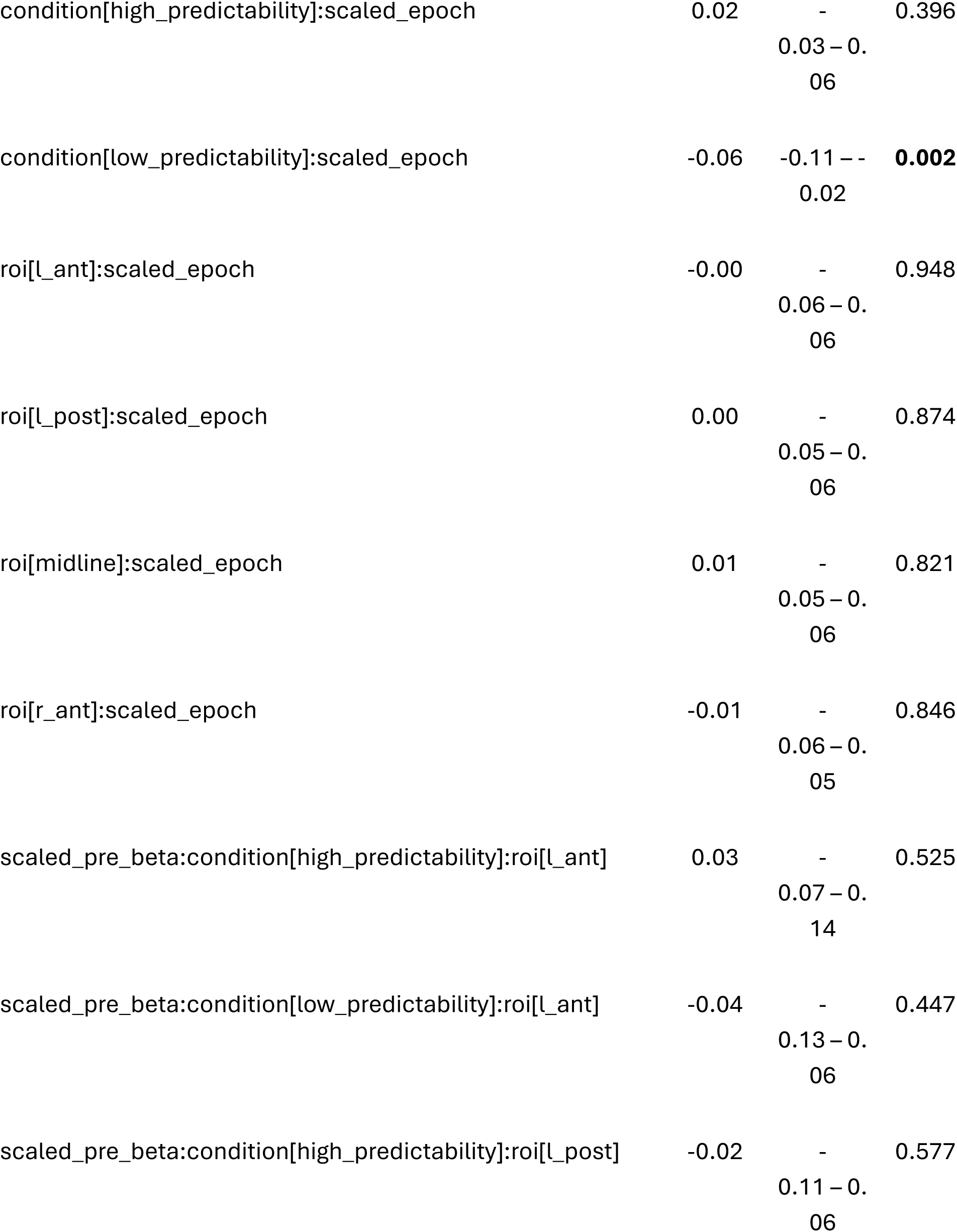
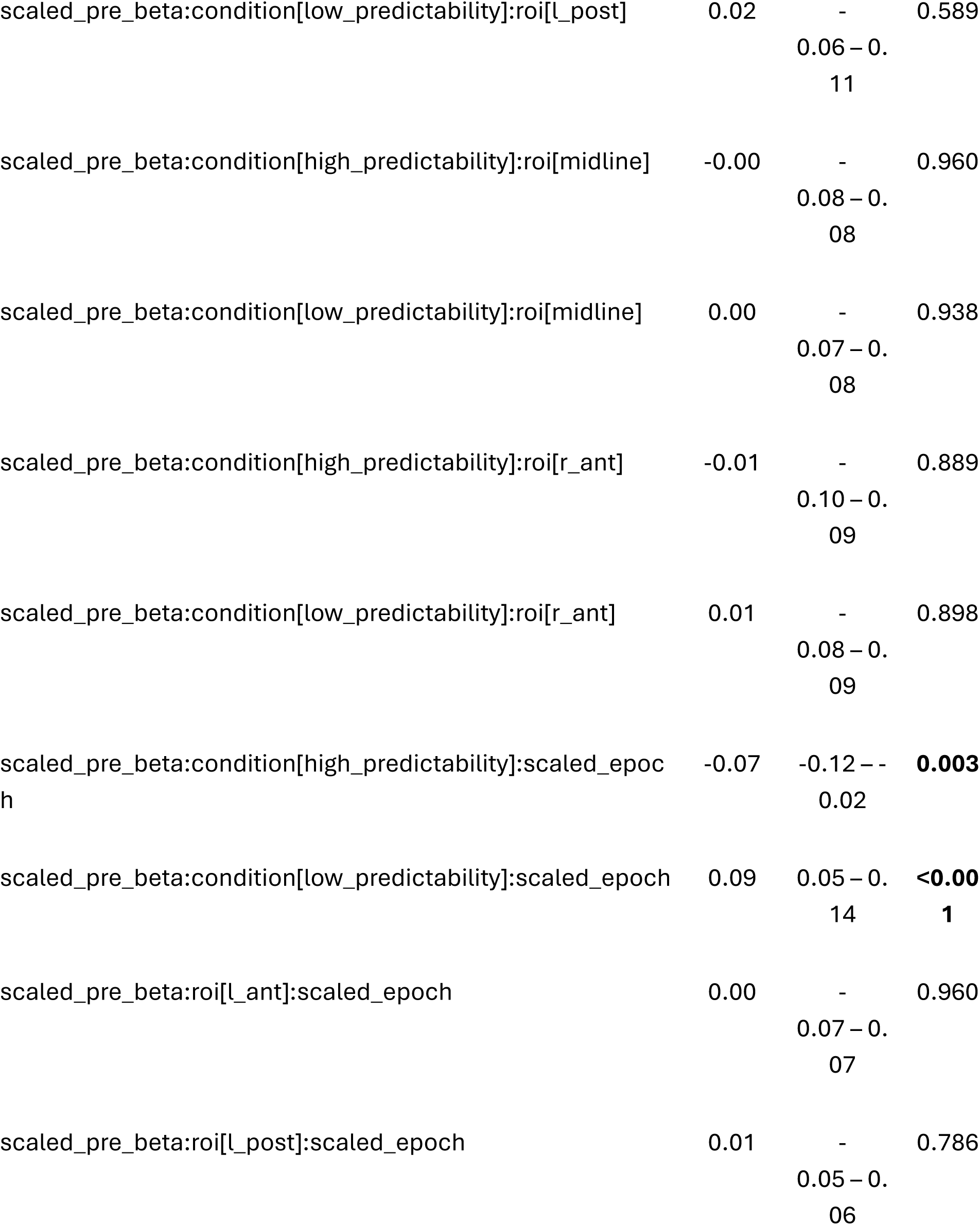
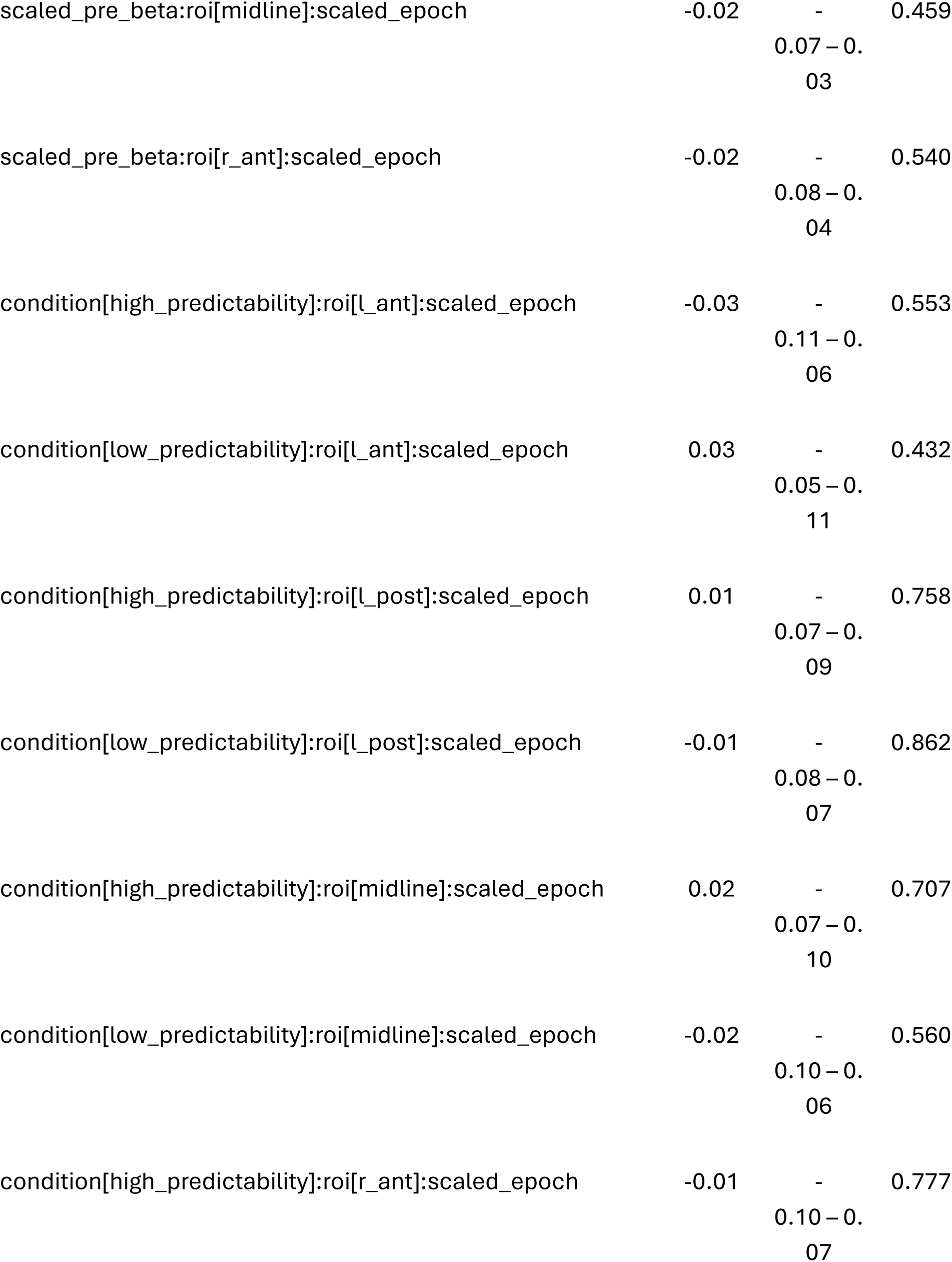
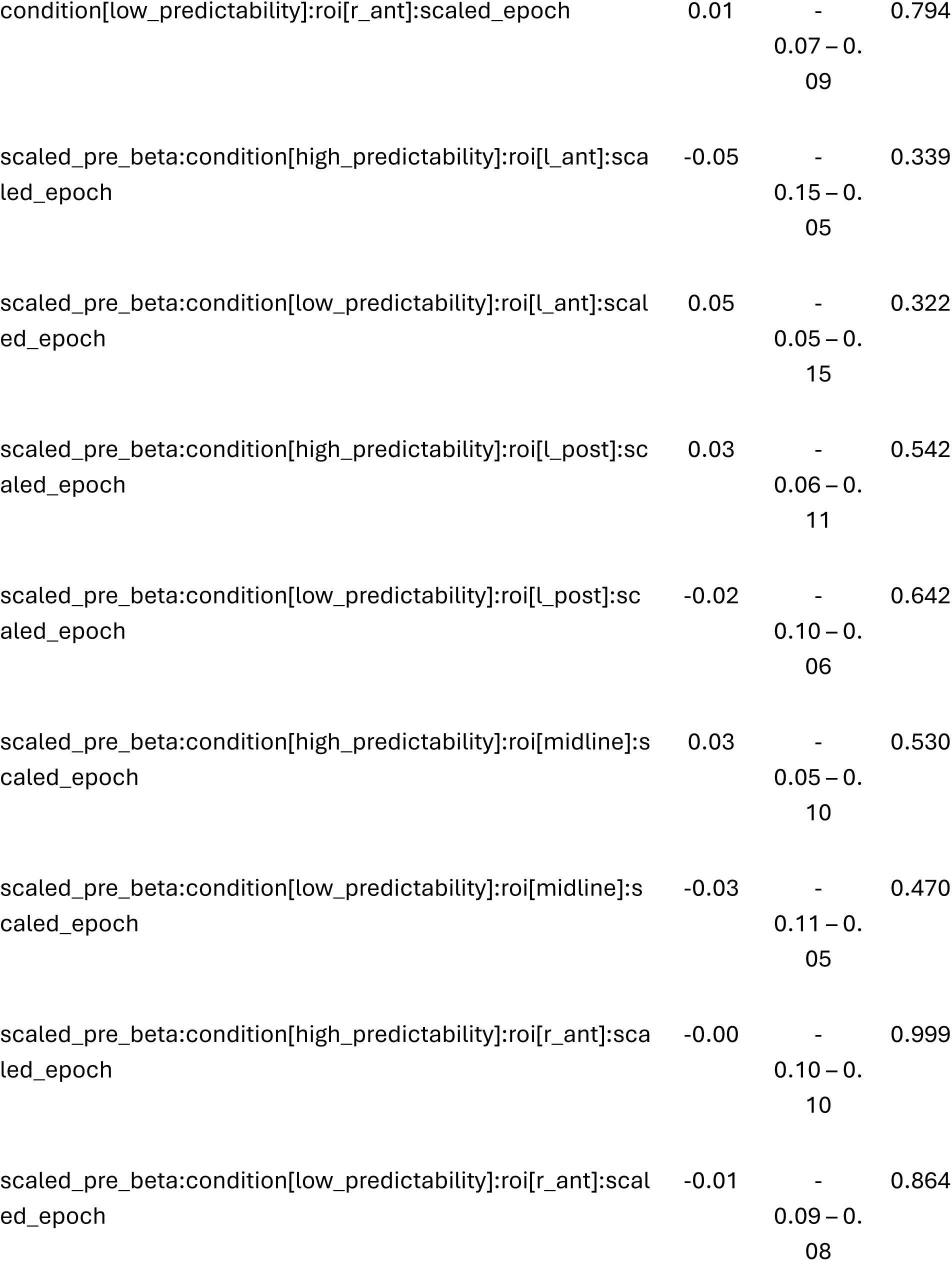
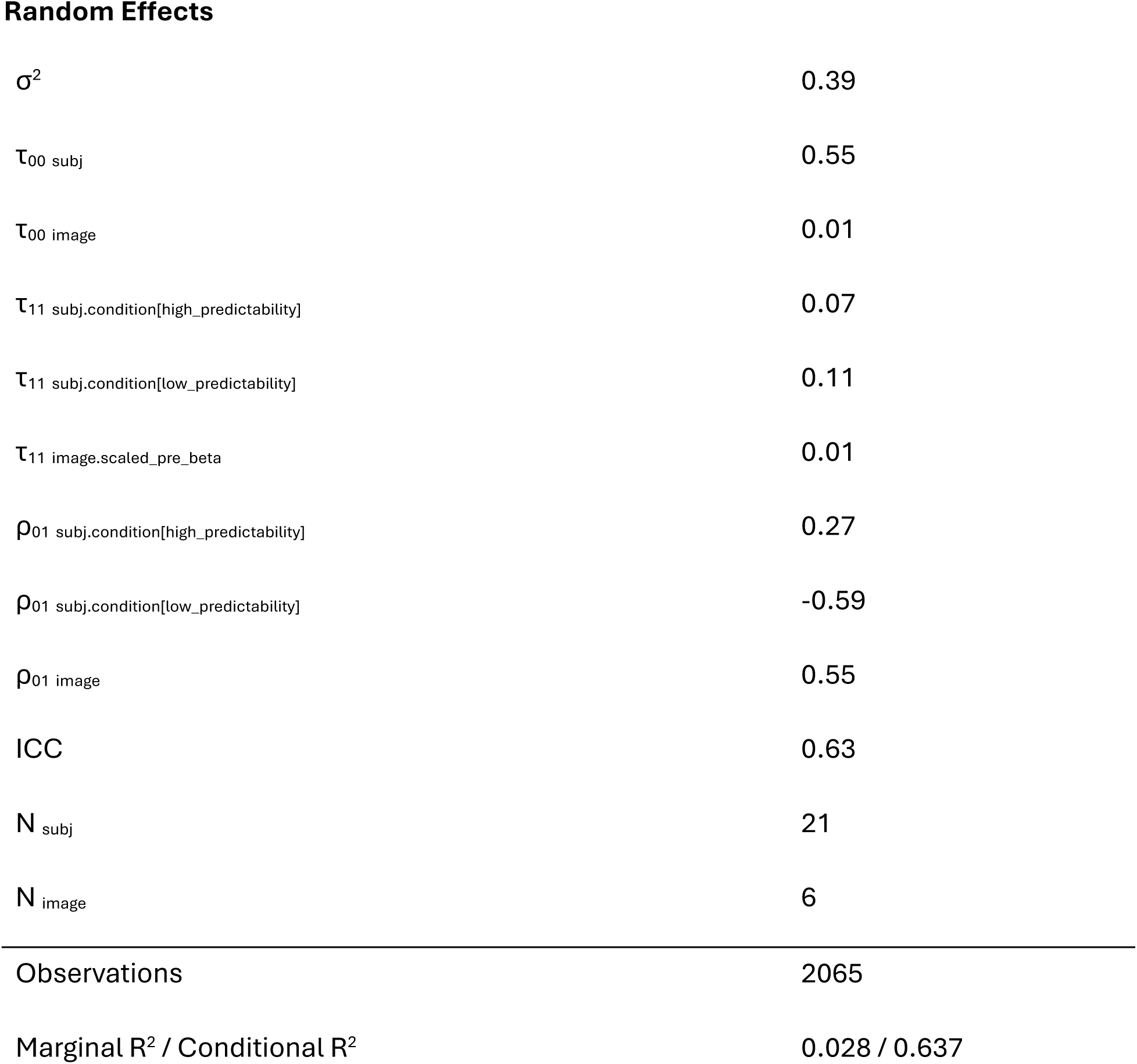

| *Predictors* | Pre-feedback beta<br>power and performance |  |  |
| --- | --- | --- | --- |
|  | *Estimates* | *CI* | *p* |
| (Intercept) | -0.02 | - 0.35 – 0.31 | 0.896 |
| scaled_pre_beta | -0.00 | - 0.08 – 0.08 | 0.990 |
| condition[high_predictability] | 0.14 | 0.02 – 0.27 | **0.022** |
| condition[low_predictability] | -0.18 | -0.33 – -0.04 | **0.014** |
| roi[l_ant] | -0.00 | - 0.06 – 0.06 | 0.922 |
| roi[l_post] | 0.00 | - 0.05 – 0.06 | 0.984 |
| roi[midline] | 0.01 | - 0.05 – 0.07 | 0.835 |
| roi[r_ant] | -0.00 | -<br>0.06 – 0.05 | 0.873 |
| scaled_epoch | 0.06 | 0.03 – 0.09 | **<0.001** |
| scaled_baseline_beta | -0.02 | -<br>0.06 – 0.02 | 0.384 |
| scaled_pre_beta:condition[high_predictability] | -0.04 | -<br>0.11 – 0.02 | 0.199 |
| scaled_pre_beta:condition[low_predictability] | 0.05 | -<br>0.01 – 0.11 | 0.107 |
| scaled_pre_beta:roi[l_ant] | 0.01 | -<br>0.06 – 0.08 | 0.723 |
| scaled_pre_beta:roi[l_post] | -0.01 | -<br>0.07 – 0.04 | 0.621 |
| scaled_pre_beta:roi[midline] | 0.00 | -<br>0.05 – 0.06 | 0.893 |
| scaled_pre_beta:roi[r_ant] | 0.00 | -<br>0.06 – 0.07 | 0.916 |
| condition[high_predictability]:roi[l_ant] | -0.00 | - | 0.962 |
|  |  | 0.09 – 0.08 |  |
| condition[low_predictability]:roi[l_ant] | 0.01 | - | 0.856 |
|  |  | 0.07 – 0.09 |  |
| condition[high_predictability]:roi[l_post] | 0.00 | - | 0.954 |
|  |  | 0.08 – 0.08 |  |
| condition[low_predictability]:roi[l_post] | 0.01 | - | 0.825 |
|  |  | 0.07 – 0.09 |  |
| condition[high_predictability]:roi[midline] | 0.01 | - | 0.807 |
|  |  | 0.07 – 0.10 |  |
| condition[low_predictability]:roi[midline] | -0.02 | - | 0.700 |
|  |  | 0.10 – 0.07 |  |
| condition[high_predictability]:roi[r_ant] | -0.01 | - | 0.841 |
|  |  | 0.09 – 0.07 |  |
| condition[low_predictability]:roi[r_ant] | 0.00 | - | 0.943 |
|  |  | 0.07 – 0.08 |  |
| scaled_pre_beta:scaled_epoch | 0.01 | - | 0.724 |
|  |  | 0.03 – 0.04 |  |
| condition[high_predictability]:scaled_epoch | 0.02 | -<br>0.03 – 0.06 | 0.396 |
| condition[low_predictability]:scaled_epoch | -0.06 | -0.11 – -0.02 | **0.002** |
| roi[l_ant]:scaled_epoch | -0.00 | -<br>0.06 – 0.06 | 0.948 |
| roi[l_post]:scaled_epoch | 0.00 | -<br>0.05 – 0.06 | 0.874 |
| roi[midline]:scaled_epoch | 0.01 | -<br>0.05 – 0.06 | 0.821 |
| roi[r_ant]:scaled_epoch | -0.01 | -<br>0.06 – 0.05 | 0.846 |
| scaled_pre_beta:condition[high_predictability]:roi[l_ant] | 0.03 | -<br>0.07 – 0.14 | 0.525 |
| scaled_pre_beta:condition[low_predictability]:roi[l_ant] | -0.04 | -<br>0.13 – 0.06 | 0.447 |
| scaled_pre_beta:condition[high_predictability]:roi[l_post] | -0.02 | -<br>0.11 – 0.06 | 0.577 |
| scaled_pre_beta:condition[low_predictability]:roi[l_post] | 0.02 | -<br>0.06 – 0.<br>11 | 0.589 |
| scaled_pre_beta:condition[high_predictability]:roi[midline] | -0.00 | -<br>0.08 – 0.<br>08 | 0.960 |
| scaled_pre_beta:condition[low_predictability]:roi[midline] | 0.00 | -<br>0.07 – 0.<br>08 | 0.938 |
| scaled_pre_beta:condition[high_predictability]:roi[r_ant] | -0.01 | -<br>0.10 – 0.<br>09 | 0.889 |
| scaled_pre_beta:condition[low_predictability]:roi[r_ant] | 0.01 | -<br>0.08 – 0.<br>09 | 0.898 |
| scaled_pre_beta:condition[high_predictability]:scaled_epoch | -0.07 | -0.12 – -<br>0.02 | **0.003** |
| scaled_pre_beta:condition[low_predictability]:scaled_epoch | 0.09 | 0.05 – 0.<br>14 | **<0.001** |
| scaled_pre_beta:roi[l_ant]:scaled_epoch | 0.00 | -<br>0.07 – 0.<br>07 | 0.960 |
| scaled_pre_beta:roi[l_post]:scaled_epoch | 0.01 | -<br>0.05 – 0.<br>06 | 0.786 |
| scaled_pre_beta:roi[midline]:scaled_epoch | -0.02 | -<br>0.07 – 0.<br>03 | 0.459 |
| scaled_pre_beta:roi[r_ant]:scaled_epoch | -0.02 | -<br>0.08 – 0.<br>04 | 0.540 |
| condition[high_predictability]:roi[l_ant]:scaled_epoch | -0.03 | -<br>0.11 – 0.<br>06 | 0.553 |
| condition[low_predictability]:roi[l_ant]:scaled_epoch | 0.03 | -<br>0.05 – 0.<br>11 | 0.432 |
| condition[high_predictability]:roi[l_post]:scaled_epoch | 0.01 | -<br>0.07 – 0.<br>09 | 0.758 |
| condition[low_predictability]:roi[l_post]:scaled_epoch | -0.01 | -<br>0.08 – 0.<br>07 | 0.862 |
| condition[high_predictability]:roi[midline]:scaled_epoch | 0.02 | -<br>0.07 – 0.<br>10 | 0.707 |
| condition[low_predictability]:roi[midline]:scaled_epoch | -0.02 | -<br>0.10 – 0.<br>06 | 0.560 |
| condition[high_predictability]:roi[r_ant]:scaled_epoch | -0.01 | -<br>0.10 – 0.<br>07 | 0.777 |
| condition[low_predictability]:roi[r_ant]:scaled_epoch | 0.01 | -<br>0.07 – 0.09 | 0.794 |
| scaled_pre_beta:condition[high_predictability]:roi[l_ant]:scaled_epoch | -0.05 | -<br>0.15 – 0.05 | 0.339 |
| scaled_pre_beta:condition[low_predictability]:roi[l_ant]:scaled_epoch | 0.05 | -<br>0.05 – 0.15 | 0.322 |
| scaled_pre_beta:condition[high_predictability]:roi[l_post]:scaled_epoch | 0.03 | -<br>0.06 – 0.11 | 0.542 |
| scaled_pre_beta:condition[low_predictability]:roi[l_post]:scaled_epoch | -0.02 | -<br>0.10 – 0.06 | 0.642 |
| scaled_pre_beta:condition[high_predictability]:roi[midline]:scaled_epoch | 0.03 | -<br>0.05 – 0.10 | 0.530 |
| scaled_pre_beta:condition[low_predictability]:roi[midline]:scaled_epoch | -0.03 | -<br>0.11 – 0.05 | 0.470 |
| scaled_pre_beta:condition[high_predictability]:roi[r_ant]:scaled_epoch | -0.00 | -<br>0.10 – 0.10 | 0.999 |
| scaled_pre_beta:condition[low_predictability]:roi[r_ant]:scaled_epoch | -0.01 | -<br>0.09 – 0.08 | 0.864 |

Random Effects
|  |  |
| --- | --- |
| $\sigma^2$ | 0.39 |
| $\tau_{00}$ subj | 0.55 |
| $\tau_{00}$ image | 0.01 |
| $\tau_{11}$ subj.condition[high_predictability] | 0.07 |
| $\tau_{11}$ subj.condition[low_predictability] | 0.11 |
| $\tau_{11}$ image.scaled_pre_beta | 0.01 |
| $\rho_{01}$ subj.condition[high_predictability] | 0.27 |
| $\rho_{01}$ subj.condition[low_predictability] | -0.59 |
| $\rho_{01}$ image | 0.55 |
| ICC | 0.63 |
| $N_{\text{subj}}$ | 21 |
| $N_{\text{image}}$ | 6 |
| Observations | 2065 |
| Marginal $R^2$ / Conditional $R^2$ | 0.028 / 0.637 |

## Appendix F: pre-feedback theta power model output

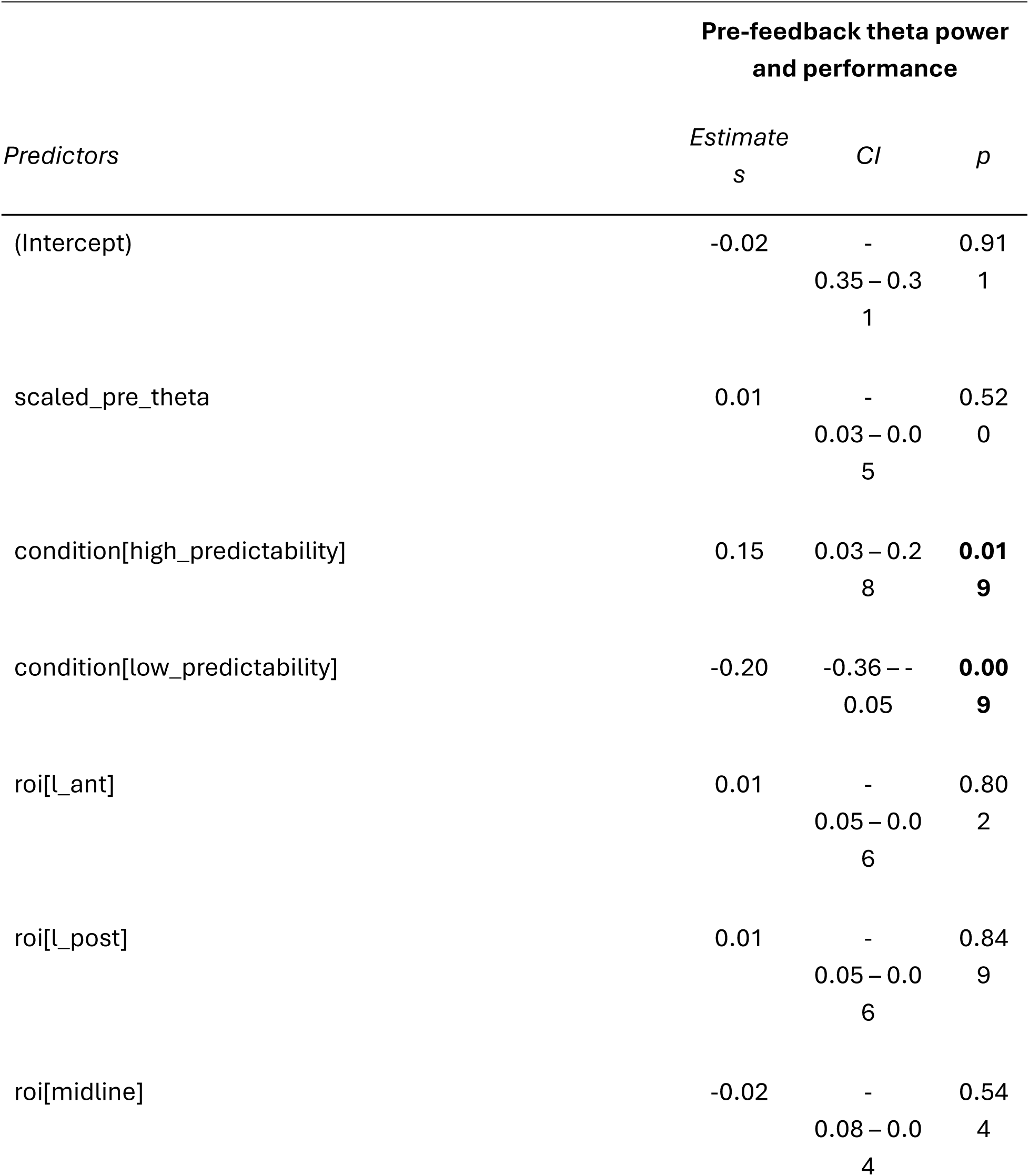
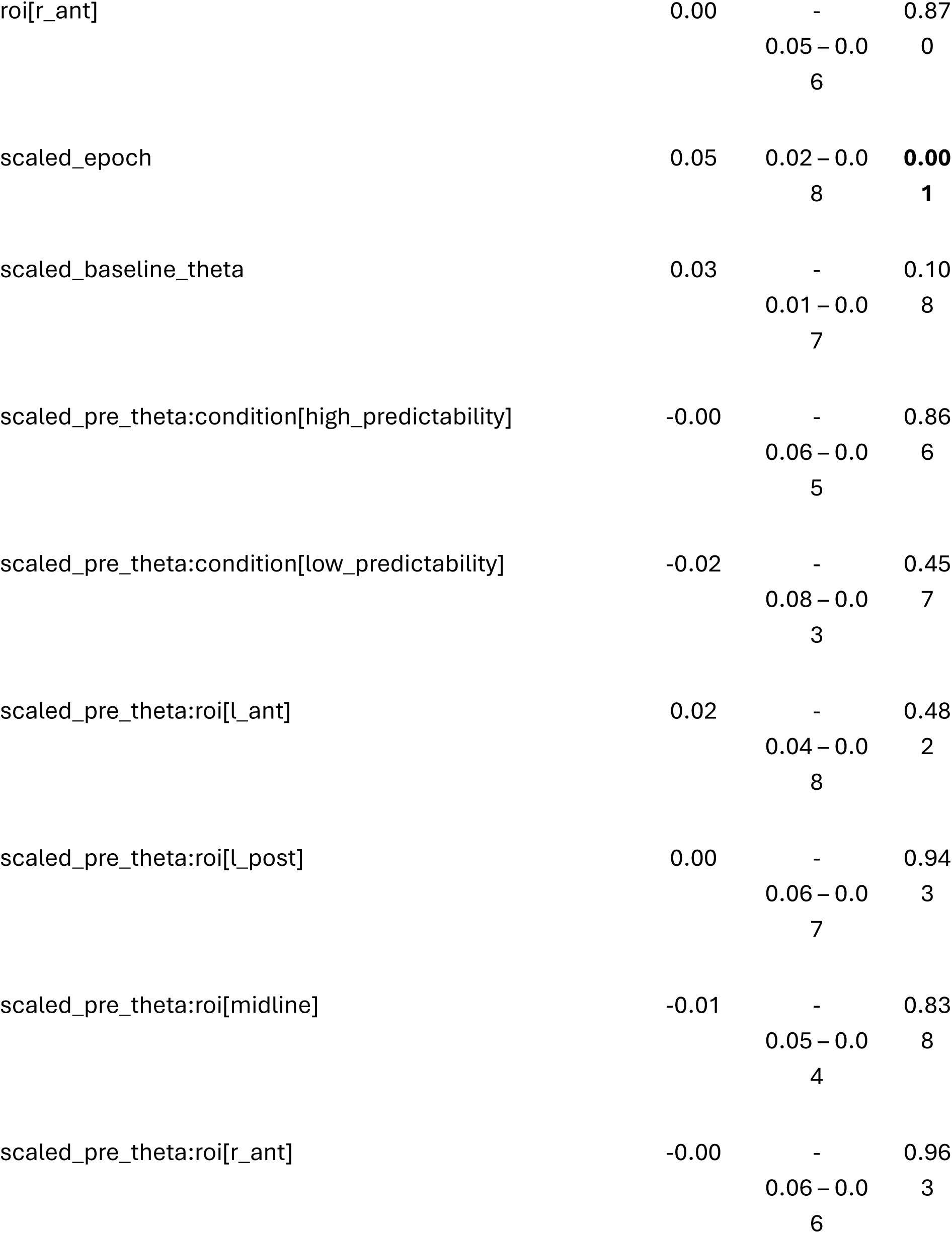
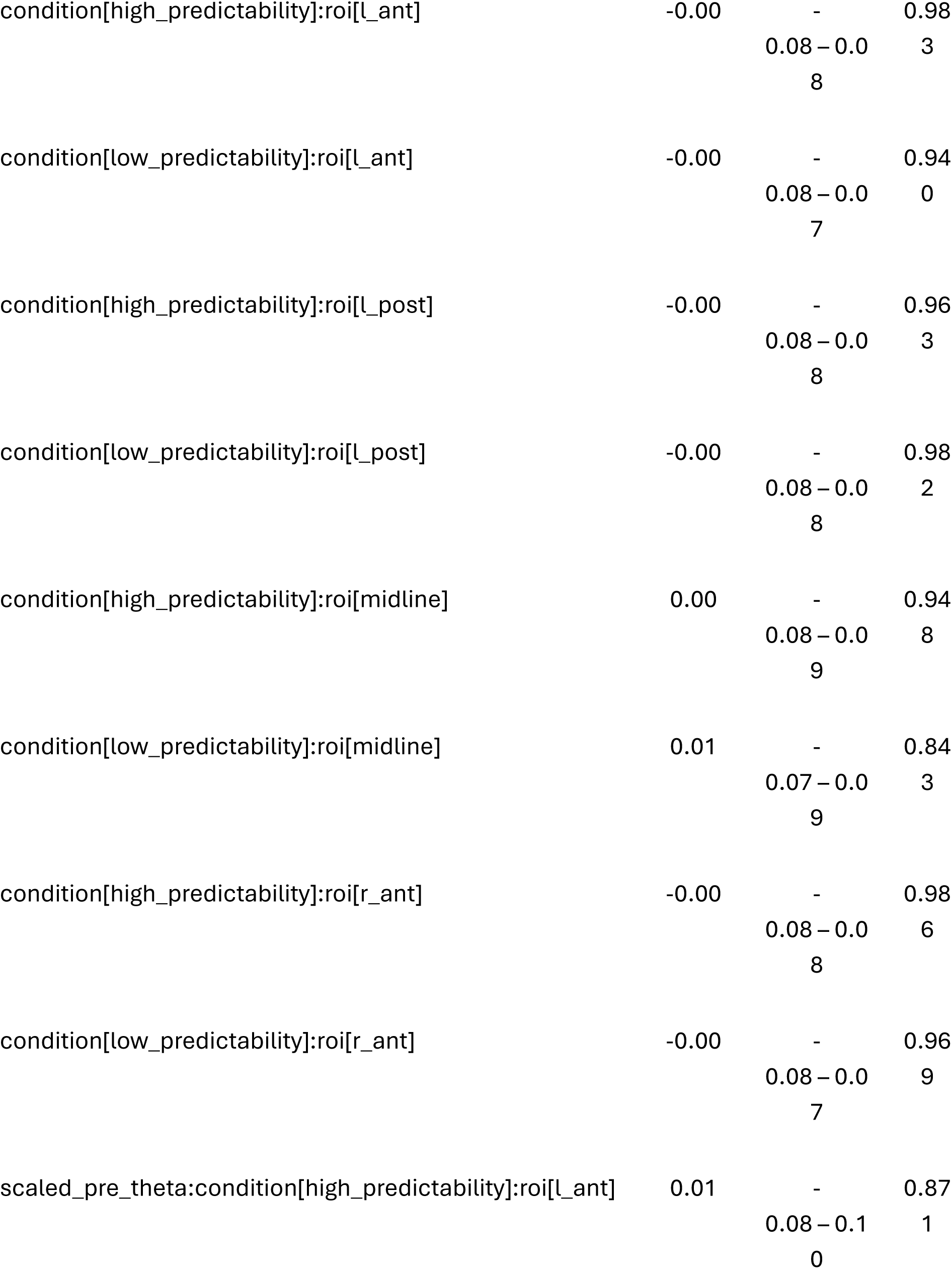
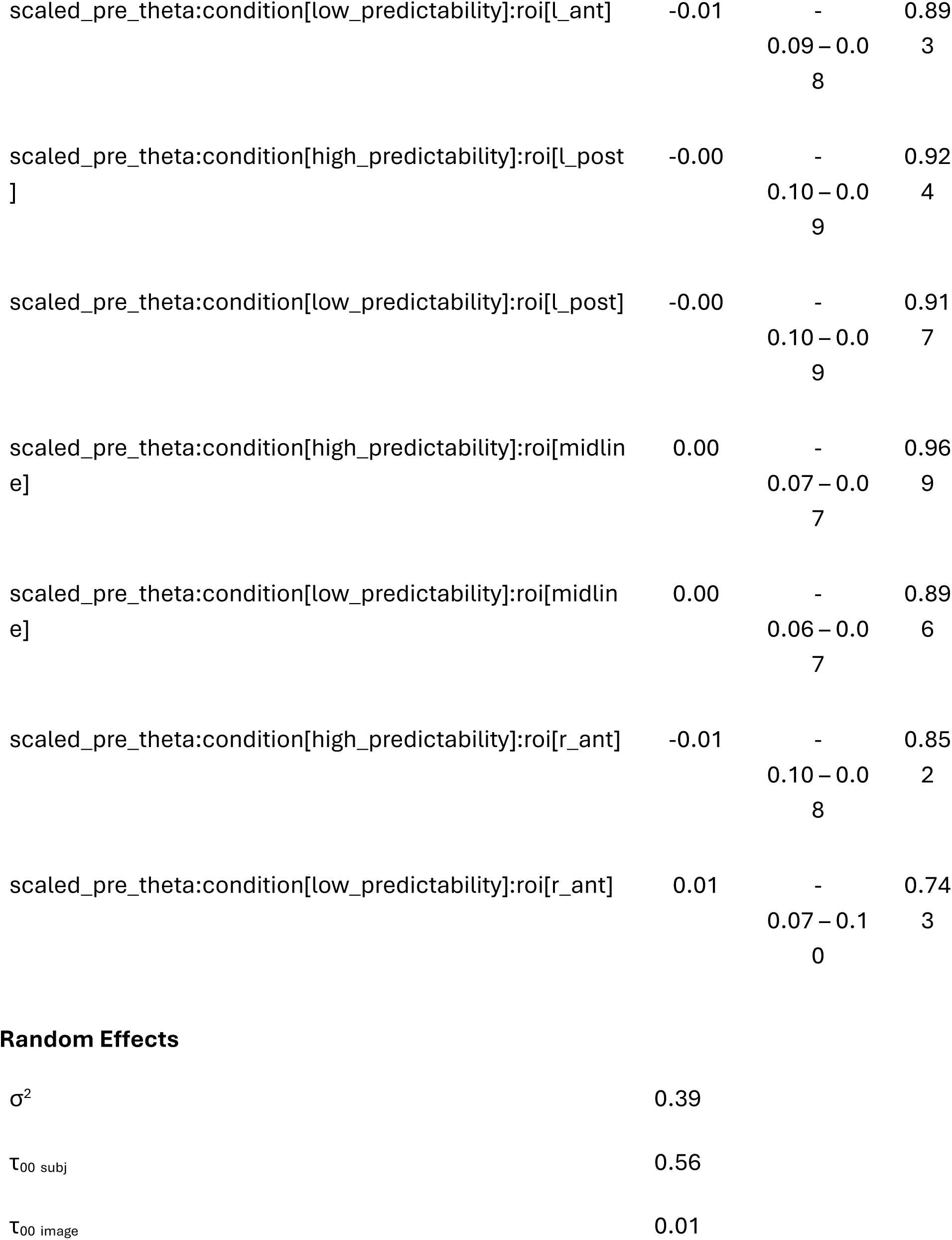
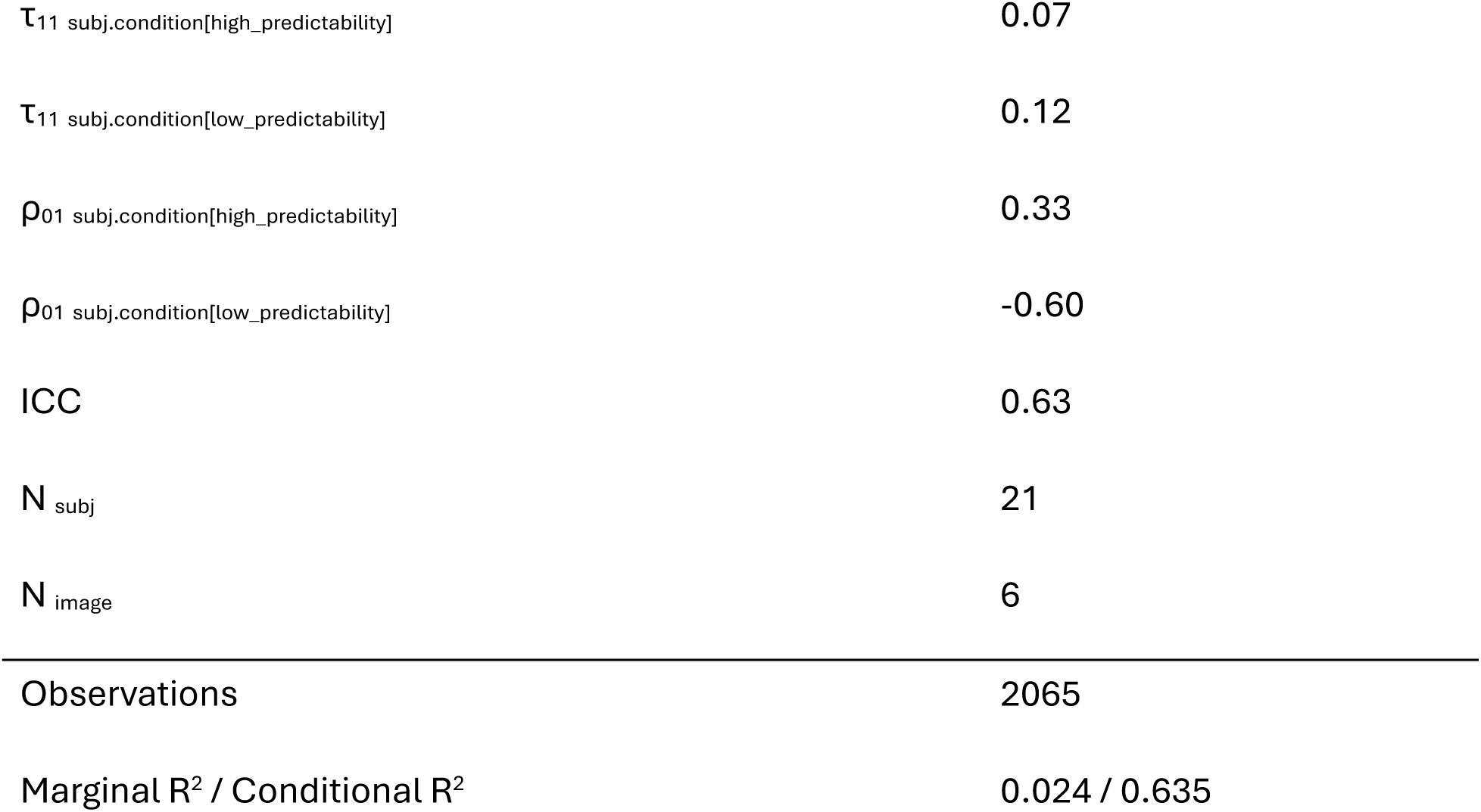

| *Predictors* | Pre-feedback theta power and performance |  |  |
| --- | --- | --- | --- |
|  | *Estimates* | *CI* | *p* |
| (Intercept) | -0.02 | -<br>0.35 – 0.31 | 0.91<br>1 |
| scaled_pre_theta | 0.01 | -<br>0.03 – 0.05 | 0.52<br>0 |
| condition[high_predictability] | 0.15 | 0.03 – 0.28 | **0.019** |
| condition[low_predictability] | -0.20 | -0.36 – -0.05 | **0.009** |
| roi[l_ant] | 0.01 | -<br>0.05 – 0.06 | 0.80<br>2 |
| roi[l_post] | 0.01 | -<br>0.05 – 0.06 | 0.84<br>9 |
| roi[midline] | -0.02 | -<br>0.08 – 0.04 | 0.54<br>4 |
| roi[r_ant] | 0.00 | -<br>0.05 – 0.0<br>6 | 0.87<br>0 |
| scaled_epoch | 0.05 | 0.02 – 0.0<br>8 | **0.00**<br>**1** |
| scaled_baseline_theta | 0.03 | -<br>0.01 – 0.0<br>7 | 0.10<br>8 |
| scaled_pre_theta:condition[high_predictability] | -0.00 | -<br>0.06 – 0.0<br>5 | 0.86<br>6 |
| scaled_pre_theta:condition[low_predictability] | -0.02 | -<br>0.08 – 0.0<br>3 | 0.45<br>7 |
| scaled_pre_theta:roi[l_ant] | 0.02 | -<br>0.04 – 0.0<br>8 | 0.48<br>2 |
| scaled_pre_theta:roi[l_post] | 0.00 | -<br>0.06 – 0.0<br>7 | 0.94<br>3 |
| scaled_pre_theta:roi[midline] | -0.01 | -<br>0.05 – 0.0<br>4 | 0.83<br>8 |
| scaled_pre_theta:roi[r_ant] | -0.00 | -<br>0.06 – 0.0<br>6 | 0.96<br>3 |
| condition[high_predictability]:roi[l_ant] | -0.00 | -<br>0.08 – 0.0<br>8 | 0.98<br>3 |
| condition[low_predictability]:roi[l_ant] | -0.00 | -<br>0.08 – 0.0<br>7 | 0.94<br>0 |
| condition[high_predictability]:roi[l_post] | -0.00 | -<br>0.08 – 0.0<br>8 | 0.96<br>3 |
| condition[low_predictability]:roi[l_post] | -0.00 | -<br>0.08 – 0.0<br>8 | 0.98<br>2 |
| condition[high_predictability]:roi[midline] | 0.00 | -<br>0.08 – 0.0<br>9 | 0.94<br>8 |
| condition[low_predictability]:roi[midline] | 0.01 | -<br>0.07 – 0.0<br>9 | 0.84<br>3 |
| condition[high_predictability]:roi[r_ant] | -0.00 | -<br>0.08 – 0.0<br>8 | 0.98<br>6 |
| condition[low_predictability]:roi[r_ant] | -0.00 | -<br>0.08 – 0.0<br>7 | 0.96<br>9 |
| scaled_pre_theta:condition[high_predictability]:roi[l_ant] | 0.01 | -<br>0.08 – 0.1<br>0 | 0.87<br>1 |
| scaled_pre_theta:condition[low_predictability]:roi[l_ant] | -0.01 | -<br>0.09 – 0.08 | 0.89<br>3 |
| scaled_pre_theta:condition[high_predictability]:roi[l_post] | -0.00 | -<br>0.10 – 0.09 | 0.92<br>4 |
| scaled_pre_theta:condition[low_predictability]:roi[l_post] | -0.00 | -<br>0.10 – 0.09 | 0.91<br>7 |
| scaled_pre_theta:condition[high_predictability]:roi[midline] | 0.00 | -<br>0.07 – 0.07 | 0.96<br>9 |
| scaled_pre_theta:condition[low_predictability]:roi[midline] | 0.00 | -<br>0.06 – 0.07 | 0.89<br>6 |
| scaled_pre_theta:condition[high_predictability]:roi[r_ant] | -0.01 | -<br>0.10 – 0.08 | 0.85<br>2 |
| scaled_pre_theta:condition[low_predictability]:roi[r_ant] | 0.01 | -<br>0.07 – 0.10 | 0.74<br>3 |

Random Effects
|  |  |
| --- | --- |
| $\sigma^2$ | 0.39 |
| $\tau_{00}$ subj | 0.56 |
| $\tau_{00}$ image | 0.01 |
| $\tau_{11}$ subj.condition[high_predictability] | 0.07 |
| $\tau_{11}$ subj.condition[low_predictability] | 0.12 |
| $\rho_{01}$ subj.condition[high_predictability] | 0.33 |
| $\rho_{01}$ subj.condition[low_predictability] | -0.60 |
| ICC | 0.63 |
| $N_{\text{subj}}$ | 21 |
| $N_{\text{image}}$ | 6 |
| Observations | 2065 |
| Marginal $R^2$ / Conditional $R^2$ | 0.024 / 0.635 |

## Appendix G: pre-feedback PSD plot

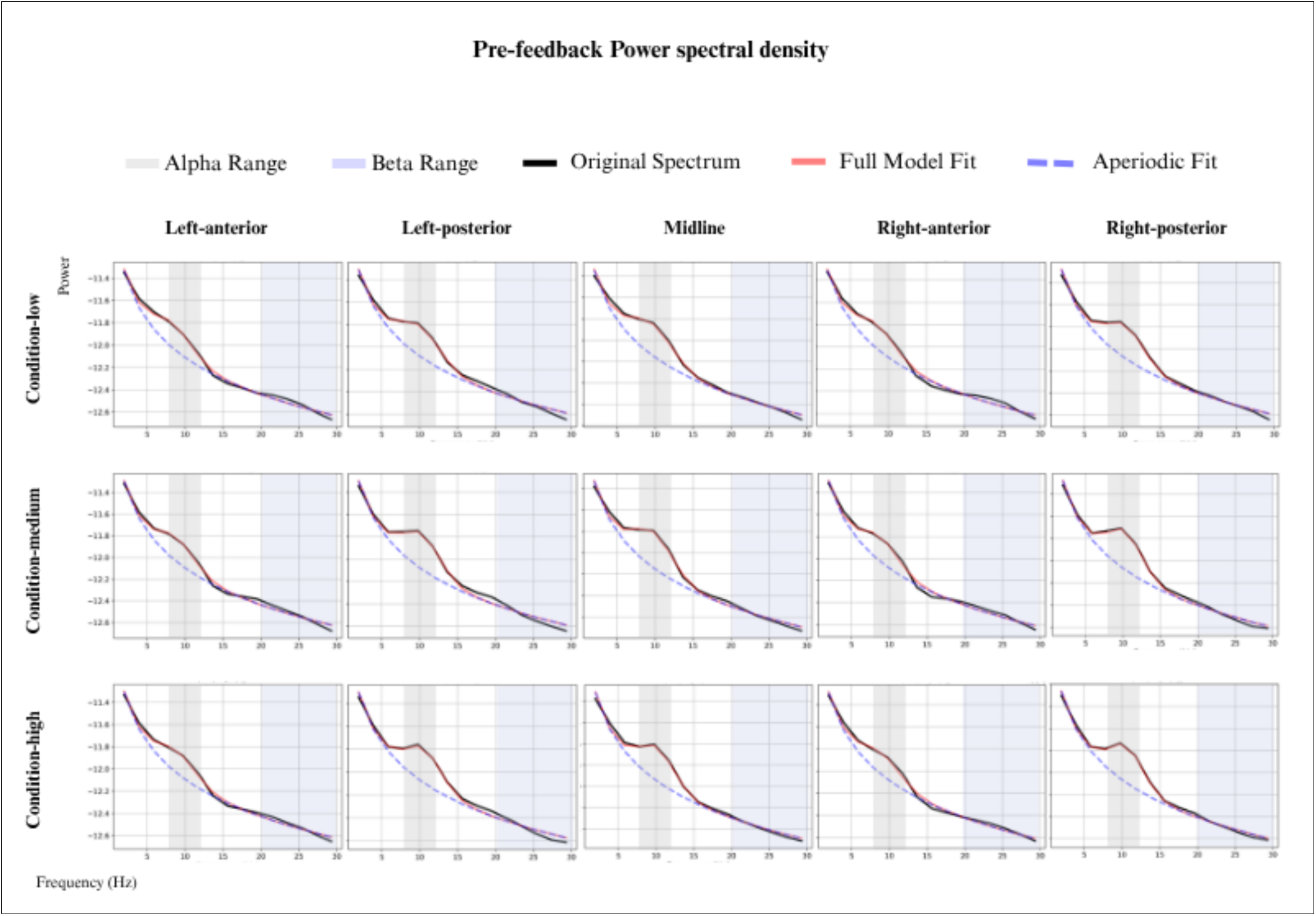

## Appendix H: feedback-related PSD plot

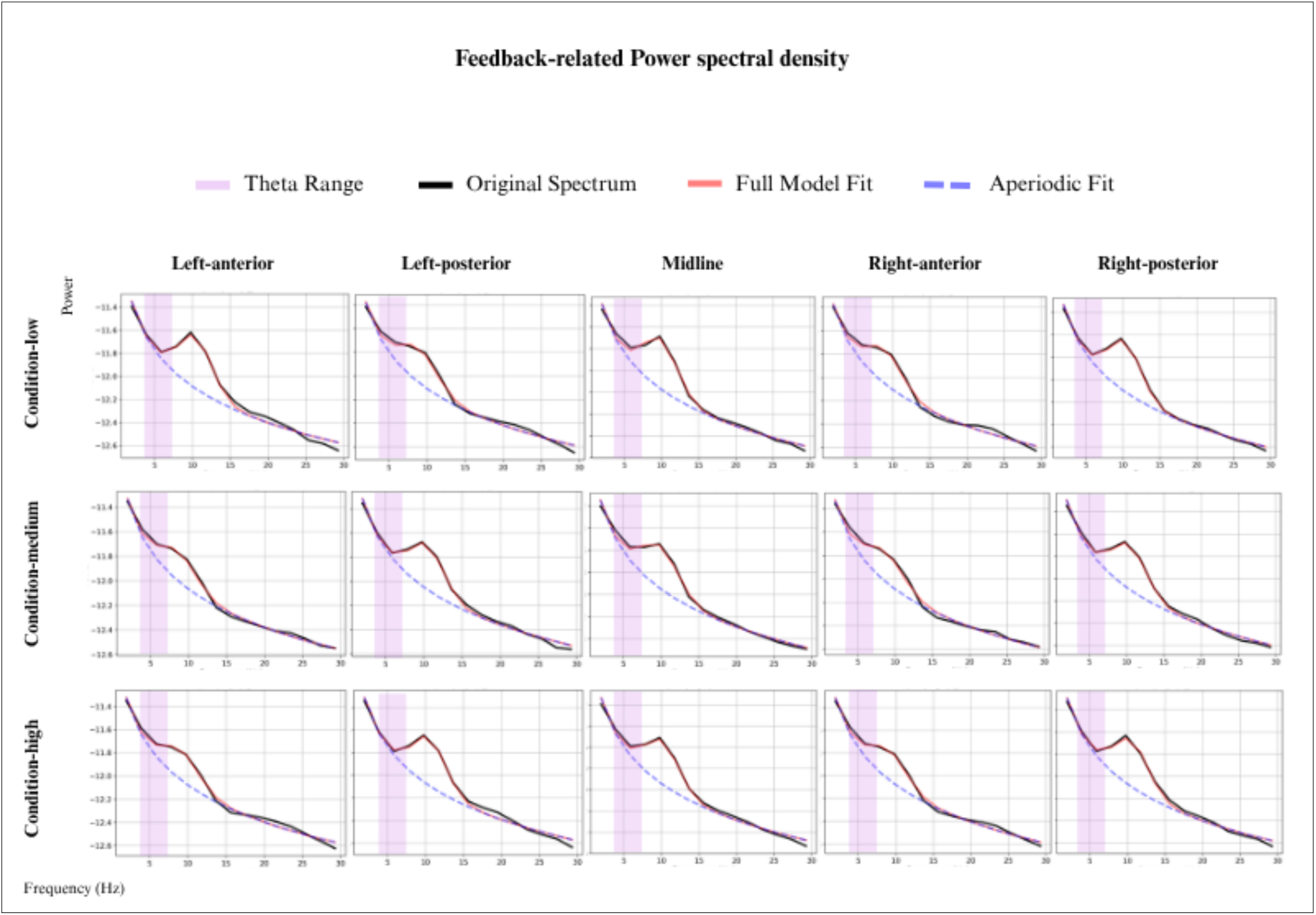

## CerdiT authorship contribution statement

**Maya Al Safadi:** Conceptualization, Data Curation, Formal analysis, Investigation, Project Administration, Visualization, Writing - Original Draft Preparation. **Alex Chatburn:** Conceptualization, Formal analysis, Supervision, Writing - Review & Editing. **Zachriah Cross:** Supervision, Writing - Review & Editing. **Shane Dawson:** Supervision, Writing - Review & Editing. **Ina Bornkessel-Schlesewsky:** Conceptualization, Formal Analysis, Project Administration, Supervision, Writing - Review & Editing.

## Declaration of completing interest

The authors declare that they have no known competing financial interests or personal relationships that could have appeared to influence the work reported in this paper.

## Data availability

Data reanalysis scripts are available upon request

## Notes

https://pmc.ncbi.nlm.nih.gov/articles/PMC10851313/

### Competing Interest Statement

The authors have declared no competing interest.

